# Global genomic diversity of the selfing nematode *Caenorhabditis tropicalis* correlates with geography

**DOI:** 10.64898/2026.04.05.716573

**Authors:** Bowen Wang, Nicolas D. Moya, Robyn E. Tanny, Michael E.G. Sauria, Lance M. O’Connor, Ayeh Khorshidian, Ryan McKeown, Lewis Stevens, Claire Buchanan, Timothy A. Crombie, Clayton Dilks, Kathryn S. Evans, Daniel E. Cook, Gaotian Zhang, Loraina A. Stinson, Nicole M. Roberto, Daehan Lee, Stefan Zdraljevic, Charlie Gosse, Clotilde Gimond, Mu-En Chen, Viet Dai Dang, John Wang, Asher D. Cutter, Matthew V. Rockman, Marie-Anne Félix, Christian Braendle, Erik C. Andersen

**Author notes:** **Corresponding author:** Erik C. Andersen Johns Hopkins University Department of Biology 3400 N. Charles St. Bascom UTL 383 Baltimore, MD 21218.

## Abstract

Self-fertilization reduces genetic diversity compared to outcrossing and hypothetically decreases the ability to adapt to diverse environments. Among *Caenorhabditis* nematodes, self-fertilization evolved three times independently in *Caenorhabditis elegans*, *Caenorhabditis briggsae*, and the more recently discovered *Caenorhabditis tropicalis*. To survey *C. tropicalis* genetic relatedness, the influence of geography and niche on species-wide variation, and the signatures of selection, we collected 785 wild strains, sequenced their genomes, and identified 622 distinct genotypes (isotypes). In contrast to *C. elegans* and *C. briggsae*, *C. tropicalis* relatedness shows substantial association with geography and no transcontinental selective sweeps or broadly sampled isotypes. Populations from the Hawaiian Islands or Taiwan harbor more genetic variation than populations from the Caribbean or Americas, suggesting a Pacific species origin similar to other members of the Elegans subclade. Punctuated genomic regions of extreme genetic variation pervade the genome. These hyper-divergent regions (HDRs) comprise less than 6% of the reference genome in any given strain despite harboring 73% of all variant sites and are enriched for genes likely involved in environmental adaptation. HDRs represent a shared genomic feature of self-fertilizing *Caenorhabditis* nematodes despite their independent evolutionary origins and suggest a mechanism to explain worldwide distributions despite low species-wide levels of genetic variation.

## Introduction

Mating system is a major driver of genetic diversity (Hartfield et al. 2017; Cutter 2019; Clo et al. 2025), with self-fertilization (selfing) increasing homozygosity, reducing effective population size, and decreasing the effectiveness of recombination, exacerbating the reduction of genetic diversity by linked selection (Pollak 1987; Charlesworth and Wright 2001; Wang et al. 2016; Teterina et al. 2023). Genetically homogeneous populations might compromise survival under environmental challenges because adaptive alleles are not present, leading to possible species extinction or an “evolutionary dead-end” (Takebayashi and Morrell 2001; Goldberg et al. 2010; Igic and Busch 2013). And yet, *Caenorhabditis elegans* genomes contain punctuated regions of extreme genetic variation, known as hyper-divergent regions (HDRs) (Lee et al. 2021). These regions might serve to maintain adaptive potential by enabling responses to diverse environmental conditions, despite the reduced variation associated with selfing (Andersen et al. 2012; Lee et al. 2021). However, it is unknown how general these global patterns for *C. elegans* apply to other selfing species (Moya et al. 2024).

Selfing evolved independently three times in the *Caenorhabditis* genus, including the widely studied model organism *C. elegans*, the comparative model species *Caenorhabditis briggsae*, and the more recently discovered species *Caenorhabditis tropicalis* (Kiontke et al. 2011; Ellis 2017). Phylogeographically, globally distributed *C. briggsae* lacks the chromosome-scale selective sweeps seen in *C. elegans* that generate highly related strains across continents (Moya et al. 2025). Instead, *C. briggsae* strains can be classified into twelve genetically divergent relatedness groups, comprising geographically restricted groups (Moya et al. 2025). Both *C. elegans* and *C. briggsae* show similar genome-wide distribution of natural variation, with lower polymorphism than related outbreeding species and widespread HDRs punctuated across the high-recombination chromosome arms (Lee et al. 2021; Teterina et al. 2023; Moya et al. 2025). *C. tropicalis* has a more restricted and primarily tropical global distribution where it is frequently isolated from decaying leaves, flowers, and fruits, and is commonly maintained at higher culture temperatures than the other two selfing species (Kiontke et al. 2011; Félix et al. 2013; Gimond et al. 2013; Félix et al. 2014; Frézal and Félix 2015; Poullet et al. 2015; Crombie et al. 2019; Crombie et al. 2022; Crombie et al. 2024). Despite limited strains characterized, analysis of wild *C. tropicalis* genomic variation indicates a strong impact of oceanic barriers to dispersal, unlike *C. elegans* and *C. briggsae*, though *C. tropicalis* genomes appear to also harbor HDRs (Noble et al. 2021).

In this study, we sequenced the whole genomes of 785 wild *C. tropicalis* strains collected from localities around the globe by the *Caenorhabditis* research community, massively expanding the scope of the previous *C. tropicalis* population studies (Gimond et al. 2013; Noble et al. 2021). We identified single-nucleotide and insertion-deletion variants to reveal extensive geographic and genetic differentiation that strongly correlates with geography. Moreover, the Hawaiian Islands and Taiwan harbor the highest levels of genetic diversity. Across the genome, *C. tropicalis* exhibits an even greater concentration of genomic diversity in HDRs compared to *C. elegans* and *C. briggsae*. In these two species, 20% and 31% of alternative SNV alleles fall within HDRs, whereas in *C. tropicalis*, less than 6% of the reference genome within any strain harbors 73% of all alternative SNVs. Similar to *C. elegans* and *C. briggsae*, *C. tropicalis* HDRs also show substantial gene content variation and are enriched for genes involved in environmental responses. Together, our findings suggest that HDRs provide a shared genomic solution that enables independently evolved selfing species to maintain adaptive potential and exhibit global distributions despite low genome-wide diversity.

## Results

### Global genome-wide variation is strongly correlated with geography

Among selfing *Caenorhabditis* species, *C. tropicalis* has the most restricted global distribution, with samples collected from tropical and subtropical regions worldwide (Fig. S1). Extensive collections are primarily from Taiwan, the Caribbean, French Guiana (Félix et al. 2013; Ferrari et al. 2017), Hawaii (Crombie et al. 2019; Crombie et al. 2022), Panama and Costa Rica (Sloat et al. 2022), and the island of Pohnpei in Micronesia (Rockman et al. 2025) (Fig. 1a-b, Table S1). Collectively, these widely distributed sampling sites indicate extensive colonization of *C. tropicalis* across tropical regions of the Pacific Rim and on tropical islands surrounding the African continent (Fig. S1). From this circumglobal collection of strains, we generated whole-genome sequence data for 785 wild *C. tropicalis* strains (see Methods, Table S2). Because *C. tropicalis* is highly selfing, wild strains are naturally inbred and essentially homozygous across the genome with clonal expansions leading to nearly identical genomes in independently sampled strains. To avoid analyzing redundant genome sequences, we classified the 785 strains into 622 distinct genome-wide haplotypes (referred to as isotypes), each represented by an isotype reference strain (Fig. S2, Table S1, Table S2). Most isotypes comprise a single strain (n=521) or multiple strains sampled within one kilometer (km) of one another (n=86). The other two selfing *Caenorhabditis* species also have most isotypes either comprising a single strain or multiple strains isolated within one km of one another (*C. elegans*: 638/684, *C. briggsae*: 630/715, CaeNDR release 20250626) (Crombie et al. 2024; Moya et al. 2025). The remaining 15 isotypes of *C. tropicalis* comprise strains isolated from the same geographic region with pairwise distances ranging from just over one km to 19 km (Fig. S2c-d).

**Figure 1:**
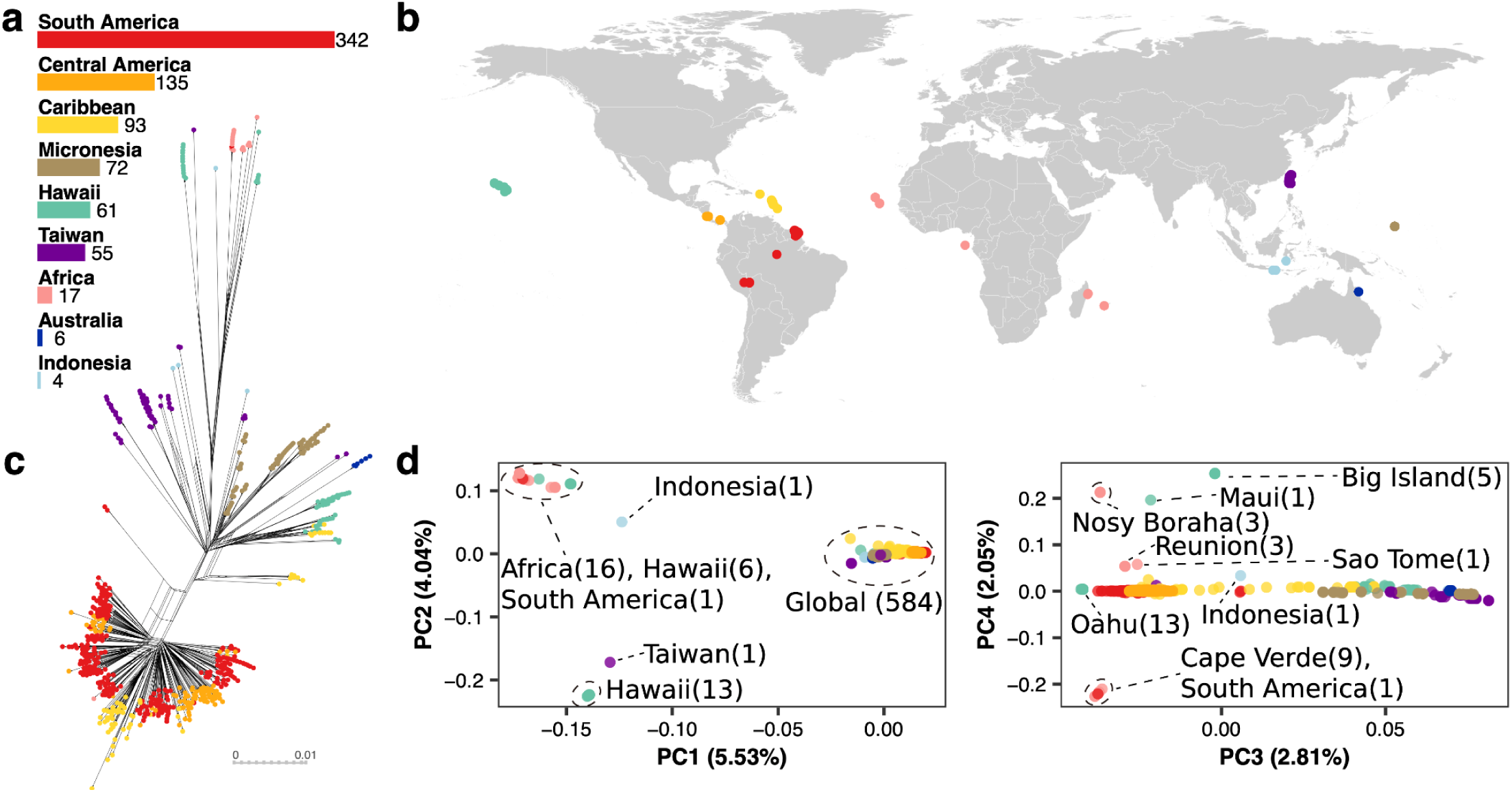
Global distribution of 622 genetically distinct strains. a, Sampling frequency of 785 strains in each geographic region. **b,** Global sampling map of 785 strains. **c,** Neighbor-joining network of 622 *C. tropicalis* isotype reference strains. The tree was generated from LD-pruned variants with r^2^ value less than 0.9 (see Methods). **d,** Principal Component Analysis (PCA) of global *C. tropicalis* isotype reference strains. Colors in each panel represent the sampling geographic region of each strain. The ellipses highlight differentiated isotypes. The numbers in parentheses represent the count of isotypes within each cluster.

Across the genomes of the 622 isotype reference strains, we identified 1,696,592 single-nucleotide variants (SNVs) and 336,230 short (<50 base pairs (bp)) insertion-deletion (indel) variants (see Data Availability, Materials and Methods). Even though the *C. tropicalis* genome is approximately 20% smaller than both the *C. elegans* and *C. briggsae* genomes, this level of genome-wide variation is less than half that observed in *C. elegans* (∼3.5 million SNVs and 0.59 million short indel variants across 684 isotypes, CaeNDR release 20250625) and less than a third seen for *C. briggsae* (∼5.65 million SNVs and 0.88 million short indel variants across 715 isotypes) (Crombie et al. 2024; Moya et al. 2025). Genome-wide linkage disequilibrium (LD) patterns were broadly similar across species, with *C. tropicalis* showing a slightly sharper decay (Fig. S3). To investigate population structure, we constructed a distance-based network using genome-wide variants (Fig. 1c). The overall structure of the network closely reflects the geography of the samples, with most isotypes from the Americas forming a major cluster, and other isotypes from varied geographic regions diverging from this cluster and forming more differentiated branches (Fig. 1c). We then performed principal component analysis (PCA) using biallelic SNVs after pruning variants in strong LD (r^2^ > 0.9, see Methods) (Fig. 1d). Nearly all isotypes (n=584) fall within a single cluster in PC1 and PC2 space, with the remaining isotypes derived from restricted geographic regions forming several discrete clusters (Fig. 1d). These discrete clusters include a cluster of 13 isotypes from Oahu, a Hawaiian Island; a cluster of 16 isotypes from Africa, one isotype from South America, and six isotypes from other Hawaiian Islands (five from the Big Island, one from Maui); a cluster formed by a single isotype from Taiwan; and another cluster formed by a single isotype from Indonesia (Fig. 1d, Fig. S4, Fig. S5, Table S3). PC3 and PC4 described additional independent axes of genomic variation and provided higher-resolution structure for several differentiated clusters defined by PC1 and PC2, including geographically distinct subclusters (Fig. 1d, Fig. S5, Table S3). Chromosome-specific PCs largely recapitulate the patterns observed genome-wide (Fig. S6). These results framed our further characterization of genetic relatedness among global *C. tropicalis* isotypes and examination of possible associations between geography and patterns of genetic variation.

### Taiwan and the Hawaiian Islands are *C. tropicalis* genetic diversity hotspots

To further investigate population structure in *C. tropicalis*, we calculated mean pairwise genetic similarity among isotypes based on the proportion of identical alleles across genome-wide SNVs (see Methods). The 19 relatedness groups were then manually defined based on major blocks in the clustered heatmap (Fig. 2, Table S1). Five of these 19 groups represent the differentiated PC clusters. The 14 remaining groups represent subdivision of isotypes from the Global PC cluster in the first two PCs. The 19 relatedness groups were named according to their primary geographic origins (Fig. 2, Table S1). These included a large Latin America and Caribbean (LAC) group, six Taiwan groups (Tw1-Tw6), two Micronesia groups (Mic1 and Mic2), a Hawaii-Caribbean (HC) group, and several smaller geographically restricted groups. The 19 relatedness groups corresponded to a discrete separation in pairwise genetic similarity, with the mean within-group genetic similarity exceeding 93% and the mean between-group similarity consistently below 92% (Fig. S7). Admixture analysis broadly recapitulated the relatedness groups (Fig. S8-S10). Using cross-validation, we identified the number of populations (*K*=28) with consistent ancestry patterns observed across replicate runs (Fig. S8). These results provided an independent ancestry-based perspective that was consistent with relatedness groups defined by genome-wide genetic similarity (Fig. S8-S11). The association between genetic similarity and geography was further supported by a significant negative correlation between genetic similarity and geographic distance both globally and within individual relatedness groups, suggesting isolation by distance (Fig. S12-S13).

**Figure 2:**
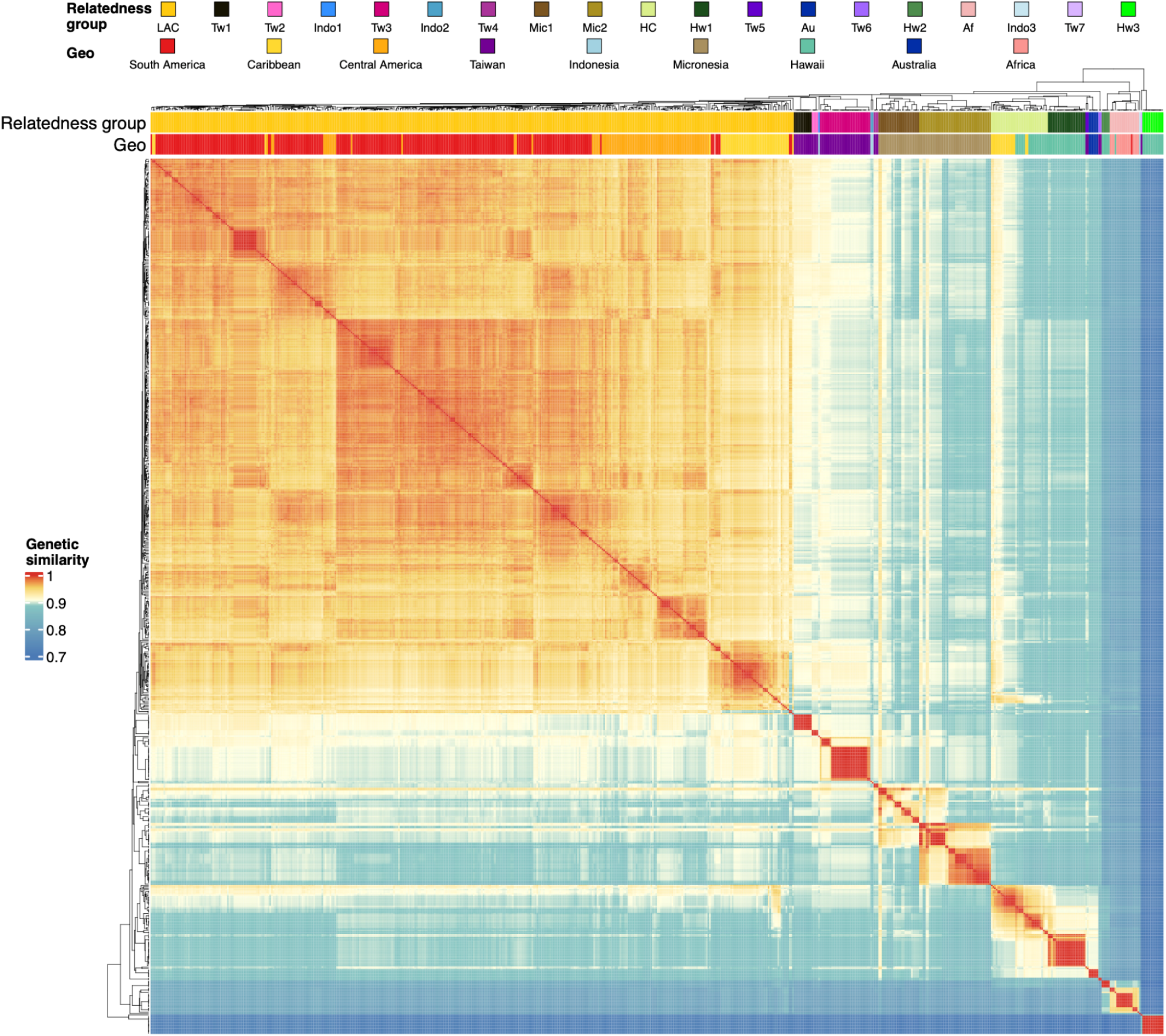
Pairwise genome-wide genetic similarity defines *C. tropicalis* relatedness groups. Heatmap showing pairwise genetic similarity between all 622 isotype reference strains. Pairwise genetic similarity was estimated by the proportion of identical alleles across all identified SNVs among the 622 reference strains. The genetic similarity color gradient has breaks specified at the 25th, 50th, and 75th percentile of the distribution of all pairwise estimates. The top bands show the geographic region of the isolation site of each isotype representative strain and the assigned relatedness group.

Given that the majority of *C. tropicalis* relatedness groups are largely confined to a limited subset of the geographic regions (from current sampling locations), we conducted more detailed investigations of the group distribution patterns within each geographic region. We found that Taiwan and the Hawaiian Islands harbor the largest numbers of distinct relatedness groups in our sample (Taiwan, seven groups; the Hawaiian Islands, five groups), which indicates the coexistence of multiple genetically divergent groups within these regions. This inference was further supported by regional comparisons of mean nucleotide diversity (π, the average pairwise sequence divergence) and Watterson’s θ (θ_W_, based on the number of segregating sites scaled by sample size) (Table S4), where Taiwan and the Hawaiian Islands exhibited the highest π and θ_W_ values among all geographic regions with more than 10 sampled isotypes. Notably, Indonesia harbors three distinct relatedness groups from only four isotypes, suggesting that additional sampling could reveal it to also be a hotspot of *C. tropicalis* diversity. Overall, despite globally low variation for *C. tropicalis*, the Pacific islands associated with Taiwan and the Hawaiian Islands represent major diversity hotspots, with an additional potential third hotspot in Indonesia. However, we note that geographic regions differ in spatial scale and sampling intensity, which might contribute to the observed distribution of diversity.

Genetically differentiated relatedness groups show contrasting geographic distributions across regions. Taiwan and the Hawaiian Islands are composed of isotypes that were sampled within restricted geographic ranges (maximum distances not exceeding 27.6 km). By contrast, several other relatedness groups show much broader spatial distributions, with isotypes spanning long distances within the same island (Tw1, maximum geographic distance up to 273.6 km), multiple islands within a region (Hw1), or even distinct geographic regions (Hawaii-Caribbean) (Fig. S14-S17). In other regions, one or two broadly distributed groups dominate and show little internal geographic structure (Fig. S18-S24).Taiwan contains the largest number of relatedness groups, including one widely distributed group (Tw1) and several geographically restricted ones (maximum distances between isotypes within the same group not exceeding 27.6 km) (Fig. S14). Overall genetic variation in Taiwan correlates with certain environmental gradients based on PCA of Taiwanese isotypes only. For example, PC1 positively correlates with rainfall and negatively correlates with elevation (Fig. S15). However, comparisons among relatedness groups using environmental variables extracted from public geographic information system (GIS) datasets did not reveal clear differences among groups. The limitations of group size might constrain our ability to draw inferences consistent with local adaptation (Fig. S15). The Hawaiian Islands are the second most diverse region (Fig. S16). Comparisons among Hawaiian relatedness groups suggest association with environmental differences in at least one geographically restricted group (Hw2), albeit with strong inference of local adaptation similarly constrained by group sizes (Fig. S17). Most other regions show a more limited relatedness group composition. The distribution of relatedness groups within the Americas and Micronesia does not correspond to major sampling areas (Fig. S18-S24). Notably, because sampling in Micronesia is limited to 72 isotypes from one island, the observed relatedness group distribution likely reflects island-specific rather than regional structure. In addition, other under-sampled regions such as Indonesia and oceanic African islands, as well as Australia, where *C. tropicalis* has been infrequently observed, are suggestive of island-specific clustering as well as shared groups across islands or regions (Fig. S254-S276). However, the current number of available strains restricts strong conclusions. Overall, strong relatedness group differentiation and potential local adaptation are most conspicuous in Taiwan and the Hawaiian Islands, whereas other geographic regions are dominated by one or two relatedness groups whose distributions do not correspond to fine-scale geography.

### Genome-wide genetic diversity is largely confined to punctuated hyper-divergent regions

We next assessed the distribution of genetic diversity along chromosomes throughout the genome. Consistent with an emergent pattern for genomes of many *Caenorhabditis* species, we found that chromosome arms harbor approximately twice as much variation as chromosome centers (Fig. 3a, Table S5; 2.00-fold higher mean π and a 1.85-fold higher mean θ_W_ in autosomal arm domains) relative to center domains (Thomas et al. 2015; Crombie et al. 2019; Noble et al. 2021; Teterina et al. 2023; Moya et al. 2025). This intra-chromosomal heterogeneity in population genetic diversity is exacerbated in *C. tropicalis* by the extensive LD associated with selfing, which strengthens the effects of linked selection in low-recombination chromosome centers (Cutter and Payseur 2013; Cutter 2019). By contrast to X chromosome patterns in other selfing *Caenorhabditis* nematodes, *C. tropicalis* exhibits markedly elevated diversity on the X chromosome arms, with mean π and θ_W_ 2.57- and 1.55-fold higher, respectively, than the center (Fig. 3a, Table S5).

**Figure 3:**
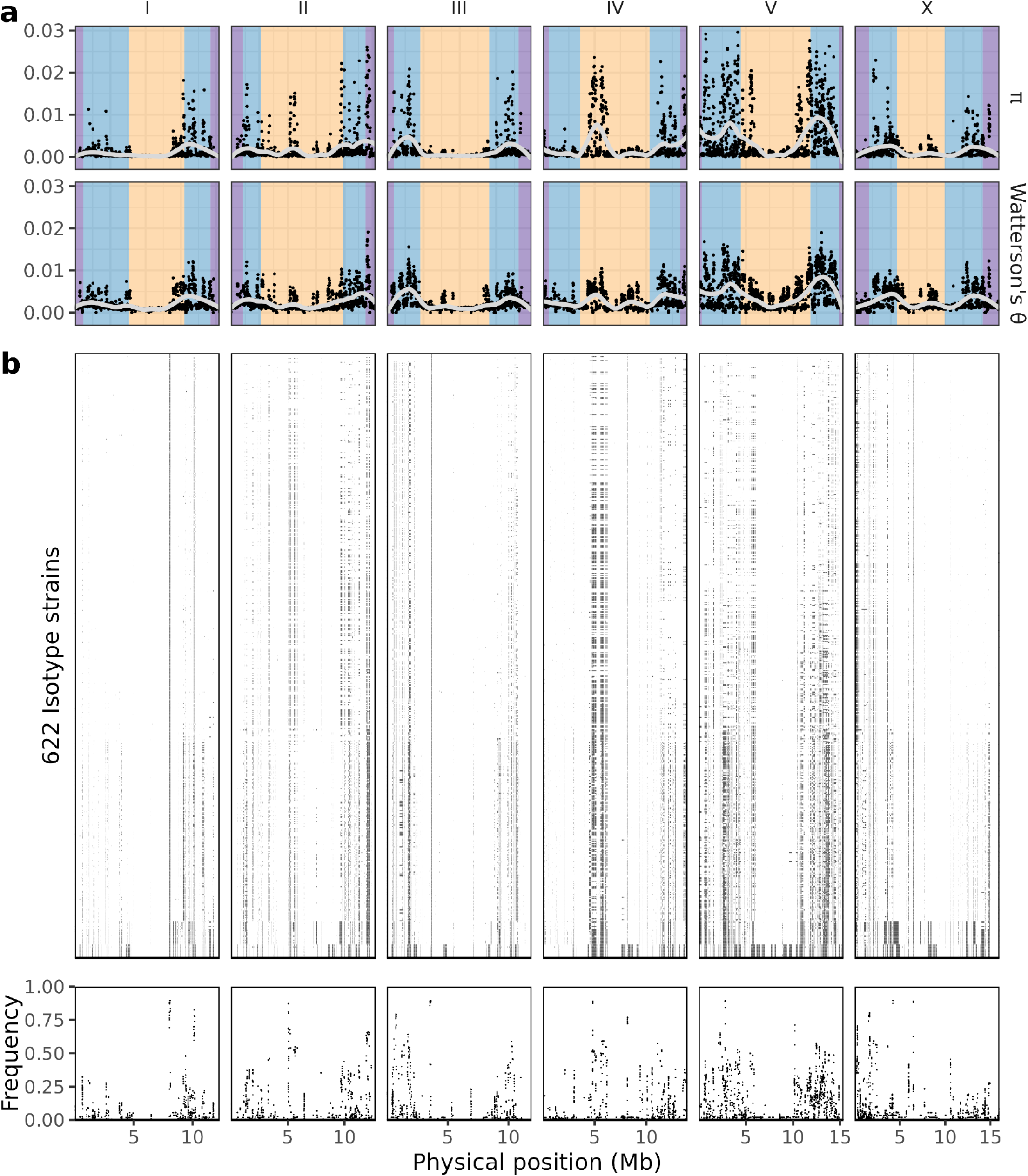
Diversity statistics and hyper-divergent regions across global *C. tropicalis* isotypes. a, Scatter plots show the nucleotide diversity (π) and Watterson’s θ estimates along physical genome positions across the six chromosomes. Each point represents an estimate of nucleotide diversity (π or θ_W_) across a 10 kb window. Grey lines are weighted LOESS fits where each local regression used 30% of the data. Colored backgrounds in nucleotide diversity panels delineate the chromosomal domain boundaries (purple, tip; blue, arm; yellow, center). **b,** In the top panel, each row represents one of 622 isotype reference strains, with HDRs in that strain relative to the NIC58 reference marked as black boxes across the genome. The frequency at which 1 kb bins are classified as hyper-divergent is shown in the bottom panel.

As observed in a much smaller dataset (Noble et al. 2021), most genetic variation in *C. tropicalis* is concentrated in a set of highly punctuated, extremely diverse genomic regions referred to as hyper-divergent regions (HDRs), a pattern previously characterized in *C. elegans* and *C. briggsae* (Lee et al. 2021; Moya et al. 2025). We applied methods established for *C. elegans* to identify HDRs using a single reference genome (NIC58) with modified thresholds optimized for *C. tropicalis* (see Methods). Across the 622 isotypes, we identified 689 non-overlapping HDRs, with an average length of 45 kb (range: 5 kb to 774 kb) (Fig. 3b, Table S6). Collectively, these HDRs span approximately 38% of the reference genome. Hyper-divergent haplotypes span less than 6% of the reference genome within any given isotype and yet harbor 73% of alternative SNV alleles on average (Fig. S28, Table S7). In comparison with *C. elegans* and *C. briggsae*, *C. tropicalis* genetic variation appears to be more concentrated in HDRs as just 20% and 31% of all alternative SNV alleles localize to HDRs in these other two species, respectively (Lee et al. 2021; Moya et al. 2025). Strains from the least genetically similar relatedness groups in *C. tropicalis* (*e.g*., Hw3, Tw7), however, have large proportions (up to 22%) of their genomes inferred to be hyper-divergent (Fig. S28, Table. S7). These elevated proportions likely reflect the greater divergence of these relatedness groups from the reference strain, such that certain hyper-divergent haplotypes are private and fixed within relatedness groups most diverged from the reference strain (Fig. S29-30). However, these private hyper-divergent haplotypes remain punctuated, clearly distinguishable from a genetic background that is substantially less variable than the HDRs themselves (Fig. S30). This pattern of punctuated fixed differences between divergent relatedness groups in *C. tropicalis* starkly differs from *C. briggsae*, where the most divergent relatedness groups displayed genome-wide elevated background variation that masked punctuated HDRs and required an approach that uses individual reference genomes for each relatedness group to identify HDRs (Fig. S31).

Genomic regions outside of HDRs show lower or similar estimates of absolute divergence (D_xy_) among relatedness groups relative to HDRs, but high-divergence windows are largely restricted to HDRs (Fig. S32). Although both D_xy_ and π are lower outside of HDRs, relatedness groups structure inferred from non-HDR regions largely recapitulates the genome-wide pattern (Fig. S33-S34). Additionally, the lower level of divergence is insufficient to disrupt the arm-center contrast outside of HDRs, a pattern shaped by recombination and linked selection across chromosome domains. Autosomal arms still harbor higher diversity than centers (1.97- and 1.61-fold for π and θ_W_), and the arm-center contrast on the X chromosome becomes more modest (1.47- and 1.13-fold for π and θ_W_) (Fig. S35, Table S8). The substantial sequence similarity outside of HDRs between *C. tropicalis* strains from genetically distant relatedness groups (mean outside of HDRs D_xy_ between LAC and Hw3 = 0.17) suggests a more recent separation of *C. tropicalis* groups than *C. briggsae* groups (Moya et al. 2025) (mean outside of HDRs D_xy_ between Tropical and KD = 0.40) and/or they evolved from a less diverse ancestral population (Table S9).

### Selfing *Caenorhabditis* species share similar genes within hyper-divergent regions

Some gene families are enriched in HDRs of both *C. elegans* and *C. briggsae*, including genes involved in environmental sensing and pathogen responses, such as C-type lectins, G-protein coupled receptors, nuclear hormone receptors, and E3 ubiquitin ligases (*e.g.*, F-box genes) (Lee et al. 2021; Moya et al. 2025). To characterize the functional role of hyper-divergent genes in *C. tropicalis*, we annotated functional protein domains and their associated gene ontology (GO) identifiers using InterProScan (Jones et al. 2014) (see Methods). This analysis identified functional protein domain annotations for approximately 71% (14,260 / 20,149) and GO identifiers for approximately 55% (10,985 / 20,149) of all NIC58 reference genes (Table S10). In total, 71.41% (14,388 / 20,149) of genes have at least one protein domain or GO annotation. To account for our expectation that chromosomal arm domains likely contain different gene classes and fewer essential genes than chromosomal centers like in *C. elegans* (Kamath et al. 2003), we performed an enrichment test for genes inside or outside of HDRs for arm domain regions where HDRs are primarily found (Fig. 4). We found that genes in HDRs on chromosomal arms are significantly enriched for functional protein domains associated with environmental and pathogenic responses (Fig. 4a, Table S11). Among enriched GO terms for the same gene sets, we found innate immune response genes like C-type lectins (115/144 genes, *e.g.*, orthologs of *C. elegans clec-59*, *clec-101*) (Fig. 4b). Additionally, we found enrichment for G protein-coupled peptide receptor activity (55/72 genes, Fig. 4c) and xenobiotic metabolism (18/21 genes, Fig. 4b), where many of the genes in xenobiotic metabolism (15/18) were identified as orthologs of *C. elegans* cytochrome P450 families *cyp-33*, *cyp-34*, and *cyp-35*. By contrast, enrichment tests for genes in non-HDR chromosomal arm regions mainly identified structural or housekeeping domains (*e.g.*, WD40/YVTN repeat-like-containing domain superfamily, 19/86 genes; and P-loop containing nucleoside triphosphate hydrolase, 53/230 genes), none of which ranked among the top 20 enriched terms. Additionally, we found enrichment of a few, non-environment related gene ontology terms in these non-HDR arm regions (biological process: translation; molecular function: structural constituent of ribosome, ATP binding). These results suggest that *C. tropicalis* HDRs, like *C. elegans* and *C. briggsae* HDRs, are enriched for environmental response genes likely associated with local adaptation to heterogeneous ecological factors.

**Figure 4:**
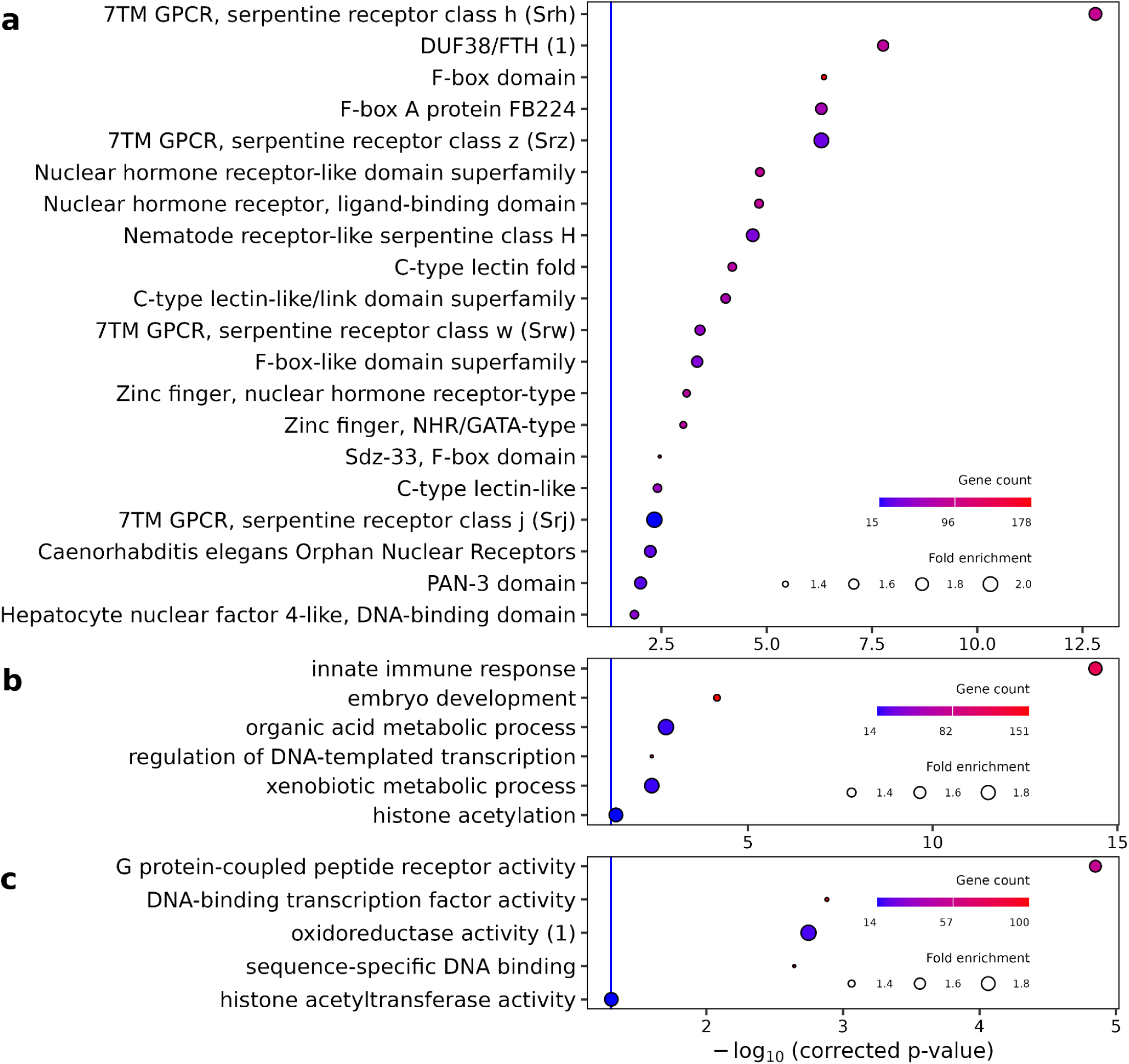
***C. tropicalis* hyper-divergent regions are enriched for environmental response genes.** (**a**) Significantly enriched InterProScan functional protein domains in genes found in hyper-divergent region chromosome arms. Each protein domain annotation on the y-axis showed a false discovery rate-corrected p-value (q-value) of lower than 0.05 (negative log-scaled adjusted p-values shown on the x-axis) biological process Significant biological process (**b**) and molecular function (**c**) gene ontology terms from gene set enrichment analysis. Each term present in the y-axis showed a Benjamini-Hochberg adjusted p-value lower than 0.05 (negative log-scaled adjusted p-values shown in the x-axis). In each row, the circle varies in size based on the fold-enrichment associated with that term, and in shading based on the number of genes in chromosome arm hyper-divergent regions associated with that term. “DUF38/FTH (1)” refers to the InterProScan description “Domain of unknown function DUF38/FTH, Caenorhabditis species”. “oxidoreductase activity (1)” refers to the gene set term “oxidoreductase activity, acting on paired donors, with incorporation or reduction of molecular oxygen, reduced flavin or flavoprotein as one donor, and incorporation of one atom of oxygen”. All enrichment tests shown use arm domain genes as a background gene set.

## Discussion

This study represents the largest global survey of *C. tropicalis* to date. We assigned the collection of 785 sampled *C. tropicalis* isolates to 622 *C. tropicalis* genomic isotypes and 19 relatedness groups that capture the major axes of genetic variation. Across the species range, some geographic regions contain representatives of many relatedness groups and are hotspots of genetic diversity (Taiwan and the Hawaiian Islands, in particular), consistent with a geographic origin of *C. tropicalis* around the Pacific, provided that high genetic diversity reflects long-term evolutionary history and the maintenance of ancestral variation near the species origin (Li and Stephan 2006; Nielsen et al. 2017; Peter et al. 2018). During range expansion, genetic diversity generally decreases because of serial founder effects, a pattern also observed in human global dispersal (Henn et al. 2012). However, the geographic density of selfing *Caenorhabditis* sampling remains sparse throughout the world, particularly because of under-sampled regions like southeast Asia and Pacific islands, which might harbor substantial diversity that is currently undiscovered. Expanding sampling in these key regions will be essential for reconstructing the dispersal history of this species. By contrast to Pacific sampling localities, nearly all isotypes from the Americas region belong to a single relatedness group (LAC) suggestive of a relatively recent colonization and spread of *C. tropicalis* across the Caribbean, Central America, and tropical South America. The lower genetic diversity in the Americas might also be shaped by environmental selective pressures. Future work integrating ecological variables, environmental gradients, with functional genomic data from the Americas will help clarify how genetic differentiation and ecological adaptation might interact within this group. Notably, the LAC group is the most similar to Taiwanese groups, which suggests that these groups might share a more recent common ancestor, suggesting a potential migration event from the Pacific to the Americas. In addition, the HC relatedness group is distributed across both the Hawaiian Islands and the Caribbean, which is consistent with the proposed possible dispersal history from the Pacific to the Americas.

Balancing selection has been considered the most likely force that enables the long-term maintenance of HDRs in selfing *Caenorhabditis* species (Lee et al. 2021; Moya et al. 2025). However, recent studies on toxin-antidote (TA) systems (including the maternal-effect TA elements often referred to as Medea elements) have provided additional insight into the origin of HDRs. TA systems are hypothesized to express a toxin that causes embryonic lethality in offspring that fail to inherit the corresponding antidote (Ben-David et al. 2021; Lee et al. 2021; Noble et al. 2021; Velazco-Cruz and Ross 2022; Widen et al. 2023; Tikanova et al. 2025). Multiple TA systems have been identified in all three selfing *Caenorhabditis* species (Seidel et al. 2008; Ben-David et al. 2017; Ben-David et al. 2021; Lee et al. 2021; Widen et al. 2023; Tikanova et al. 2025), often found within structurally complex and deeply diverged haplotype regions (hereafter referred to as TA regions), which are also HDRs (Lee et al. 2021; Rockman 2025). Consequently, we compared previously described *C. tropicalis* TA regions with HDRs and found extensive overlap (Fig. S36, Table S12). Recent theoretical work further clarifies why HDRs and TA elements might overlap (Rockman 2025). TA gene-drive systems reduce gene flow between incompatible haplotypes, thus maintaining those distinct haplotypes and allowing them to further accumulate divergence. In partially selfing systems, TA elements do not operate through classical within-population balancing selection (*e.g*., heterosis, temporally-varying selection, negative frequency-dependent selection) but instead exhibit positive frequency dependence such that a common TA haplotype drives rare haplotypes toward local extinction allowing distinct TA haplotypes to monopolize local populations while maintaining different haplotypes across a metapopulation (Rockman 2025). Selfing amplifies this drive effect by reducing heterozygosity to facilitate the long-term maintenance of large, ancient haplotype blocks fixed in separate populations. This mechanism provides a potential non-adaptive explanation for the persistence of HDRs. In addition, a recent study revealed that repeated tandem duplications of an ancestral locus containing the essential gene *fars-3* and an F-box gene can give rise to multiple independent toxin-antidote systems in *C. tropicalis* (Tikanova et al. 2025). Moreover, a TA system involving F-box genes in *C. nigoni* contributes to inter-species hybrid incompatibility with *C. briggsae* (Xie et al. 2026), pointing to how such evolution can contribute to reproductive isolation and speciation. This work collectively helps to explain how *C. tropicalis* could maintain so many distinct HDRs while also being consistent with demonstrations of outbreeding depression and multiple incompatible haplotypes (Gimond et al. 2013; Tikanova et al. 2025). Given that HDRs harbor extensive structural variation and large segments absent from or poorly represented in the reference genome, future pangenome-based haplotype reconstruction will be essential to resolve the precise boundaries and structural diversity of HDR and TA regions across relatedness groups.

Pangenome-based analyses and haplotype reconstruction for *C. tropicalis* would offer additional context for deciphering demographic history as well as the structural features of HDRs and their contributions to genetic diversity. Compared to *C. briggsae*, *C. tropicalis* relatedness groups remain relatively similar to one another outside of HDRs, which suggests that genome-wide divergence among groups is limited (Moya et al. 2025). Consistent with this observation, *C. tropicalis* HDRs are sharply punctuated and discrete, which suggests a more recent separation among relatedness groups and/or the origin of distinct relatedness groups from a less diverse ancestral population. Clarifying these alternatives will require pangenome-based analyses to better characterize variations within HDRs. We also found *C. tropicalis* exhibits a more pronounced arm-center contrast of X-linked genetic diversity than is observed in either *C. elegans* or *C. briggsae* (Cutter and Payseur 2013; Cutter 2019), with depressed genetic diversity in center regions not restricted to autosomes for *C. tropicalis.* Future pangenome-based analyses and haplotype reconstruction will enable a more comprehensive characterization of potential structural features of these HDRs in autosomal centers.

The global *C. tropicalis* population resource generated in this study also provides an extensive foundation for diverse future investigations, including genome-wide association analyses (GWAS), experimental molecular genetic interrogation of *C. tropicalis* as a satellite model to *C. elegans*, and comparative biology more broadly within *Caenorhabditis*. Given the highly structured relatedness groups of global *C. tropicalis*, future GWAS can be conducted within specific relatedness groups to reduce false positives caused by population stratification (Evans et al. 2017). In addition, the extensive haplotype diversity within HDRs make pangenome-based mapping particularly important for accurately identifying functional variants embedded within highly diverged haplotypes. Combined with the extensive resources also available for natural strains of *C. elegans* and *C. briggsae* (Lee et al. 2021; Crombie et al. 2024; Moya et al. 2025), including whole-genome sequences and diverse phenotypic measurements, *Caenorhabditis* now provides valuable opportunities for comparative mappings across species. By integrating population genomic data from all three selfing *Caenorhabditis* species, future comparative analyses could facilitate the identification of shared evolutionary processes (Yin et al. 2018), refining our understanding of general principles in the evolution of selfing systems. With continued advances in long-read sequencing, pangenome construction, and nematode high-throughput phenotyping, joint mapping across these three species is expected to reveal deeper connections between genome structure and phenotypic variation in selfing *Caenorhabditis*.

## Materials and Methods

### Strains

Nematodes were reared at room temperature (22-25°C) using *Escherichia coli* bacteria (strain OP50) grown on modified nematode growth medium (NGMA) (Andersen et al. 2014) containing 1% agar and 0.7% agarose to prevent animals from burrowing. Nearly all *C. tropicalis* isotypes are available from the *Caenorhabditis* Natural Diversity Resource (CaeNDR, release 20250627), excluding the Costa Rican strains that are covered by restrictive permits (Crombie et al. 2024).

To assess potential strain contamination, we examined the correspondence between geographic origin and genome-wide genetic similarity among isotypes. Two isotypes, JU1635 and JU1818, exhibited geographic origins that did not match with those of their closest genetic relatives, suggesting a possible strain swap during laboratory processing. These two strains arrived in the laboratory in the same batch and were processed on overlapping dates, a pattern consistent with a simple sample exchange. Accordingly, they were flagged as potentially misidentified and pending further verification by resequencing. A third isotype, ECA806, despite being collected from the Hawaiian Islands, showed unexpected genetic relatedness to African isotypes. However, detailed examination of laboratory records indicated that ECA806 was collected earlier, processed independently from the African strains, handled by different personnel, and sequenced in a separate pool. Therefore, laboratory contamination of ECA806 is not supported by laboratory records.

### Short-read DNA sequencing

DNA extraction was performed following established protocols (Cook et al. 2016). In brief, to extract DNA, we transferred nematodes from three 10 cm NGMA plates spotted with OP50 *E. coli* into a 15 ml conical tube by washing with 10 mL of M9. We then used gravity to settle animals on the bottom of the conical tube, removed the supernatant, and added 10 mL of fresh M9. We repeated this wash method three times to serially dilute the *E. coli* in the M9 and allow the animals time to purge ingested *E. coli*. Genomic DNA was isolated from 100 to 300 µl nematode pellets using the DNAEasy Blood and Tissue isolation kit (cat# 69506, QIAGEN, Valencia, CA). The DNA concentration was determined for each sample with the Qubit dsDNA Broad Range Assay Kit (cat# Q32850 Invitrogen, Carlsbad, CA). All 785 strains were used to construct short-read libraries using NEBNext® Ultra™ II FS DNA Library Prep (cat# E6177L). These libraries were sequenced on the Illumina NovaSeq 6000 (paired-end 150 bp reads) in Duke Center for Genomic and Computational Biology, Novogene, NUSeq, and the New York University core facility. The raw short-read sequencing data are available from the NCBI Sequence Read Archive (Project PRJNA1127592).

### Long-read DNA sequencing

The reference strain (NIC58) was sequenced by Oxford Nanopore at DNA Technologies and Expression Analysis Cores at the University of California, Davis). A total of 27 other strains were sequenced using PacBio HiFi at Cold Spring Harbor Laboratory (ECA1307, ECA1518, ECA1599, ECA797, JU1373, NIC1594, NIC1649, NIC491, NIC975, QG1004, QG2899), Duke Center for Genomic and Computational Biology (JU1634, NIC125, QG3856), or Maryland Genomics (ECA790, ECA796, EG6180, JU1630, JU1632, NIC1656, NIC85, NIC957, QG3521, QG3790, QG3800, QG5652, QG834). Nematodes were collected using the same protocol as for short-read DNA sequencing, except that 14 10 cm plates were used for NIC58 and 15 10 cm plates were used for each other wild strain, rather than three plates (see *Short-read DNA sequencing)*. Animals for each strain were washed from the plates with 25 ml of M9 and pooled into 50 ml conical tubes to settle by gravity. Worm pellets were washed three times using M9 and then frozen at −80°C. Worm pellets for PacBio sequencing (Revio platform) were submitted to the Cold Spring Harbor Laboratory Genome Center and for Oxford Nanopore sequencing (PromethION platform) to the DNA Technologies and Expression Analysis Cores at the University of California, Davis for genomic DNA extraction, library preparation, and sequencing.

### NIC58 Hi-C library preparation

A Hi-C library for *C. tropicalis* NIC58 was prepared using a modified protocol adapted from a previously published method(Crane et al. 2015). Approximately 12,000 adult nematodes were collected and washed in M9 buffer, then crosslinked with 2% (v/v) formaldehyde. The crosslinked sample was dounced in 1 ml lysis buffer (10 mM Tris–HCl, pH 8.0; 10 mM NaCl; 0.1% [v/v] protease inhibitors) to disrupt nematode pellets. Chromatin was digested overnight with *Dpn*II, followed by incubation at 65°C for 15 minutes to deactivate the enzyme. DNA ends were biotinylated at 23°C for four hours and blunt-end ligated using T4 DNA ligase at 16°C for four hours. Crosslinks were reversed and proteins degraded by adding Proteinase K (50 µl of 10 mg/ml) to the sample. DNA was purified using phenol-chloroform extraction in equal volumes (1:1) of phenol and chloroform in 15 ml phase-lock tubes, followed by centrifugation at 1500 g for five minutes. The aqueous phase was transferred to a 35 ml tube and DNA was precipitated with 10% volume of 3 M sodium acetate and 2.5 volumes of ice-cold 100% ethanol. The pellet was air-dried at room temperature and resuspended in 5 ml of TE buffer. Biotin was removed from unligated DNA ends by treating 5 µg of DNA with 5 µl T4 DNA polymerase (NEB), after which DNA was sheared to 100–300 bp fragments using a Covaris M220 apparatus. Biotinylated fragments were captured with streptavidin beads and resuspended in a ligation buffer, followed by adapter and Illumina index ligation. Beads were pelleted using a Magnetic Particle Concentrator (Thermo Fisher Scientific), washed multiple times, and resuspended in 20 µl NEBuffer 2 (New England Biolabs). The final library was prepared using the Illumina TruSeq kit and sequenced on an Illumina HiSeq 4000 platform to generate approximately 180 million 50 bp paired-end reads.

### NIC58 short-read RNA-seq

Illumina RNA-seq libraries for different developmental stages and both males and hermaphrodites of *C. tropicalis* NIC58 were prepared simultaneously in a single 96-well plate. Total RNA (1 µg) was subjected to mRNA purification and enrichment using the NEBNext Poly(A) mRNA Magnetic Isolation Module (New England Biolabs, catalog no. E7490L). RNA fragmentation, first- and second-strand cDNA synthesis, and end-repair were performed using the NEBNext Ultra II RNA Library Prep Kit with Sample Purification Beads (New England Biolabs, catalog no. E7775L). The resulting cDNA library was adapter-ligated using adapters and unique dual indexes from the NEBNext Multiplex Oligos for Illumina (New England Biolabs, catalog no. E6440L). All steps were carried out according to the manufacturer’s instructions. Library concentration was quantified using the Qubit dsDNA BR Assay Kit (Invitrogen, catalog no. Q32853). The libraries were pooled and quality assessed using a 2100 Bioanalyzer (Agilent) at Novogene (CA, USA), followed by sequencing on a single lane of an Illumina NovaSeq 6000 platform to generate 150 bp paired-end reads.

### NIC58 long-read RNA-seq

A PacBio Iso-Seq full-length transcript sequencing library for *C. tropicalis* NIC58 was prepared using 300 ng of total RNA. Library preparation was performed at the Duke Center for Genomic and Computational Biology’s Sequencing and Genomic Technologies Core Facility using the NEBNext Single Cell/Low Input cDNA Synthesis and Amplification Module (New England Biolabs, catalog no. E6421) and the SMRTbell Express Template Prep Kit 2.0 (Pacific Biosciences, catalog no. 100-938-900). The library was sequenced on three SMRT cells.

### NIC58 reference genome assembly

Oxford Nanopore Technologies (ONT) sequencing data for NIC58 were downsampled to approximately 200x coverage using FiltLong (v0.2.0; https://github.com/rrwick/Filtlong), assuming a genome size of 86 Mb. The subsampled long reads were assembled using Flye v2.8.1-b1676 (Kolmogorov et al. 2018) using the longest 100x reads for disjointig assembly (--asm-coverage 100). Assemblies were aligned to the previous version of the NIC58 genome using nucmer v3.1 (Delcher et al. 2003). Among the assemblers tested, Flye consistently produced the most contiguous assemblies. ONT reads were aligned to the Flye assembly using minimap2 v2.17-r941 (Li 2018), and the resulting alignments were used by racon v1.4.13 (Vaser et al. 2017) for error correction with ONT-recommended parameters (−m 8 −x −6 −g −8 −w 500). The corrected assembly and ONT reads were subsequently processed with Medaka v1.1.2 (https://github.com/nanoporetech/medaka) for additional consensus polishing using the ‘r941_prom_high_g360’ model. Remaining errors were corrected by aligning paired-end Illumina reads to the assembly using bwa mem v0.7.17-r1188 (Li 2013) and polishing with Pilon v1.23 (Walker et al. 2014) using the --fix bases option.

### Hi-C scaffolding

To scaffold the polished assembly into complete chromosomes, Hi-C data were downsampled to approximately 50x coverage using seqtk v1.3-r106 (https://github.com/lh3/seqtk). The Juicer/3D-DNA pipeline (Juicer v1.6; 3D-DNA v180114) was used to align the Hi-C data to the assembly and generate chromosomal scaffolds using default parameters, specifying *Dpn*II as the restriction enzyme (Durand et al. 2016; Dudchenko et al. 2017). We observed that 3D-DNA incorrectly fragmented contigs within highly repetitive regions, a behavior that could not be prevented by disabling misjoin correction (−r 0). To address this issue, the early-exit option in 3D-DNA (–early-exit) was used, followed by manual editing of the assembly file to construct large chromosomal scaffolds guided by the Hi-C contact map.

### Curation and QC

Base-level accuracy of the Hi-C-scaffolded assembly was evaluated by estimating the QV score using Merqury v1.1 (Rhie et al. 2020) in combination with the Illumina short-read dataset. Unplaced contigs were manually examined along with their associated reads using *gap5* (Bonfield and Whitwham 2010), and each contig was also aligned to the assembly using BLASTN. This analysis revealed that all unplaced contigs consisted of redundant repetitive sequences. Consequently, all unplaced contigs were excluded from the final assembly. In addition, read alignments at chromosome ends were manually inspected in *gap5* to identify soft-clipped telomeric repeat sequences. The *gap5* realign function was then used to derive an updated consensus sequence at these regions, adding a total of 101,196 bases to the assembly. As a result, all chromosomal scaffolds terminated in telomeric repeats except for V left and X right. The curated chromosomal scaffolds were subsequently reoriented relative to the previous *C. tropicalis* NIC58 reference genome (Noble et al. 2021).

### NIC58 protein-coding gene prediction

Protein-coding gene models were generated using methods described previously in *C. briggsae* (Stevens et al. 2022). Briefly, short-read RNA-seq data were aligned to the NIC58 reference soft-masked genome assembly using STAR v2.7.3a (Dobin et al. 2013) in two-pass mode, with a maximum intron length of 10 kb. The resulting alignments, together with the soft-masked genome, were provided to the BRAKER pipeline v2.1.6 (Hoff et al. 2019) for gene prediction. In parallel, high-quality transcripts were generated from long-read RNA sequencing data using the IsoSeq3 pipeline v3.4.0 (https://github.com/PacificBiosciences/IsoSeq). PacBio high-quality transcripts were aligned to the genome assembly using minimap2 (Li 2018), and transcriptome assembly was performed with StringTie v2.1.2 (Kovaka et al. 2019). Coding sequences (CDS) were predicted from assembled transcripts using TransDecoder v5.5.0 (https://github.com/TransDecoder/TransDecoder). StringTie gene models with incomplete CDS were identified and extracted using the *agat_sp_remove_incomplete_gene_models.pl* script from AGAT v0.8.1 (Dainat et al. 2024). CDS were repaired for most incomplete models using *agat_sp_fix_longest_ORF.pl*, and any remaining incomplete models were subsequently removed. Complete StringTie models were then merged with repaired models using *agat_sp_merge_annotations.pl*. StringTie gene models containing more than one noncoding exon were excluded, resulting in a final curated set of StringTie annotations. These models were merged with BRAKER gene predictions using *agat_sp_merge_annotations.pl*. Gene features with overlapping CDS were fused using *agat_sp_fix_overlapping_genes.pl*. Because AGAT only resolves overlaps when genes share a CDS, genes that were fully overlapped by others but lacked a shared CDS were identified, removed based on their coordinates using *agat_sp_filter_records_by_coordinate.pl*, and replaced with the original BRAKER predictions in their respective coordinates. Finally, redundant isoforms with identical CDS and intron structures were removed, and all annotation feature identifiers were renamed with consistent prefixes using *agat_sp_manage_IDs.pl*.

### Alignments and variant calling

Adapters and low quality sequences were removed from raw reads using *fastp* (v0.20.0) and default parameters (Chen 2023). Reads shorter than 20 bp after trimming were discarded. Subsequently, a nanopore-based assembly of the *C. tropicalis* NIC58 genome (June 2021) (Noble et al. 2021) was used as the reference for alignment with *Burrows-Wheeler Aligner, BWA* (v0.7.17) (Li and Durbin 2009). *Sambamba* was used to merge and index libraries of the same strain (v0.7.0) (Tarasov et al. 2015), and duplicate reads were flagged using Picard (v2.21.3) (Anon 2019). Strains with an average sequencing depth below 10x were excluded from subsequent analyses.

Variant calling for each strain was performed using the *HaplotypeCaller* module in *GATK* (v4.1.4.0) (Poplin et al. 2018). The resulting gVCF files were combined and jointly genotyped using *GenomicsDBImport* and *GenotypeGVCFs* in *GATK* (v4.1.4.0) (Poplin et al. 2018). The aggregated variants were further processed and filtered using the Andersen Lab variant calling pipeline (https://github.com/AndersenLab/wi-gatk), incorporating custom scripts (Cook 2020), *GATK* (v4.1.4.0), and *bcftools* (v1.10) (Danecek et al. 2021). Because *C. tropicalis* is a highly selfing species, heterozygous single nucleotide variants (SNVs) are likely to represent sequencing or mapping errors. To correct for this potential source of error, biallelic heterozygous SNVs were converted to homozygous reference (REF) or alternate (ALT) alleles when sufficient read support justified this change. Specifically, SNVs were adjusted based on normalized Phred-scaled likelihoods (PL), where smaller PL values indicate higher confidence: heterozygous sites were converted to homozygous ALT if PL-ALT/PL-REF ≤ 0.5 and PL-ALT ≤ 200, and to homozygous REF if PL-REF/PL-ALT ≤ 0.5 and PL-REF ≤ 200. SNVs that did not meet these criteria were left unchanged. At the site level, variants were filtered to ensure quality (QUAL > 30), quality normalized by depth (QD > 20), limited strand bias (SOR < 5; FS < 100), missing genotype fraction < 95%, and post-polarization heterozygosity < 10%. At the sample level, variants with read depth (DP) ≤ 5 were excluded. Most heterozygous sites were converted to missing (./.), except for heterozygous sites on the mitochondrial genome, which were retained. After filtering, invariant sites (those containing only homozygous REF or ALT calls across all samples) were removed. The resulting dataset was referred to as the hard-filtered VCF file.

### Isotype characterization

Highly related strains were grouped into the same group called isotypes following a standardized procedure (https://github.com/andersenlab/isotype-nf). Pairwise genetic similarity among strains was calculated using the hard-filtered VCF file and *bcftools gtcheck*, defined as the ratio of shared variants to total variants for each strain pair (total SNVs: 1,699,166). Isotype assignment was performed by hierarchical agglomerative clustering of strain pairwise genetic similarity using complete linkage. A cutoff of 99.9915% similarity was used to cut the tree resulting in isotype groups with all pairwise genetic similarities exceeding this threshold. Using this approach, no strain was assigned to more than one isotype group. The threshold was chosen to partition strains into genetically homogenous and geographically isolated groups. Unlike *C. briggsae* and *C. elegans*, which have lower thresholds for calling isotype groups, *C. tropicalis* shows much less genetic heterogeneity across the population thus requiring a more stringent cutoff. Although we acknowledge that the cutoff and isotype concept are arbitrary, certain situations are confounded by high relatedness between individual strains, such as genome-wide association studies or summarizing inter-population genetic differences. In these contexts, it is useful to reduce the highly related individuals to a representative strain (Fig. S37).

### Relatedness of isotype reference strains

We analyzed relatedness among *C. tropicalis* isotype reference strains using an LD-pruned VCF derived from the hard-filtered VCF through the following steps. Genome-wide biallelic SNVs from all chromosomes were first extracted and normalized by merging separate biallelic sites into multiallelic records with *bcftools norm -m +* (bcftools v1.14). Biallelic SNVs were then extracted and LD-pruned in PLINK v1.90b6.21 using --indep-pairwise 50 10 0.9 to remove highly correlated markers (Purcell et al. 2007; Chang et al. 2015). After excluding sites with missing data, the remaining loci were extracted from the normalized VCF to produce the final LD-pruned VCF containing only biallelic, non-missing variants. PCA was conducted with smartpca (EIGENSOFT v7.2) (Patterson et al. 2006; Price et al. 2006). The PED, SNP, and IND files generated by PLINK were used as input to compute Eigenstrat values and Tracy-Widom statistics. In total, 213 principal components were required to explain 80% of the genetic variance, and 160 components had eigenvalues greater than 1.

### Species-wide tree construction

We converted the LD-pruned VCF files into PHYLIP format using the *vcf2phylip.py* script (Ortiz 2019). Tree reconstruction was performed with IQ-TREE2 v2.4.0 (Minh et al. 2020) using the PHYLIP (Felsenstein 1993) file as input. The GTR+F+ASC+R5 model was selected as the best-fit maximum-likelihood model under the Bayesian information criterion (BIC), with the search restricted to extensions of the GTR model. Node support was evaluated with 1,000 ultrafast bootstrap (UFBoot2) replicates. The resulting tree was visualized in R v4.3.2 using the ggtree v3.10.1 package (Yu et al. 2017). Pairwise genetic similarity among 622 *C. tropicalis* isotypes, as well as among subsets of isotypes, was clustered and visualized using the ComplexHeatmap v2.18.0 package in R (Gu 2022). The Neighbor-joining network was also constructed using LD-pruned PHYLIP format file based on p-distances using in SplitsTree v6.4.13 (Huson and Bryant 2024).

### GIS and environmental data analysis

Haversine distances between sampling locations were calculated using the *distm()* function in the geosphere R package (v1.5-20) (Hijmans 2024). The publicly available geographic information system (GIS) data was processed in R (v4.3.2) using the terra (v1.8-60) (Hijmans 2025) and sf (v1.0-21) (Pebesma 2018) packages. For Taiwan, four GIS datasets (mean annual temperature in °C, mean annual rainfall in mm, elevation in m, and area solar radiation in WH/m²) were obtained from an open-access environmental dataset of 1 km grid data for average values from 2010 to 2013 (Wan-Jyun et al. 2020). For the Hawaiian Islands, data on mean annual rainfall, air temperature, surface temperature, soil moisture, leaf area index (LAI), and elevation were extracted from publicly available databases. Mean annual rainfall was obtained from the Rainfall Atlas of Hawaii at approximately 250 m resolution (Giambelluca et al. 2013; Frazier et al. 2016). Elevation data were obtained from the United States Geological Survey (USGS) 3D Elevation Program (3DEP) seamless digital elevation model at 1/3 arc-second resolution (about 10 m). Other average annual climate variables at our sampling sites, including air temperature, surface temperature, soil moisture, and LAI were obtained at 250 m resolution from the Hawaii Climate Data Portal (HCDP).

### Admixture analysis

We ran the admixture analysis using ADMIXTURE v1.3.0 (Alexander et al. 2009) with a PED file as input. This PED file includes the LD-pruned sites exported by PLINK when generating the LD-pruned VCF. To identify the best K, we ran ten independent ten-fold cross-validations and chose the point where the CV error curve first reached a stable minimum. We tested K from 2 to 30. Non-admixed representative strains were defined using a maximum subpopulation fraction above 99.9%. We minimized the cross-validation (CV) error by comparing CV estimates across 10 runs for each assumed K value (from K = 2 to 30) and found support for at least 28 subpopulations (Fig. S8, S9). At K=28, non-admixed individuals from each subpopulation were assigned uniquely to one relatedness group, with certain relatedness groups containing several subpopulations (Fig. S10). Across ten independent runs under K=28, we observed that eight relatedness groups were sometimes assigned to the same subpopulation, although the remaining relatedness groups consistently stayed discrete (Fig. S11).

### Repeat Masking

We identified *C. tropicalis* genome-wide repetitive elements using methods described previously (Stevens et al. 2022) with minor modifications. First, repetitive sequences were modeled *de novo* using RepeatModeler from RepeatMasker v2.0.6 (Smit et al. 2015). Transposable elements were then identified using TransposonPSI v1.0.0 (http://transposonpsi.sourceforge.net/). Long terminal repeat (LTR) retrotransposons were identified with LTRharvest and then annotated with LTRdigest from GenomeTools v1.6.5 (Ellinghaus et al. 2008; Gremme et al. 2013), with HMM profiles from the Gypsy Database v2.0 (Llorens et al. 2011) and Pfam v37.4 domains (Mistry et al. 2021). Repetitive sequences were then further filtered with the gt-select tool to remove sequences lacking conserved protein domains. We then obtained all Rhabditid repeats from RepBase (Bao et al. 2015) and Dfam (Hubley et al. 2016). The generated repeat libraries were merged into a single redundant repeat library, clustered with VSEARCH v2.22.1 (Rognes et al. 2016) from QIIME2 v2024.10 (Bolyen et al. 2019), and then classified with the RepeatClassifier tool from RepeatModeler. Finally, the unclassified repeats that had BLASTX v2.16.0 hits to *C. elegans* or *C. tropicalis* proteins were removed using RepeatMasker to eliminate genuine gene sequences that had been misidentified as repeats.

### Population genomic statistics

Genome-wide nucleotide diversity (π), Watterson’s θ (θ_W_), and Tajima’s D were all calculated with scikit-allel v1.3.5 (Miles et al. 2024) per 10 kb window using the hard-filtered VCF file as the input. Only SNV positions were kept; repeat regions and sites with over 80% missing genotype data were excluded. To avoid bias caused by structural variation and indels that alter the number of callable sites when calculating π, θ_W_, and Tajima’s D using scikit-allel, we masked the inaccessible sites (Korunes and Samuk 2021). For π and θ_W_, inaccessible sites within each window were masked using the *is_accessible* parameter so that denominators reflected the number of callable bases. For Tajima’s D, genotypes at inaccessible sites were masked before calculation. D_xy_ between relatedness groups were calculated using Pixy v1.2.11.beta1 (Korunes and Samuk 2021) in 10 kb windows, with the hard-filtered VCF as the input. Accessibility to variable sites in the denominator is handled by default settings in Pixy (Korunes and Samuk 2021).

### Long-read genome assembly and protein-coding gene prediction

Sequenced PacBio HiFi genomic reads were demultiplexed and adapters were removed by the sequencing cores using using *lima* (Duke: v2.9.0, CSHL: v2.2.0, UMD: v2.12.0 (https://lima.how). Demultiplexed and adapter-trimmed samples from the same strain sequenced from different facilities were merged using *samtools* v1.20 (Danecek et al. 2021). PCR duplicates were removed using *pbmarkdup* v1.1.1 (https://github.com/PacificBiosciences/pbmarkdup). Primary assemblies were generated from deduplicated HiFi FASTAs using *hifiasm* v0.24.0-r702 (Cheng et al. 2021) with the initial bloom filtered disabled (*-f0*, recommended for small genomes) and haplotig purging disabled (*-l0*, recommended for homozygous genomes). Genome assemblies generated from merged samples from the same strain were compared against assemblies generated from each individual sample, and the assembly with the highest N50 was kept. After assembly, contigs were taxonomically classified using DIAMOND 2.1.13 (Buchfink et al. 2021) with parameters *--faster* and an e-value of 1e-10 against the UniProt database (Release 2025_03) (UniProt Consortium 2025) with the inclusion of the *C. tropicalis* reference proteome (taxon ID: 1561998). Contigs that were annotated as non-*Nematoda* were purged using BlobToolKit v4.4.5 (Challis et al. 2020) with *--param bestsumorder_phylum--Inv=Nematoda*. For each assembled genome, protein-coding gene models were generated using BRAKER3 v3.0.8 (Gabriel et al. 2023) guided with protein sequences from *C. elegans* N2 (WormBase, WS283). Genome and gene model biological completeness was estimated using BUSCO v5.0.4 (Simão et al. 2015) with the *nematoda_odb10* database. Summary statistics for each genome assembly are reported (Table S13).

### Identification of hyper-divergent regions

HDRs were identified using methods described previously in *C. elegans* (Lee et al. 2021) with modifications. First, we aligned the 27 newly assembled isotype strain genomes against the reference NIC58 genome using *nucmer* v3.1 (custom parameters --mingap 500 --mincluster 100) (Marçais et al. 2018). The reference genome was then partitioned into 1 kb bins using *bedtools* v2.31.1 (*makewindows*) (Quinlan and Hall 2010), and the identity of each bin was estimated from the average identity of the alignment spanning the bin. When multiple alignments of variable length mapped to a single bin, the alignment with the longest span was selected. Instead of the coverage fraction used previously (proportion of read depth of the bin relative to the genome-wide depth average) (Lee et al. 2021) we estimated the percentage of bases covered by at least 1x in each bin (percent bases covered). Bins under 97% identity or 60% bases covered were classified as hyper-divergent (Fig. S38). Non-hyper-divergent bins that were flanked by two hyper-divergent bins were re-classified as hyper-divergent (gap re-classification) and contiguous bins classified as hyper-divergent were clustered into HDRs (bin clustering).

Next, to determine the optimal parameters to call hyper-divergent regions species-wide using short-read sequencing data, we first aligned the Illumina short-read DNA sequences of the 27 isotypes with long-read genomes against the reference genome and called variants using GATK as described above (see *Alignments and variant calling*). For each strain, we estimated the single-nucleotide variant count using *bcftools* v1.21 (Danecek et al. 2021) and *bedtools coverage* v2.31.1 (Quinlan and Hall 2010), and the percentage of bases covered in every reference bin using mosdepth v0.3.10 (Pedersen and Quinlan 2018). We tested a wide range of variant count (5 to 25 SNVs per kb) and percent bases covered (5% to 90%) thresholds to classify and cluster bins into HDRs using short-read data for the 27 isotype strains where we previously generated long-read HDR calls. We compared the HDR calls from short-read data at each threshold pair against the respective long-read HDR calls for each strain. We estimated the quality of overlaps between short-and long-read HDR calls using overlap fraction (proportion of a long-read call overlapped by a short-read call) and excess fraction (proportion of a short-read call that exceeded the boundary of an overlapping long-read call). For each strain at each threshold pair, we estimated the mean overlap fraction and mean excess fraction for every overlap between the short- and long-read HDR calls (Fig. S39-40). We also estimated the completeness and accuracy of the short-read hyper-divergent calls by calculating recall (proportion of long-read calls that had an overlap with short-read calls) and precision (proportion of short-read calls that had an overlap with long-read calls) (Fig. S41-42). Recall and precision were used to estimate an F1 score for each threshold pair (Fig. S43). We selected the top N (starting at N=1) threshold pairs based on the F1 score for each strain, and increased the value of N by 1 until we identified a threshold pair that was present among all of the 27 strains in this comparison (referred to as ‘consensus optimal’). The consensus optimal threshold pair was reached at N=19 (top 5% of all threshold pairs), where the optimal thresholds selected were a count of 9 variants paired with 90% bases covered (Fig. S44). We classified bins with estimated variant count or percent bases covered under these optimal thresholds as hyper-divergent, genome wide, across all 622 wild isotype strains. We then performed the same gap re-classification and bin clustering steps that were applied to HDRs called with long-read sequencing data. Proximal HDRs separated by less than 5 kb were merged and HDRs with a span smaller than 5 kb after merger were filtered out.

### Functional enrichment analysis

InterProScan v5.75 (Jones et al. 2014) was used to identify functional protein domains and their associated Gene Ontology IDs that are enriched in genes found in hyper-divergent region arm domains and non hyper-divergent region arm domains. InterProScan was run with options --goterms, --iprlookup, and --disable-precalc on the longest-isoform proteome of NIC58 created using *agat_sp_keep_longest_isoform.pl* from AGAT v1.4.1 (Dainat et al. 2024) and gffread v0.12.7 (Pertea and Pertea 2020). A total of 14,260 NIC58 proteins were annotated with InterProScan functional protein domains, and 10,985 proteins were annotated with Gene ontology IDs. Of the 14,260 proteins with domain annotations, genes found on chromosome arms were used as the background set for enrichment analysis, resulting in 5,411 genes being used. Of the 10,985 proteins with Gene Ontology ID annotations, genes found on chromosome arms were used as the background set for Gene Ontology enrichment analysis, resulting in 3,966 genes being used. Of the reference genes found within HDRs on chromosomal arms, approximately 61% (2,603 / 4,245) received InterProScan functional protein domain annotations and approximately 41% (1,750 / 4,245) received gene ontology identifiers. All NIC58 genes found on chromosome arms genes were separated based on their presence or absence in HDRs identified among all 684 *C. tropicalis* strains. For InterProScan domain enrichment analysis, a one-sided hypergeometric test was conducted in R, with a false discovery rate adjusted p-value using the R package *stats* v.4.2.1. InterProScan functional protein domains were classified as significantly enriched if they had an adjusted p-value less than 0.05. Gene Ontology enrichment analysis was performed using the R package clusterProfiler v4.6.2 (Yu et al. 2012; Wu et al. 2021; Xu et al. 2024). Gene Ontology terms were classified as significantly enriched if they met the criteria of having both a Benjamini-Hochberg adjusted p-value and a q-value of less than 0.05. To identify NIC58 orthologs of N2 genes, OrthoFinder v3.1.0 (Emms et al. 2025) was run using proteomes translated from the reference protein-coding gene annotations of NIC58 (Noble et al. 2021) and N2 (WormBase, WS283) using only the longest isoforms for each protein-coding gene model.

## Data availability

The raw short-read sequencing reads for the wild *C. tropicalis* strains are publicly available from the NCBI Sequence Read Archive (project PRJNA1127592). The raw PacBio long-read data are publicly available from the NCBI Sequence Read Archive (project PRJNA1406775). Strain information and short-read genomic variation data are available from the CaeNDR (release 20250627) (www.caendr.org)(Crombie et al. 2024). Scripts and related raw and processed data are available at GitHub (https://github.com/AndersenLab/Ct_pop_gen_project)

## Author contributions

B.W, N.D.M., and E.C.A. conceived and designed the study.

B.W., N.D.M., L.M.O., and E.C.A. analyzed the data and wrote the manuscript.

R.E.T., M.E.G.S., R.M., A.D.C., M.V.R., and M.-A.F. edited the manuscript.

R.E.T., A.K, and N.M.R performed whole-genome sequencing for 785 *C. tropicalis* wild isolates.

R.E.T. and A.K. performed long-read sequencing for 27 *C. tropicalis* wild isolates.

R.E.T., C.B., T.A.C., C.D., K.S.E., D.E.C., G.Z., L.S., N.R., D.L., S.Z, C.G, C.G., M.-E.C., V.D.D., J.W., A.D.C.,

M.V.R., M.-A.F., C.B., and E.C.A. performed sample collection and isolation of 785 *C. tropicalis* strains.

M.E.G.S. performed isotype characterization for 785 *C. tropicalis* wild strains.

L.S. curated the *C. tropicalis* NIC58 reference genome and its gene annotation.

## Acknowledgements

We thank members of the Andersen Lab for providing comments on this manuscript. We especially thank Michael Ailion, Yen-Ping Hsueh, Isabelle Nuez, Fabrice Besnard, Jean David, Alexander Peluffo, Hagus Tarno and Emha Rais for contributing wild *C. tropicalis* strains to CaeNDR.

