## Supplementary Figures for "Global genomic diversity of the selfing nematode *Caenorhabditis tropicalis* correlates with geography"

Figure S1: Global sampling map of *C. elegans*, *C. briggsae*, and *C. tropicalis* showing *C. tropicalis* have been found exclusively in tropical and subtropical regions

Figure S2: Geographic distribution of 622 *C. tropicalis* isotype reference strains collected worldwide

Figure S3: Genome-wide linkage disequilibrium (LD) decay across species

Figure S4: Maximum-likelihood tree of 622 *C. tropicalis* isotype reference strains

Figure S5: Unrooted equal angle maximum likelihood tree of 622 *C. tropicalis* isotype reference strains

Figure S6: Principal Component Analysis (PCA) using each chromosome genotype

Figure S7: Mean genetic similarity within and between relatedness groups

Figure S8: Minimization of cross-validation error across ADMIXTURE runs

Figure S9: Summary of ADMIXTURE analysis

Figure S10: Population structure of 622 global *C. tropicalis* isotype reference strains

Figure S11: The non-admixed isotypes by relatedness groups across 10 ADMIXTURE runs, displaying only the eight relatedness groups (Tw2, Tw3, Tw4, Tw5, Tw6, Mic1, Mic2 and Indo2)

Figure S12. Isolation by distance across global isotypes

Figure S13. Isolation by distance within relatedness groups

Figure S14: Distribution of relatedness groups in Taiwan

Figure S15: Environmental parameters of relatedness groups in Taiwan

Figure S16: Distribution of relatedness groups across the Hawaiian Islands

Figure S17: Environmental parameters of relatedness groups across the Hawaiian Islands

Figure S18: Distribution of relatedness groups in the Caribbean

Figure S19: Distribution of relatedness groups in Central America

Figure S20: The genetic similarity of the strains in Central America

Figure S21: Distribution of relatedness groups in South America

Figure S22: The genetic similarity of the strains in South America

Figure S23: Distribution of relatedness groups in Micronesia

Figure S24: The genetic similarity of the strains in Micronesia

Figure S25: Distribution of relatedness groups in the Malay Archipelago

Figure S26: Distribution of relatedness groups in Africa

Figure S27: The genetic similarity of the strains in Africa

Figure S28: Hyper-divergent regions (HDRs) contain the vast majority of genetic variation within a small proportion of the genome

Figure S29: The extent of HDRs decreases linearly with genetic similarity to the NIC58 reference genome

Figure S30: Divergent Hawaiian relatedness group carries private and shared HDRs relative to the reference LAC relatedness group

Figure S31: Genome alignments between highly divergent isotype pairs in all three selfing species

Figure S32: HDRs display higher or no difference in absolute divergence across different relatedness group comparisons

Figure S33: Pairwise genetic similarity across the HDRs

Figure S34: Pairwise genetic similarity across the non-HDRs

Figure S35: Diversity statistics of non-HDRs along physical genome positions across the six chromosomes

Figure S36: Overlap between previously described toxin-antidote systems (TAs) and HDRs identified in this study

Figure S37: Genetic similarity score distribution and isotype cutoff

Figure S38: Size and identity features of long-read based HDRs at variable identity thresholds

Figure S39: Mean overlap fraction between short- and long-read HDR calls

Figure S40: Mean excess fraction between short- and long-read HDR calls

Figure S41: Precision of short-read HDR calls

Figure S42: Recall of long-read HDR calls

Figure S43: F1 of short-read HDR calls

Figure S44: Strain and consensus optimal threshold pairs for HDR calls

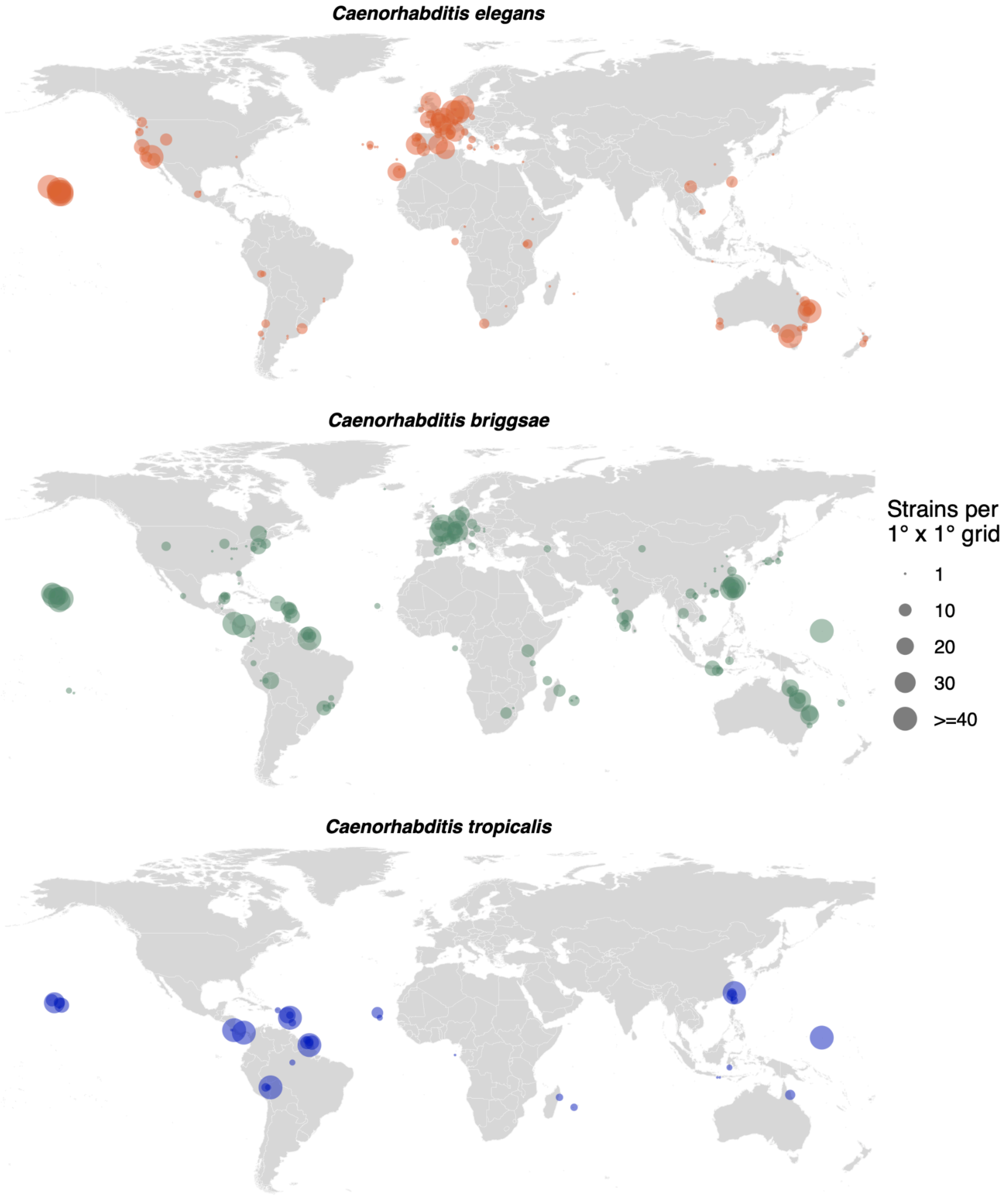


**Figure S1: Global sampling map of *C. elegans*, *C. briggsae*, and *C. tropicalis* showing *C. tropicalis* have been found exclusively in tropical and subtropical regions.** Point size represents the number of strains per 1° x 1° grid. Strain data were based on the latest CaeNDR release (*C. elegans* 20250625, *C. briggsae* 20250626, *C. tropicalis* 20250627).

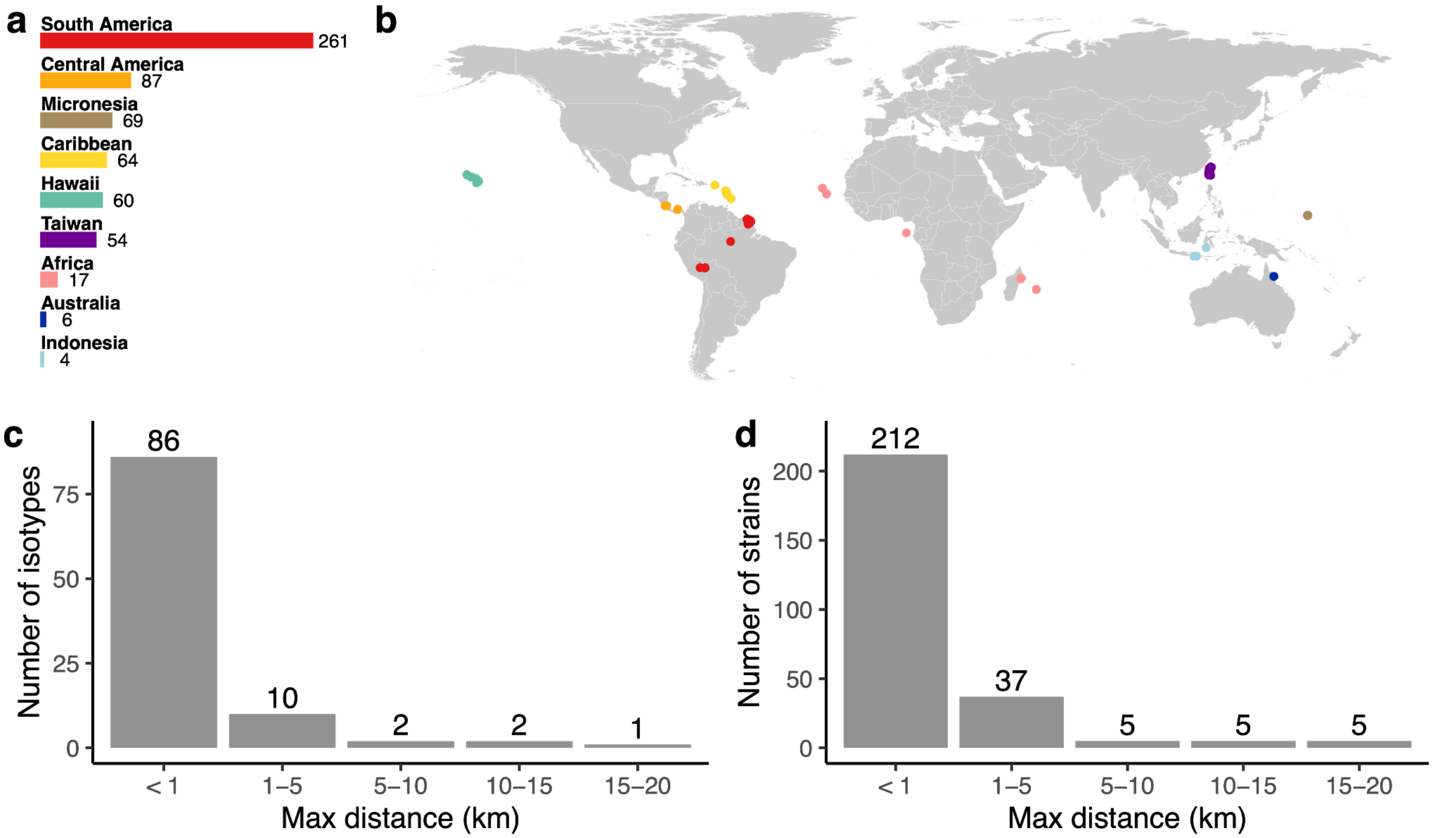


**Figure S2: Geographic distribution of 622 *C. tropicalis* isotype reference strains collected worldwide. a,** Total number of isotypes in each geographic region. **b,** Map showing the isolation locations of isotype reference strains. **c,** Number of isotypes across bins of maximum geographic distance between strains within each isotype. **d,** Number of strains corresponding to isotypes in panel c in each maximum geographic distance bin. The 521 single-strain isotypes are omitted in panels c and d.

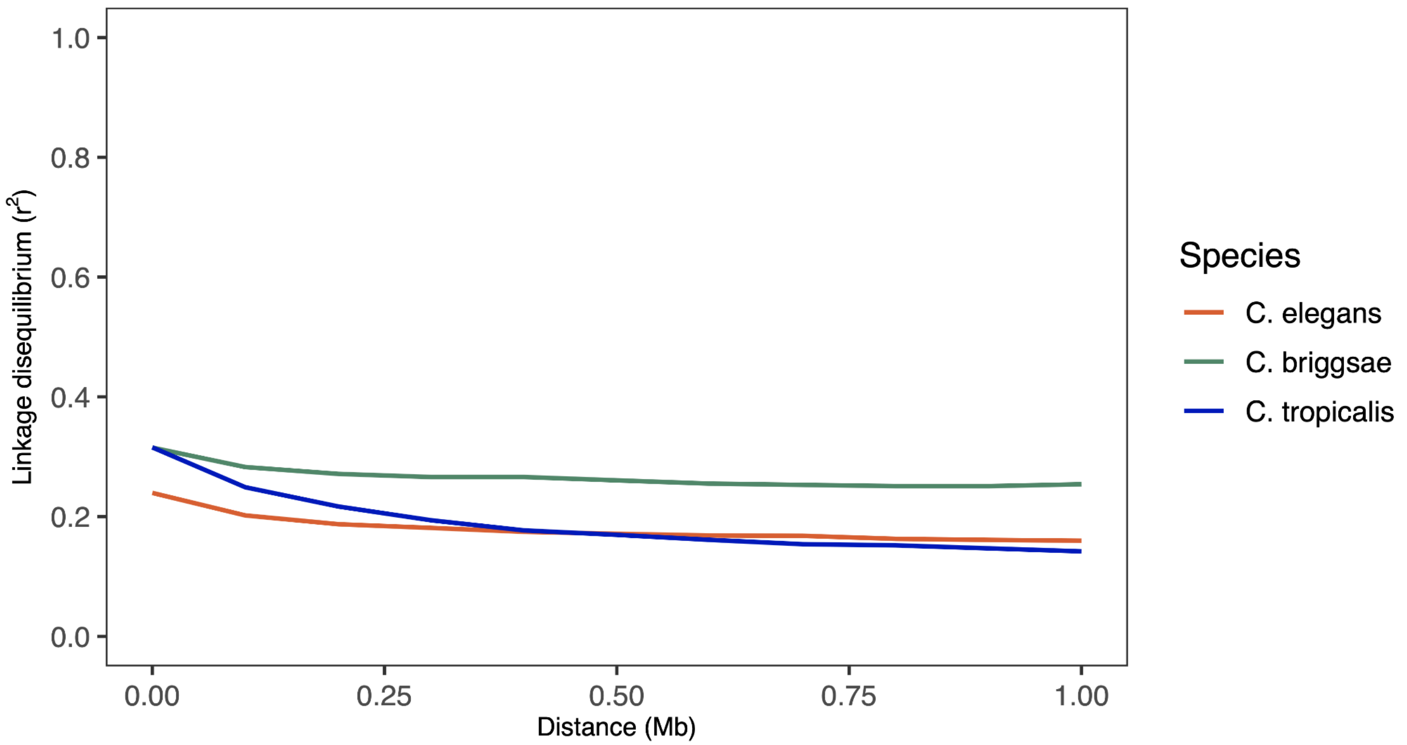


**Figure S3: Genome-wide linkage disequilibrium (LD) decay across species.** The $r^2$ of LD is plotted against physical distance between variant sites for *C. elegans*, *C. briggsae*, and *C. tropicalis*. Lines represent the median.

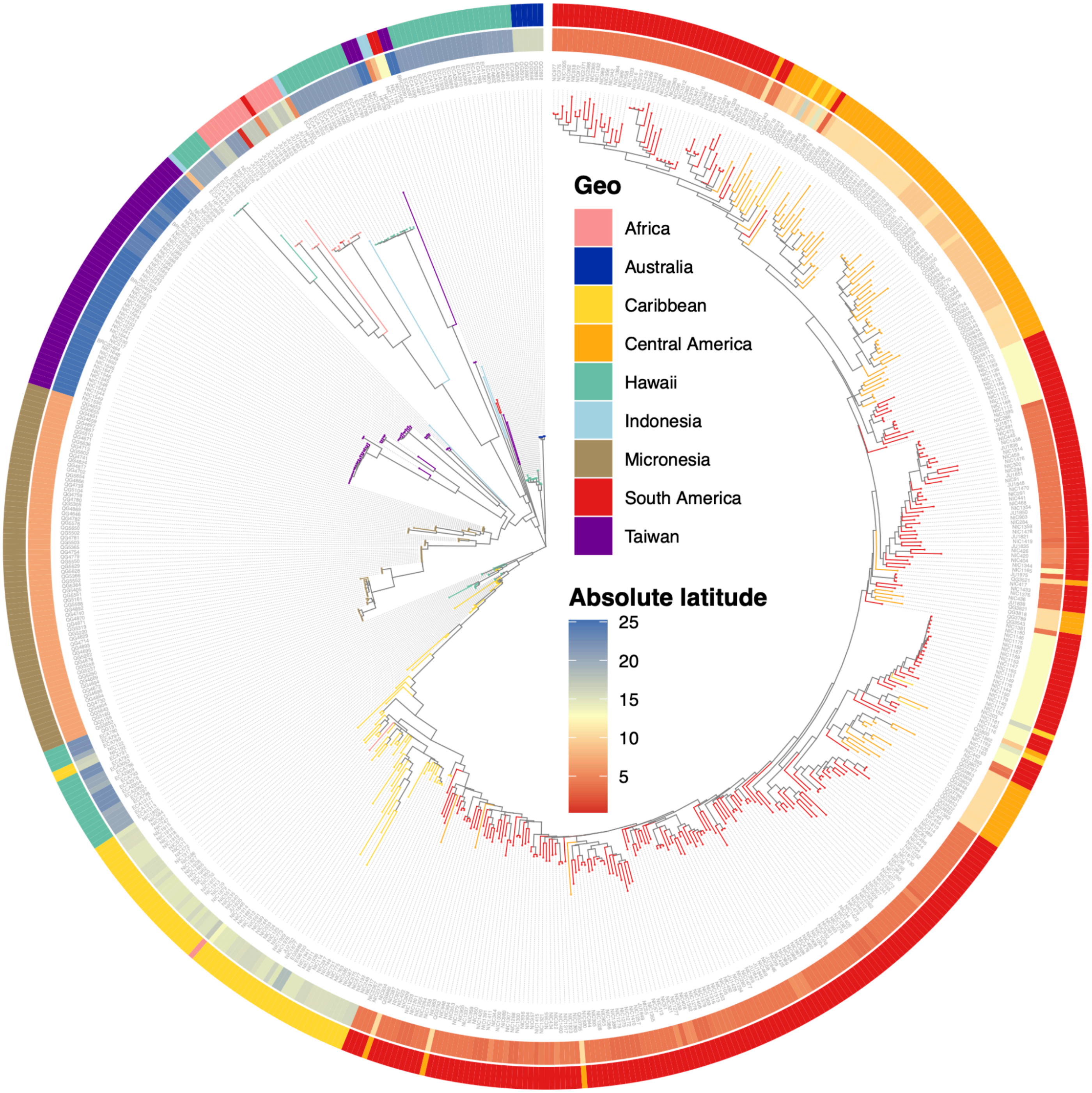


**Figure S4: Maximum-likelihood tree of 622 *C. tropicalis* isotype reference strains.** The tree was generated from LD-pruned variants with $r^2$ value less than 0.9 using the GTR+F+ASC+R5 maximum-likelihood substitution model (see Methods). The color of each leaf branch corresponds to the geographic region of isolation of each strain. The tree is midpoint-rooted. Rings were colored by absolute latitude and geographic region.

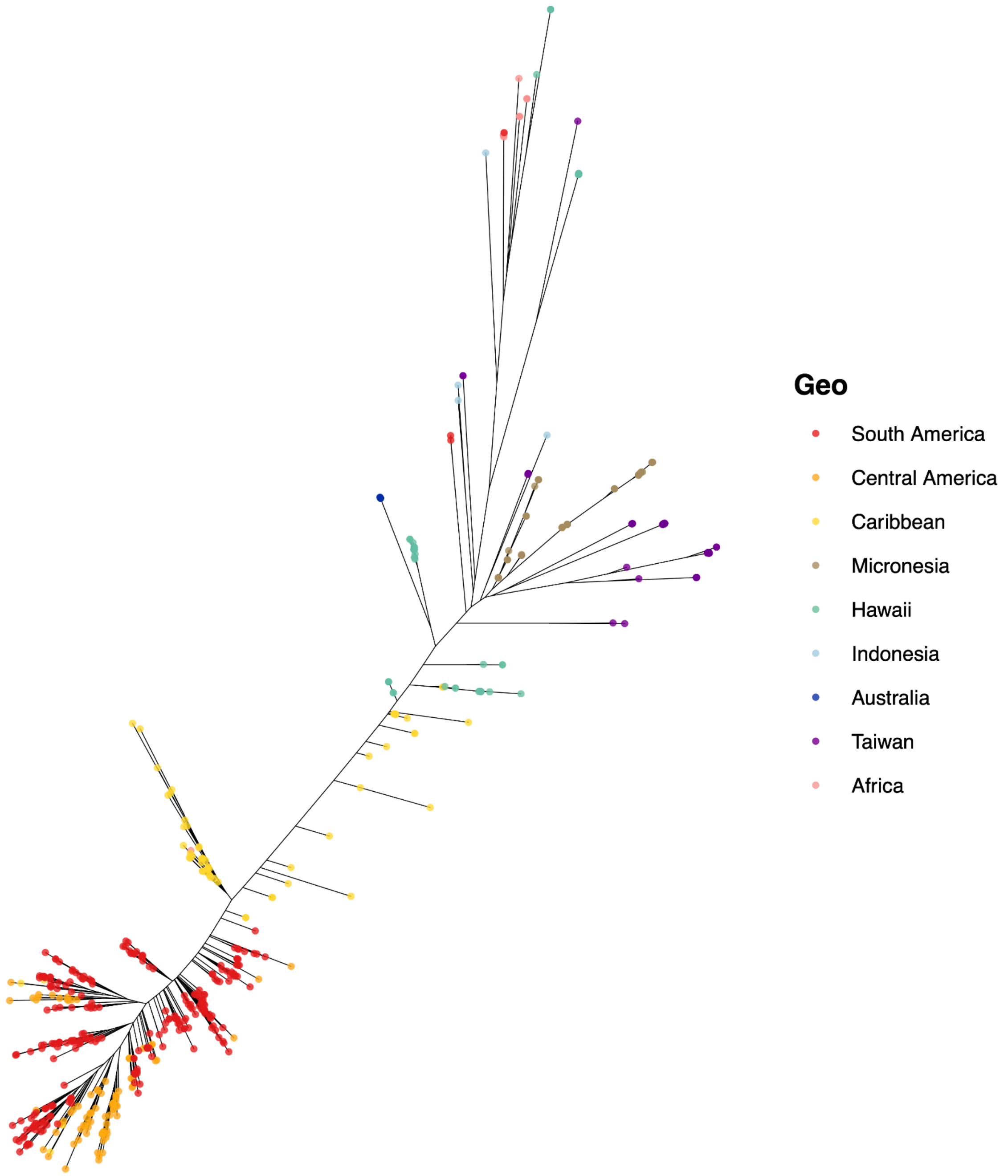


**Figure S5: Unrooted equal angle maximum likelihood tree of 622 *C. tropicalis* isotype reference strains.** The tree was generated from LD-pruned variants with r2 value less than 0.9 using the GTR+F+ASC+R5 maximum-likelihood substitution model (see Methods).

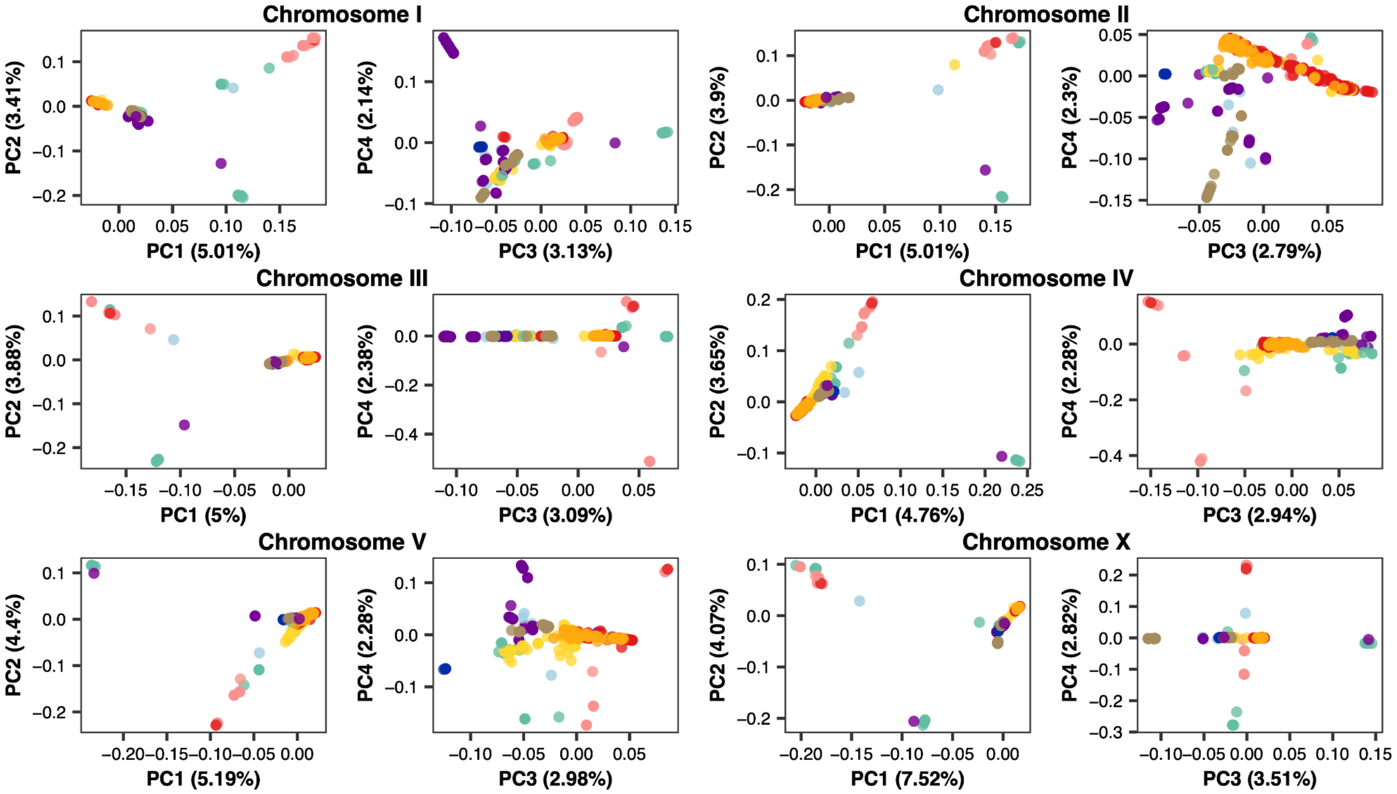


**Figure S6: Principal Component Analysis (PCA) using each chromosome genotype.** Colors in each panel represent the sampling geographic region of each strain.

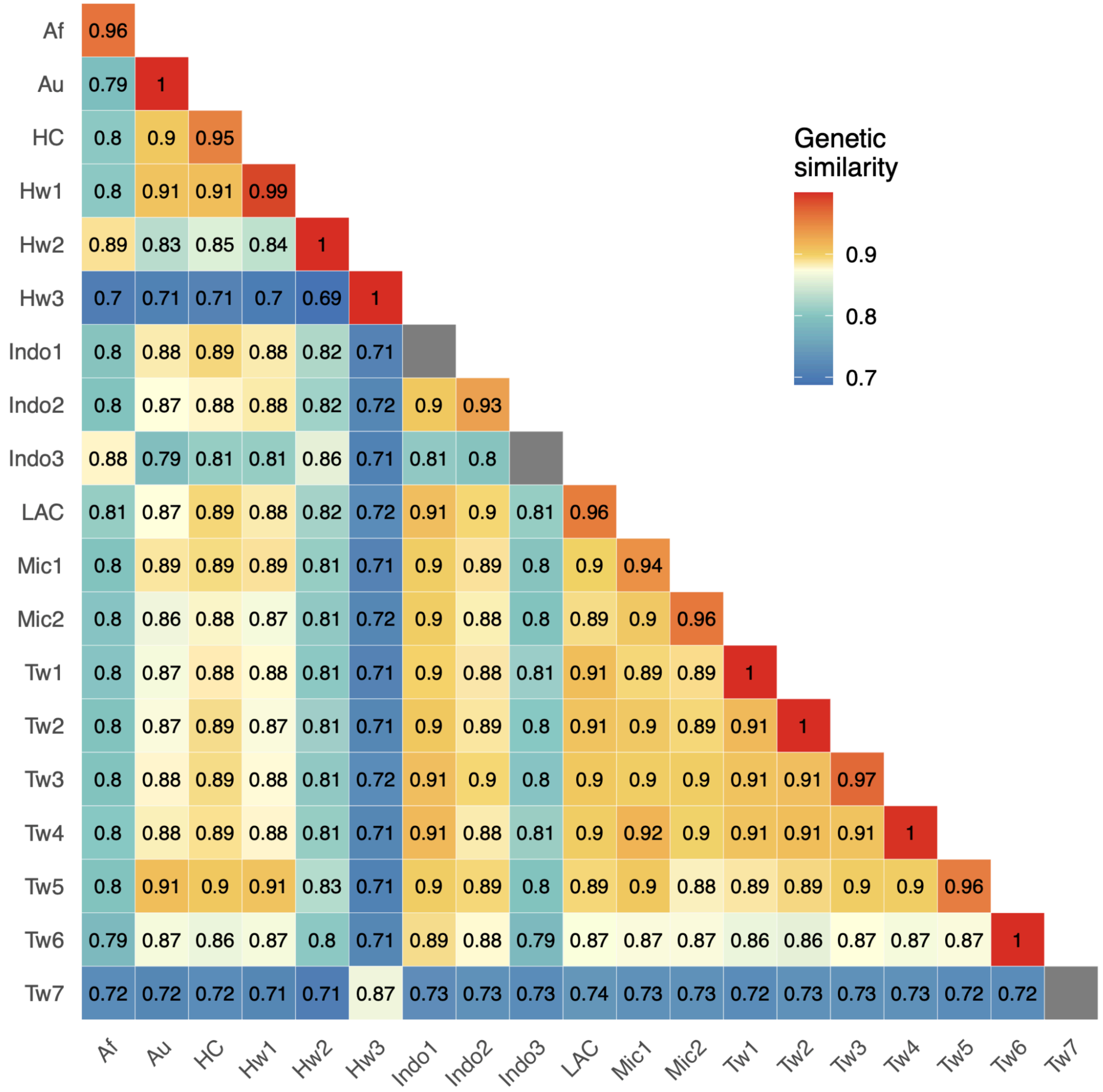


**Figure S7: Mean genetic similarity within and between relatedness groups.** Relatedness groups are shown on the x- and y-axis. Mean estimates of pairwise genetic similarity across all isotypes of any given relatedness group comparison are shown. Genetic similarity is estimated by the proportion of identical alleles at all identified SNV among the 622 isotype strains in this study. Gray squares represent that the group contains only one isotype, so the within-group mean genetic similarity cannot be calculated.

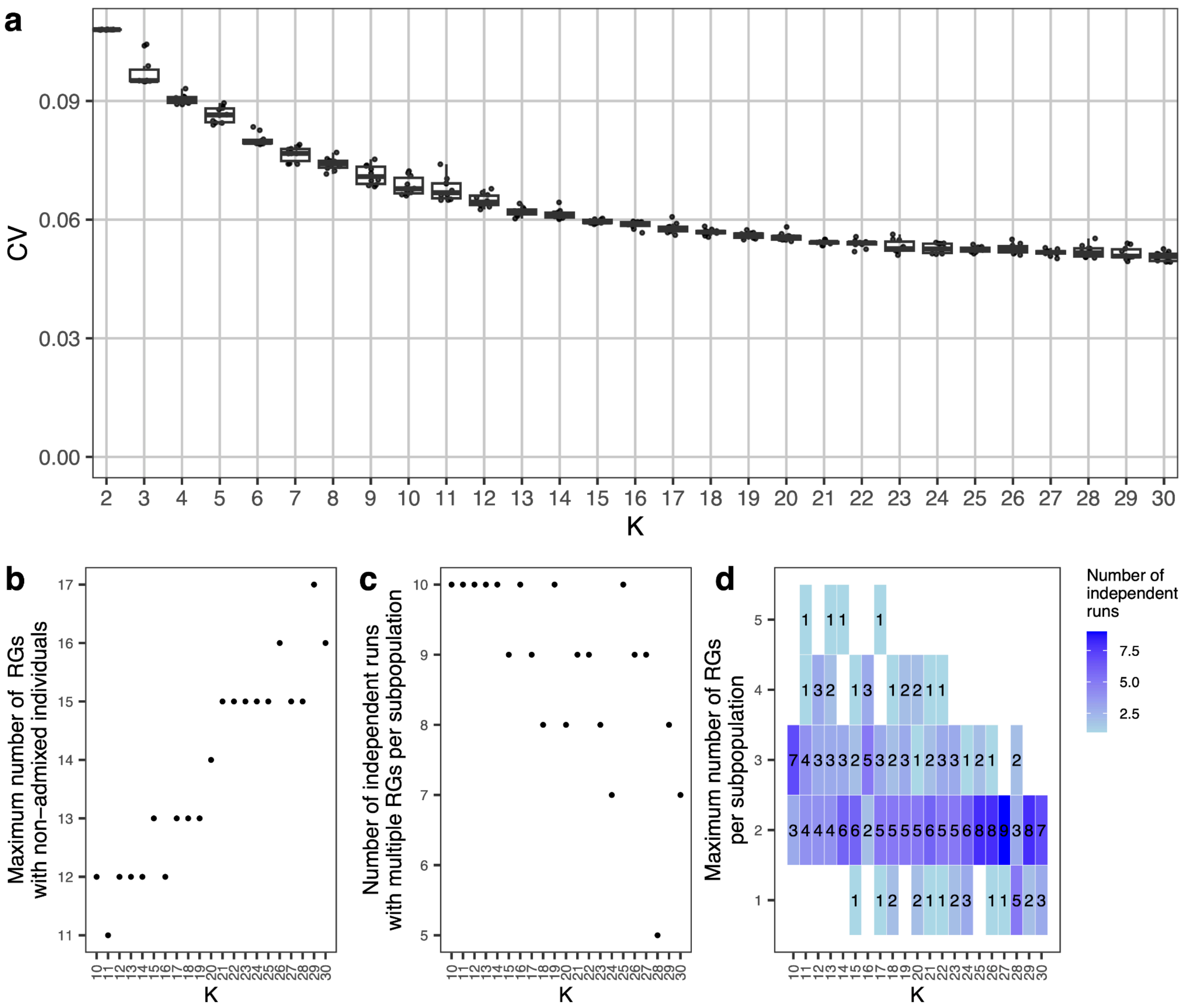


**Figure S8: Minimization of cross-validation error across ADMIXTURE runs. a,** Cross-validation error (y-axis) of ten independent ADMIXTURE runs across K assumed subpopulations. **b,** number of relatedness groups with non-admixed isotypes across K=10 to K=30. **c,** number of ADMIXTURE runs where multiple relatedness groups had individuals assigned to a single subpopulation across K=10 to K=30. **d,** Maximum number of relatedness groups assigned to a single subpopulation across K=10 to K=30.

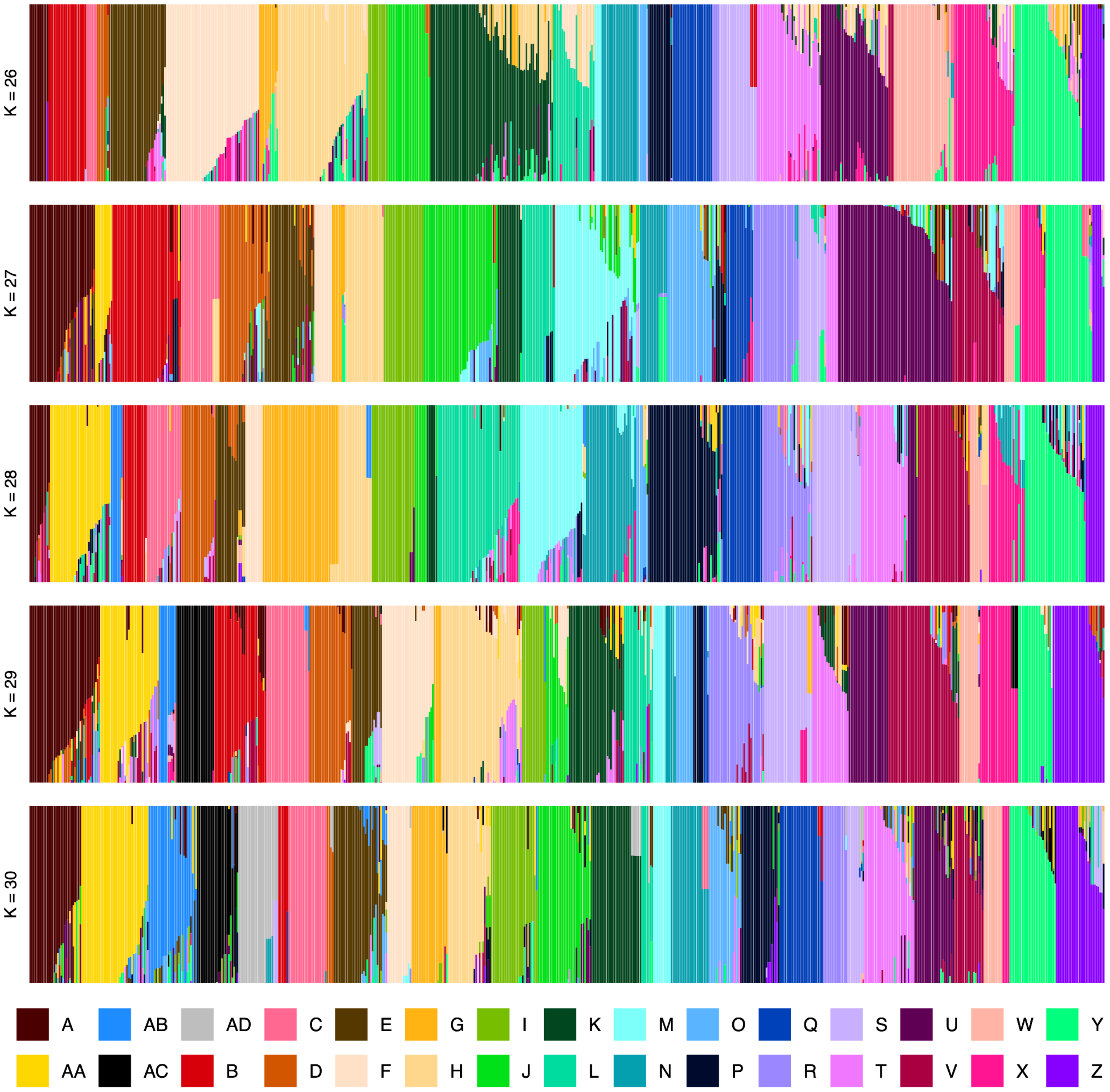


**Figure S9: Summary of ADMIXTURE analysis.** Subpopulation fractions estimated by ADMIXTURE across K ranges ± 2 of best K, 28. Each vertical line represents an isotype, which is partitioned into colored segments that represent the membership fractions for the overall subpopulations.

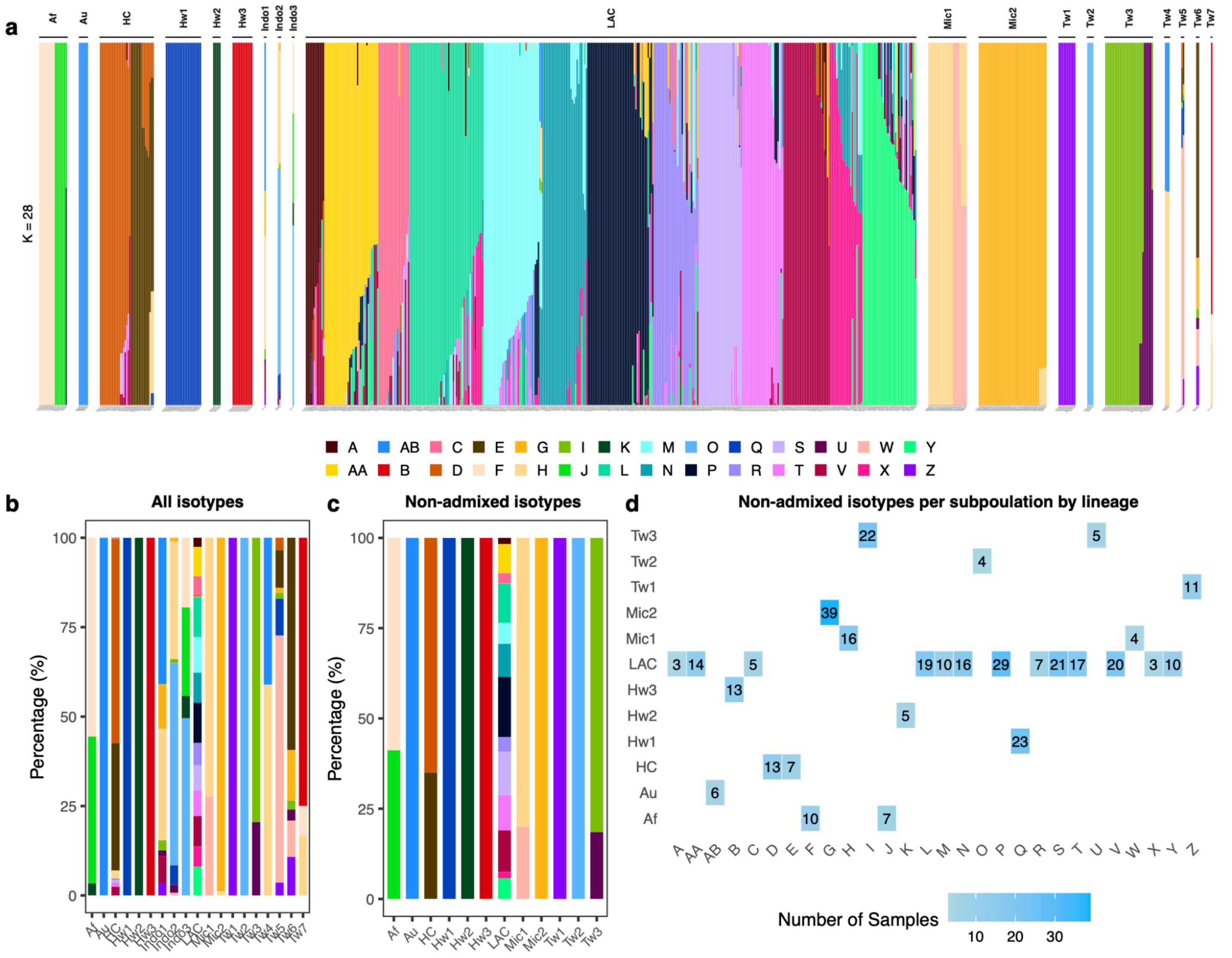


**Figure S10: Population structure of 622 global *C. tropicalis* isotype reference strains. a,** Admixture bar graph of population structure in global *C. tropicalis*. Each column represents an isotype reference strain. The labels above represent the relatedness groups. **b,** Population proportions among all isotype reference strains in each relatedness group. **c,** Population proportions of non-admixed representative isotype reference strains in each relatedness group. **d,** Heatmap for non-admixed representative isotype reference strains per population in each relatedness group. A non-admixed representative isotype reference strain was defined as a strain with a maximum population fraction over 99.9%. The smallest fraction (0.00001) automatically assigned by the software to each subpopulation was removed when plotting the stacked fractions in panel c. For panel d, only the largest fraction within each non-admixed isotype was used to calculate the stacked bars. Every group contains individuals with detectable ancestry from at least one other group except for Tw3, but this pattern does not hold across other runs (See Figure S8).

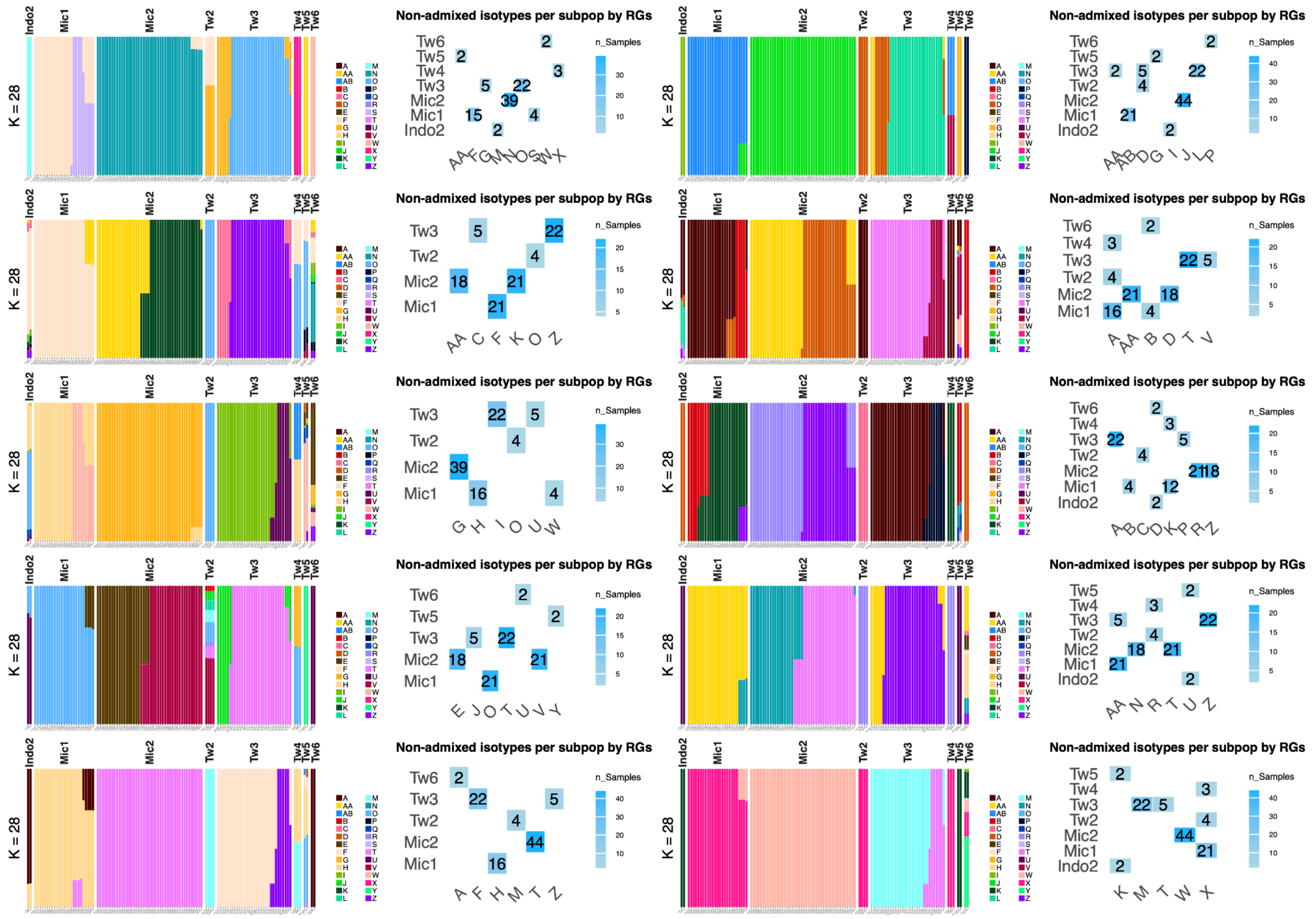

**Figure S11: The non-admixed isotypes by relatedness groups across 10 ADMIXTURE runs, displaying only the eight relatedness groups (Tw2, Tw3, Tw4, Tw5, Tw6, Mic1, Mic2 and Indo2).** Occasionally, the same subpopulation might be assigned to multiple different relatedness groups among these eight.

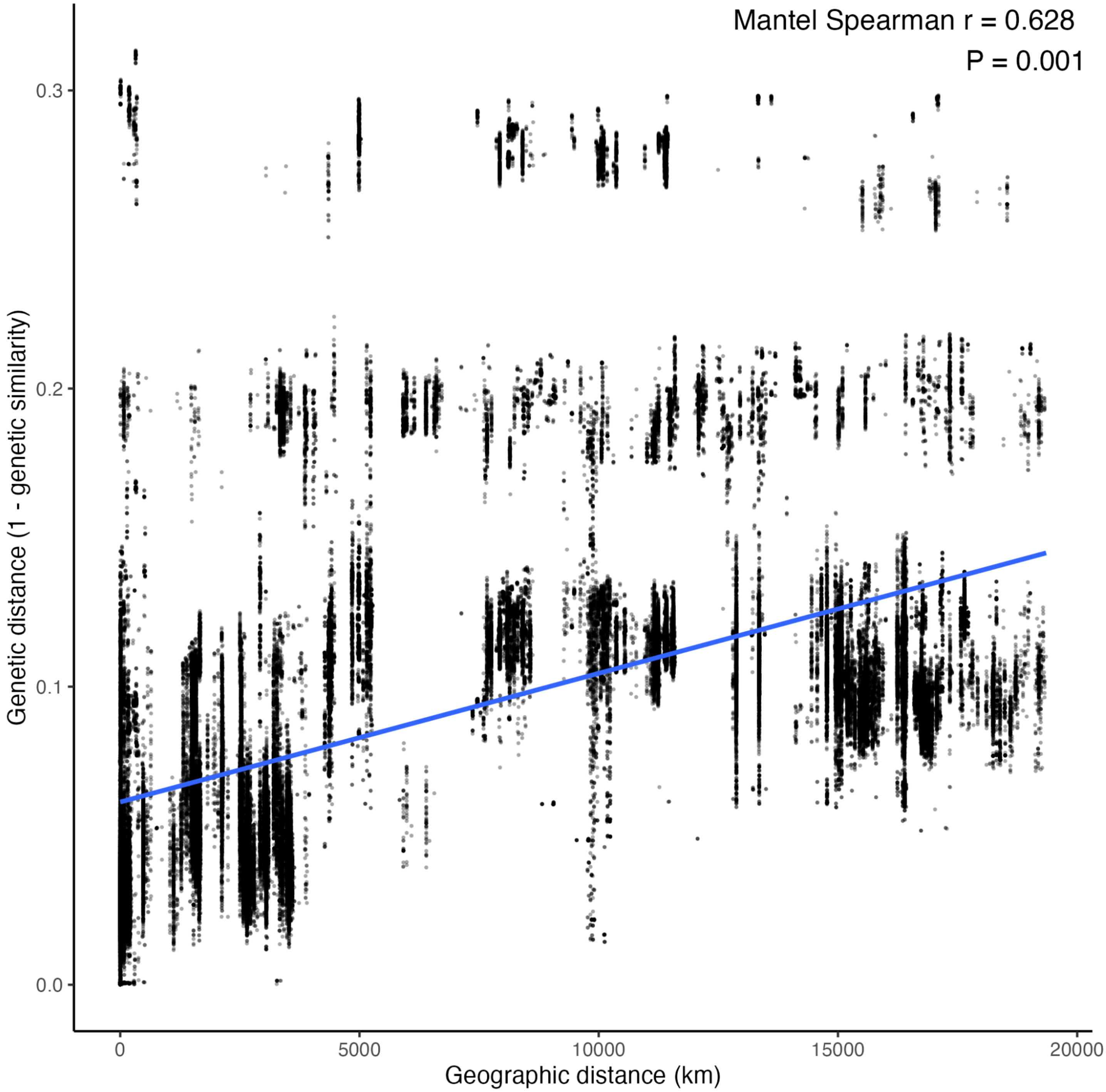


**Figure S12: Isolation by distance across global isotypes.** Each point represents a pairwise comparison between two isotypes. The blue line shows the linear fit. The correlation was assessed using a Mantel test based on Spearman's rank correlation (999 permutations).

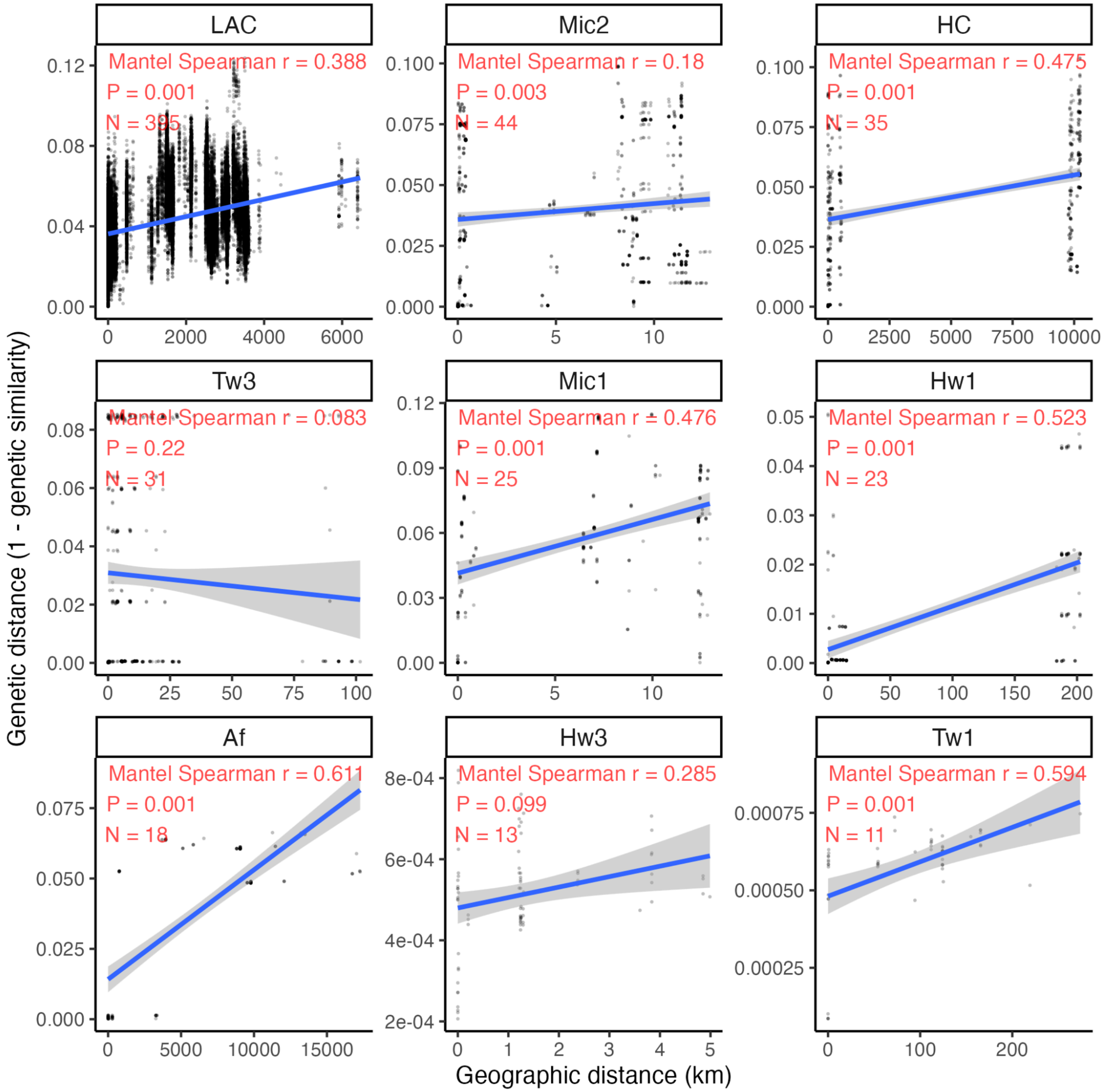


**Figure S13: Isolation by distance within relatedness groups.** Only relatedness groups with more than 10 isotypes were included. Each point represents a pairwise comparison between two isotypes. The blue line shows the linear fit. The correlation was assessed using a Mantel test based on Spearman's rank correlation (999 permutations).

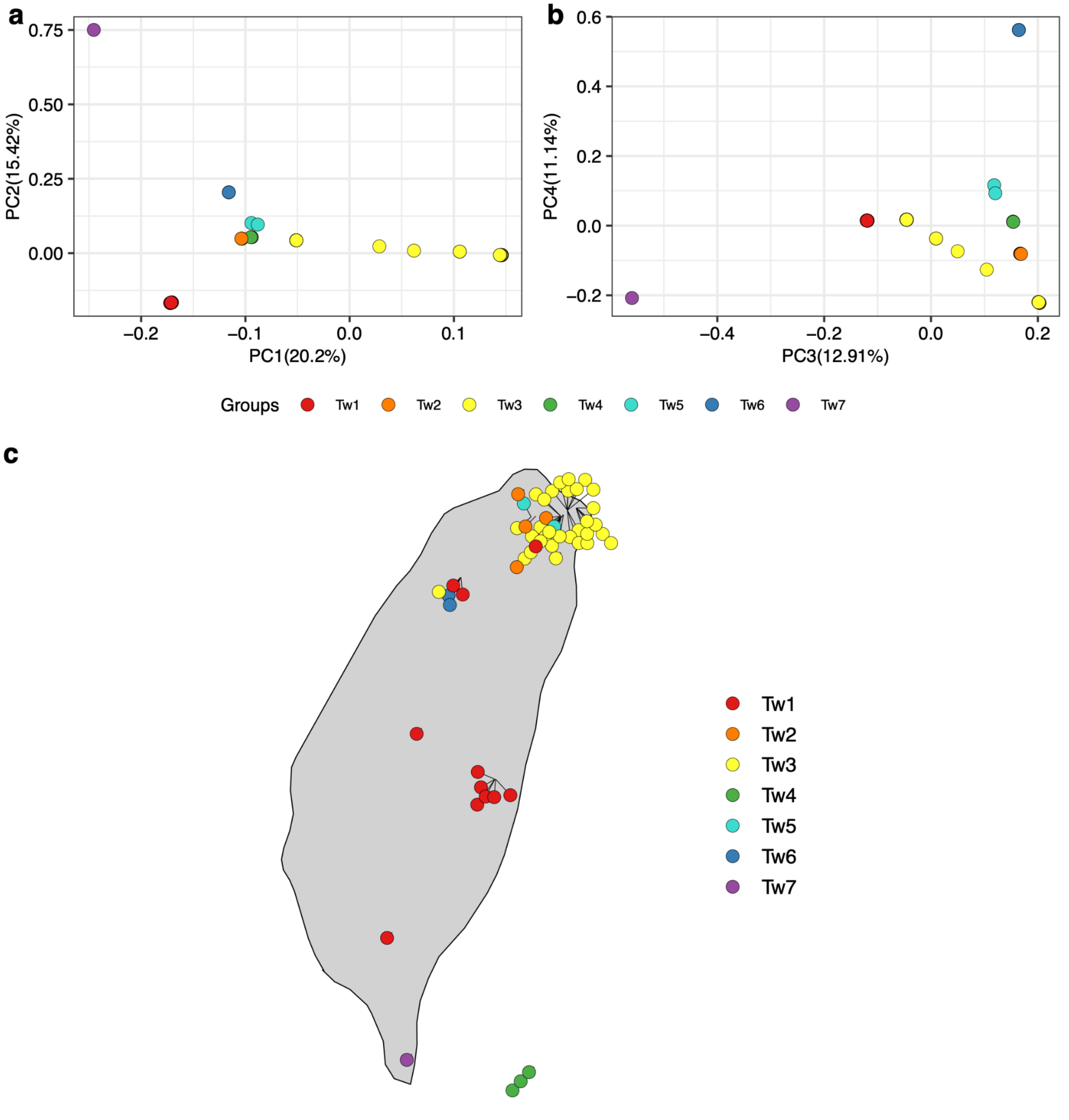


**Figure S14: Distribution of relatedness groups in Taiwan. a,b,** Principal Component Analysis (PCA) of *C. tropicalis* isotype reference strains in Taiwan. **c.** Sampling map of isotype reference strains in Taiwan. Colors in each panel represent the relatedness groups of each strain.

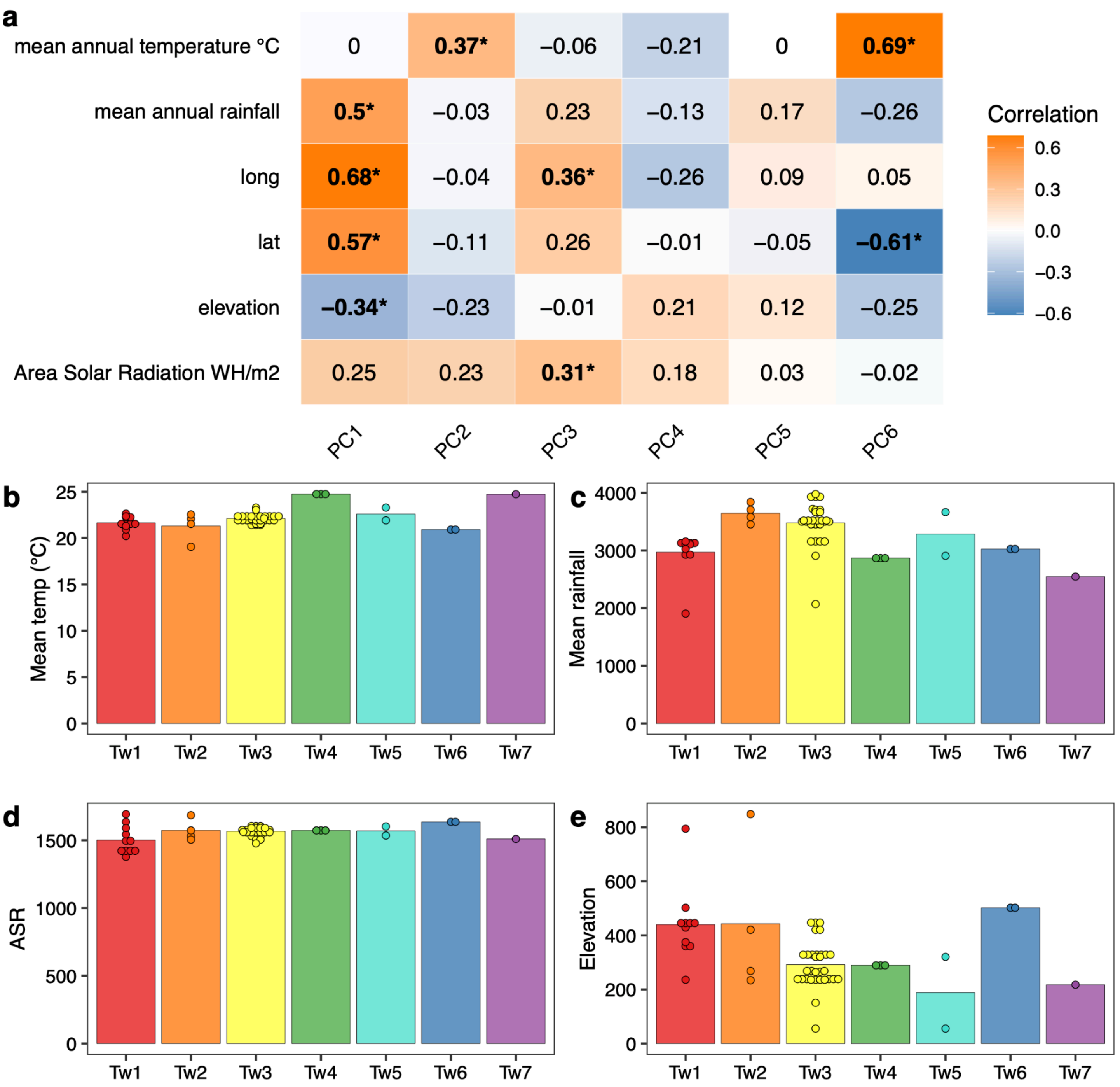


**Figure S15: Environmental parameters of relatedness groups in Taiwan. a,** Correlations between environmental parameters and eigen values of *C. tropicalis* isotype reference strains in Taiwan. **b-e**. Bar plots are shown by relatedness group assignments for different environmental parameters. The asterisk symbol in panel A indicates a significant correlation ($p<0.05$ after Bonferroni correction, Pearson's correlation). Colors in each pane of B-E represent the relatedness groups. ASR stands for Area Solar Radiation (see Materials and Methods).

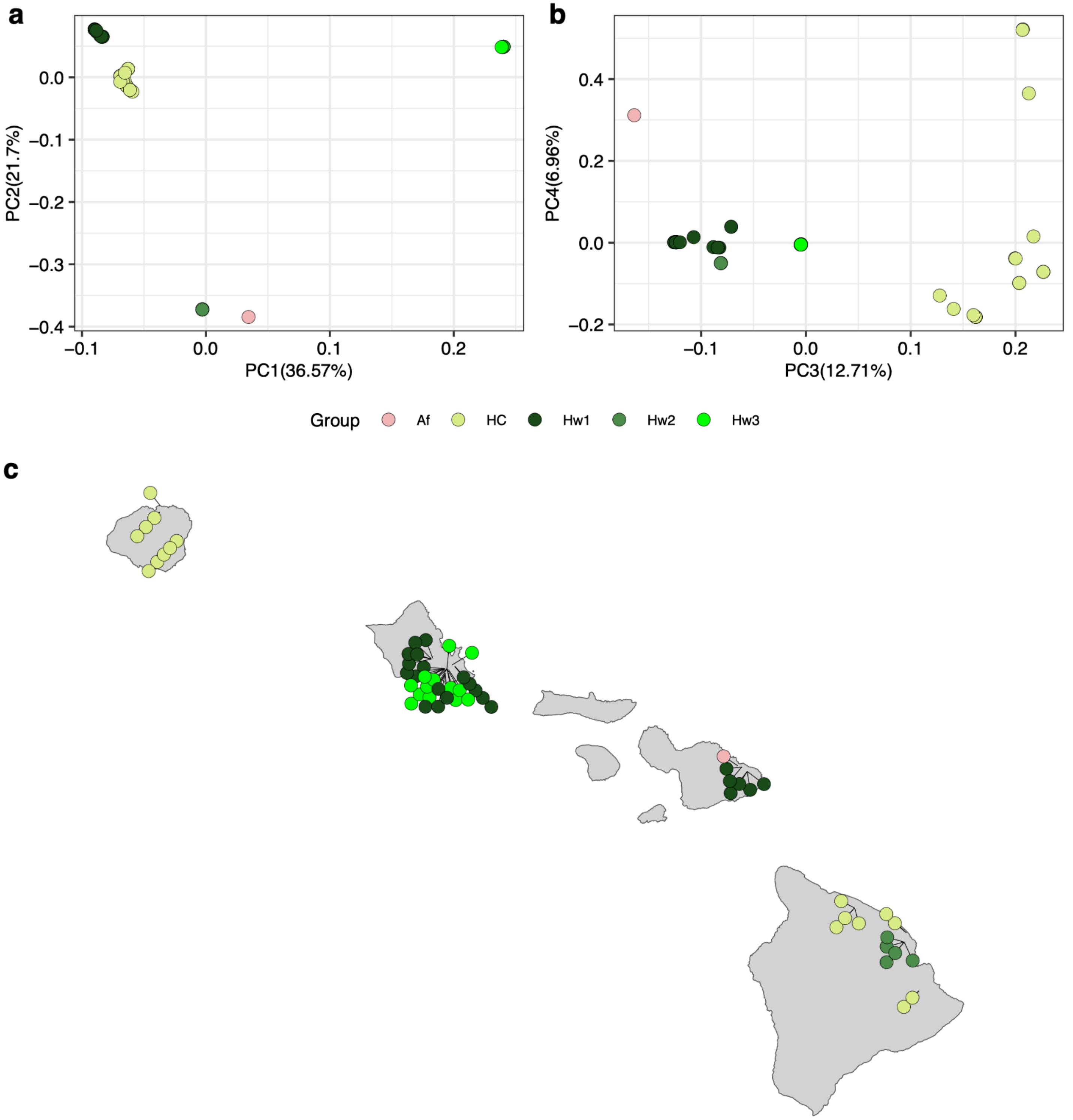


**Figure S16: Distribution of relatedness groups across the Hawaiian Islands. a,b,** Principal Component Analysis (PCA) of *C. tropicalis* isotype reference strains from the Hawaiian Islands. **c.** Sampling map of isotype reference strains. Colors in each panel represent the relatedness groups of each strain.

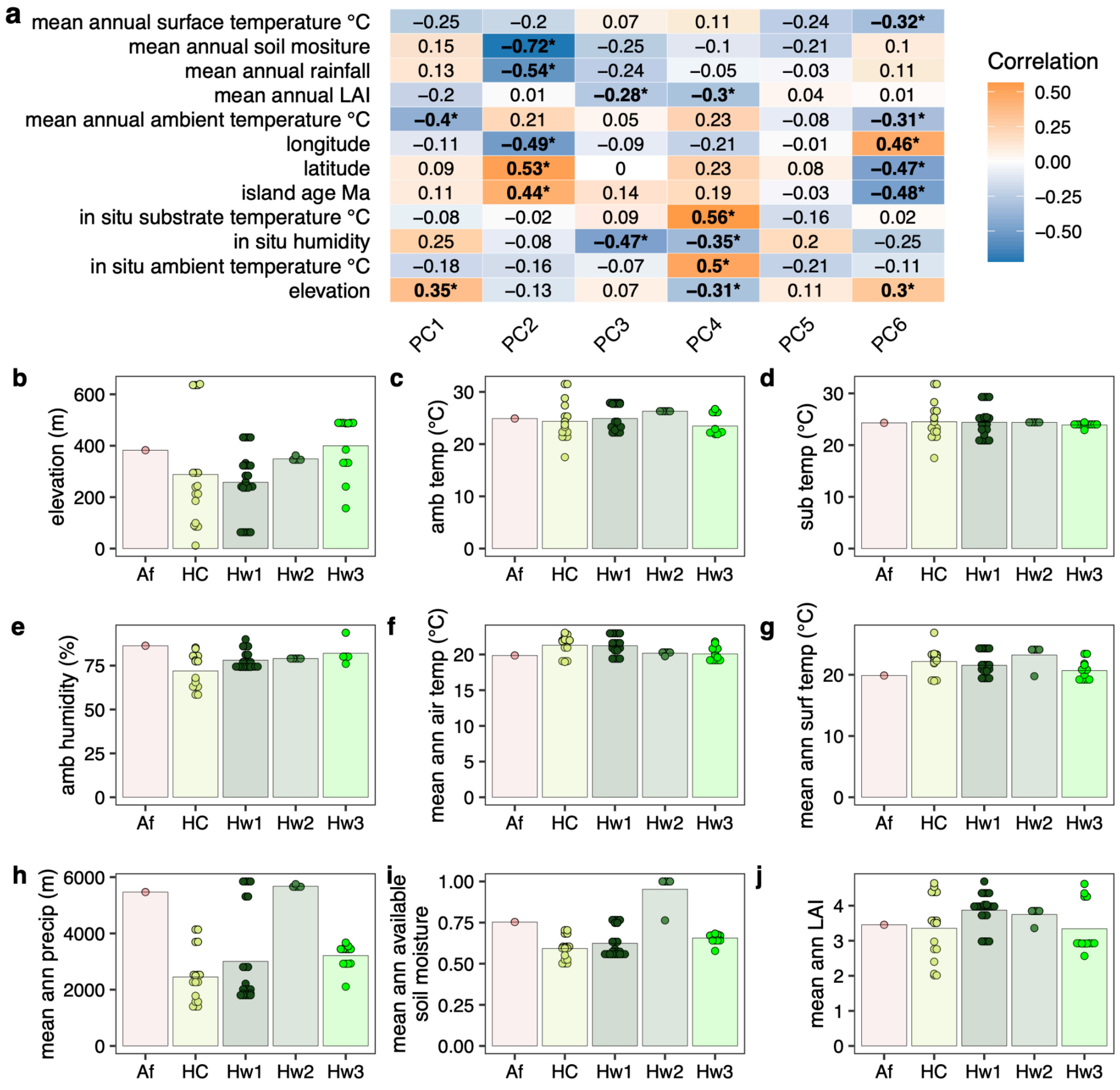


| | PC1 | PC2 | PC3 | PC4 | PC5 | PC6 |
|---|---|---|---|---|---|---|
| mean annual surface temperature °C | −0.25 | −0.2 | 0.07 | 0.11 | −0.24 | **−0.32*** |
| mean annual soil mositure | 0.15 | **−0.72*** | −0.25 | −0.1 | −0.21 | 0.1 |
| mean annual rainfall | 0.13 | **−0.54*** | −0.24 | −0.05 | −0.03 | 0.11 |
| mean annual LAI | −0.2 | 0.01 | **−0.28*** | **−0.3*** | 0.04 | 0.01 |
| mean annual ambient temperature °C | **−0.4*** | 0.21 | 0.05 | 0.23 | −0.08 | **−0.31*** |
| longitude | −0.11 | **−0.49*** | −0.09 | −0.21 | −0.01 | **0.46*** |
| latitude | 0.09 | **0.53*** | 0 | 0.23 | 0.08 | **−0.47*** |
| island age Ma | 0.11 | **0.44*** | 0.14 | 0.19 | −0.03 | **−0.48*** |
| in situ substrate temperature °C | −0.08 | −0.02 | 0.09 | **0.56*** | −0.16 | 0.02 |
| in situ humidity | 0.25 | −0.08 | **−0.47*** | **−0.35*** | 0.2 | −0.25 |
| in situ ambient temperature °C | −0.18 | −0.16 | −0.07 | **0.5*** | −0.21 | −0.11 |
| elevation | **0.35*** | −0.13 | 0.07 | **−0.31*** | 0.11 | **0.3*** |



**Figure S17: Environmental parameters of relatedness groups across the Hawaiian Islands. a,** Correlations between environmental parameters and eigen values of *C. tropicalis* isotype reference strains across the Hawaiian Islands. b-j. Bar plots are shown by relatedness group assignments for different environmental parameters. The asterisk symbol in panel A indicates a significant correlation (p<0.05 after Bonferroni correction, Pearson's correlation). Colors in each pane of b-j represent the relatedness groups. LAI stands for leaf area index (see Materials and Methods).

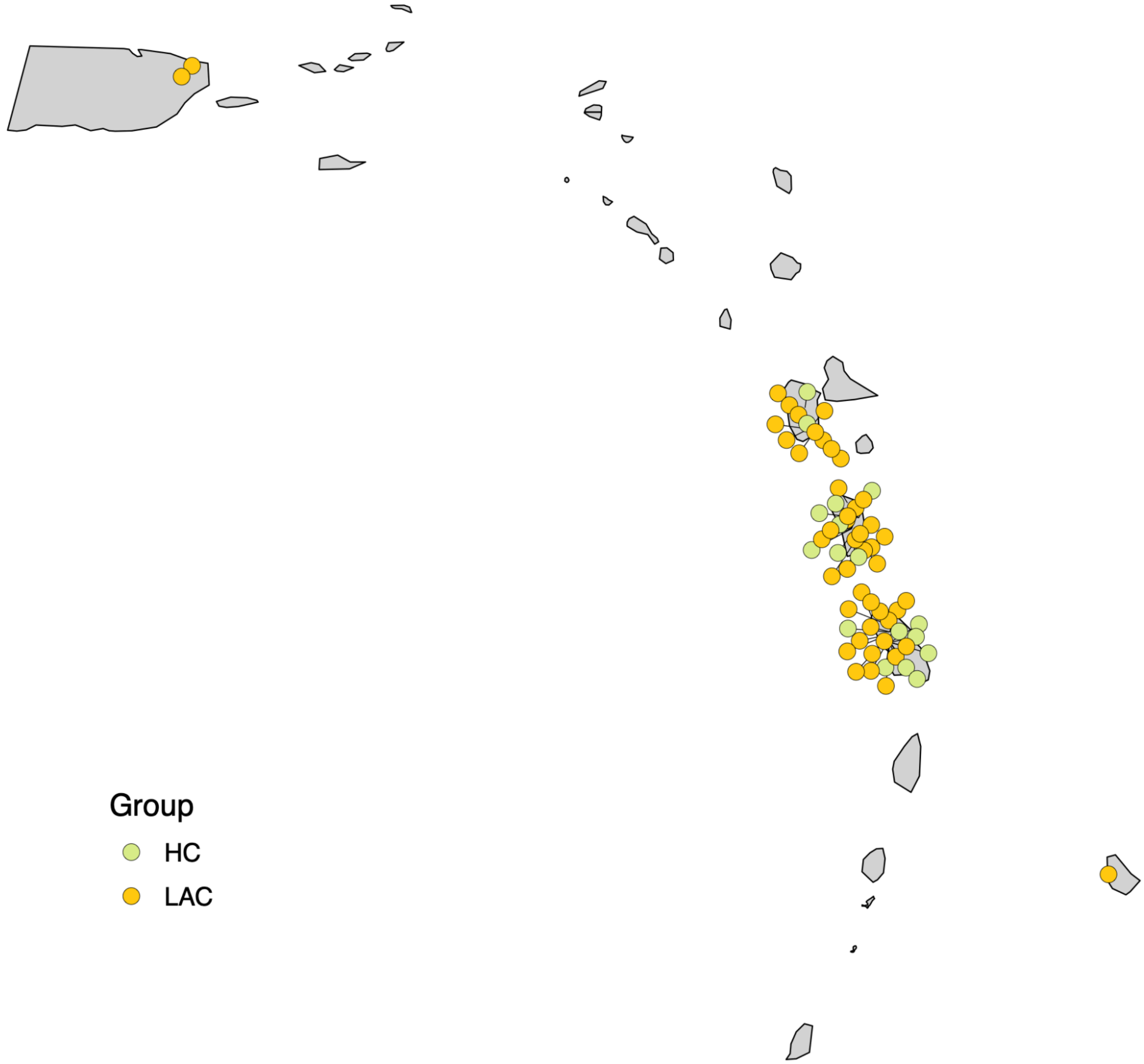


**Figure S18: Distribution of relatedness groups in the Caribbean.** Colors in each panel represent the relatedness groups of each strain.

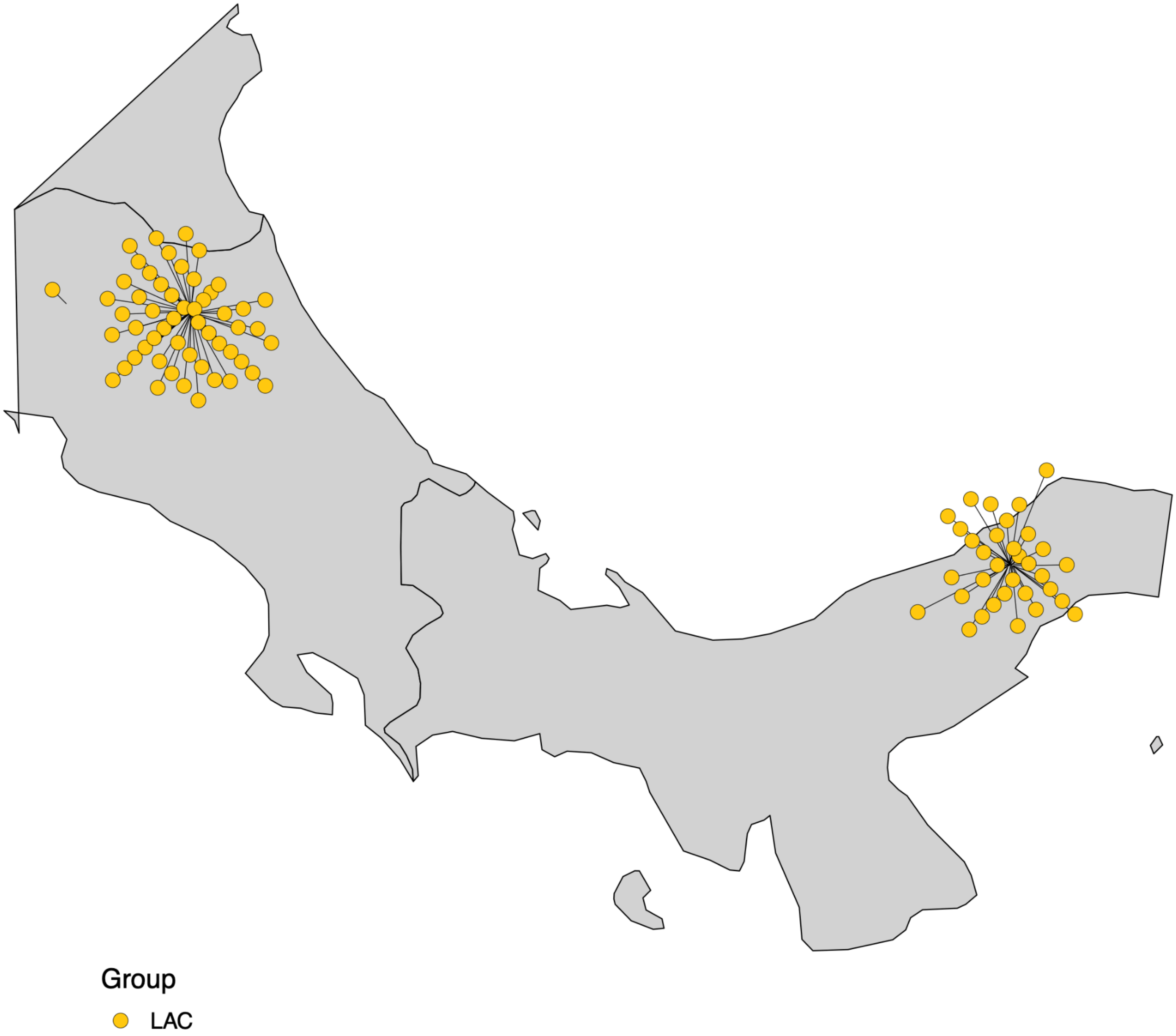


**Figure S19: Distribution of relatedness groups in Central America.** Colors in each panel represent the relatedness groups of each strain.

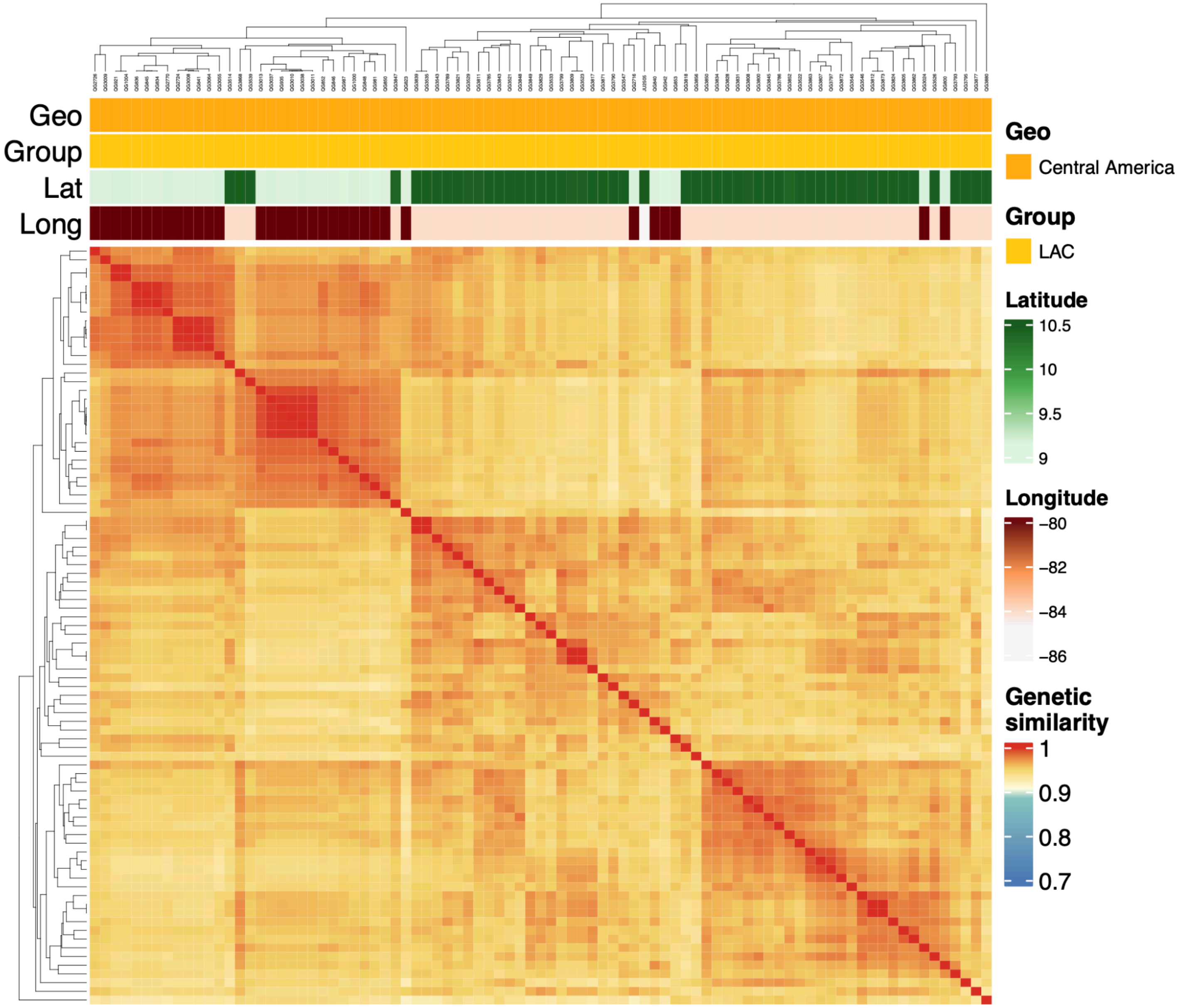


**Figure S20: The genetic similarity of the strains in Central America.** Pairwise genetic similarity was estimated by the proportion of identical alleles across all identified SNVs. The genetic similarity color gradient has breaks specified at 0.8853456, 0.9068701, and 0.9537771, which is consistent with the 25th, 50th, and 75th percentile of the distribution of pairwise estimates for all 622 global isotypes as shown in the Fig. 2. In order from top to bottom, bands show the geographic region, relatedness group, latitude, and longitude of the isolation site of each strain.

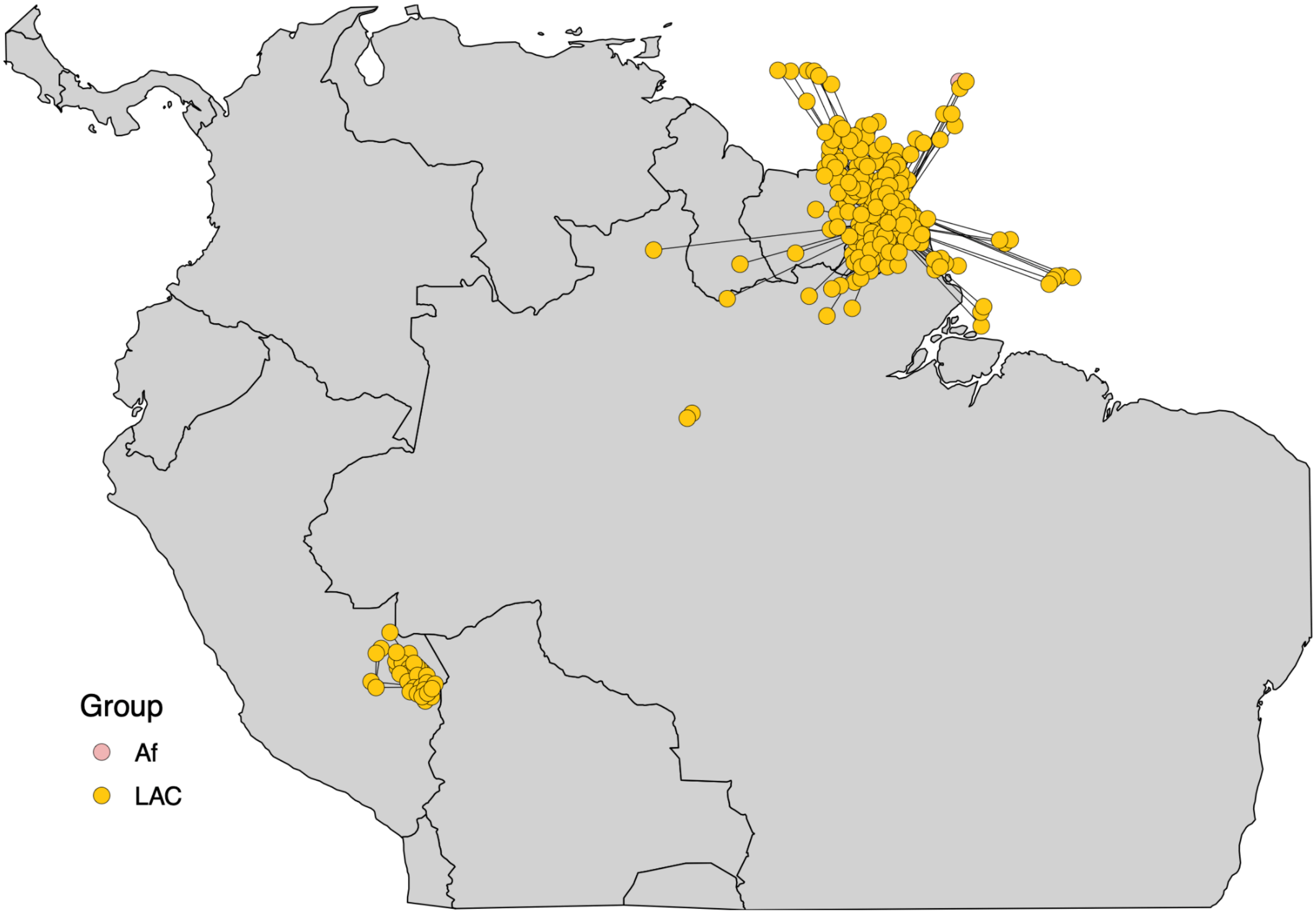


**Figure S21: Distribution of relatedness groups in South America.** Colors in each panel represent the relatedness groups of each strain.

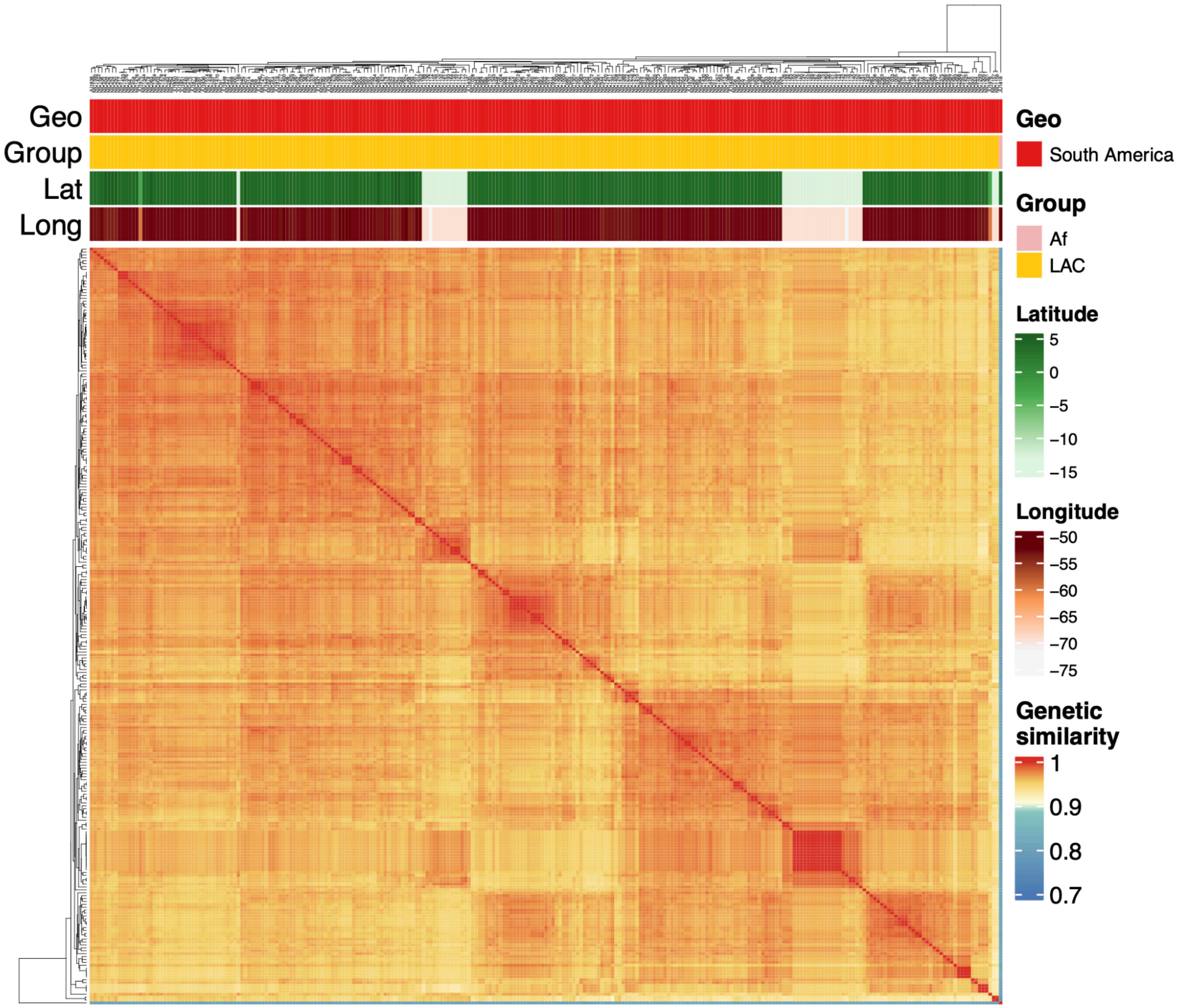


**Figure S22: The genetic similarity of the strains in South America.** Pairwise genetic similarity was estimated by the proportion of identical alleles across all identified SNVs. The genetic similarity color gradient has breaks specified at 0.8853456, 0.9068701, and 0.9537771, which is consistent with the 25th, 50th, and 75th percentile of the distribution of pairwise estimates for all 622 global isotypes as shown in the Fig. 2. In order from top to bottom, bands show the geographic region, relatedness group, latitude, and longitude of the isolation site of each strain.

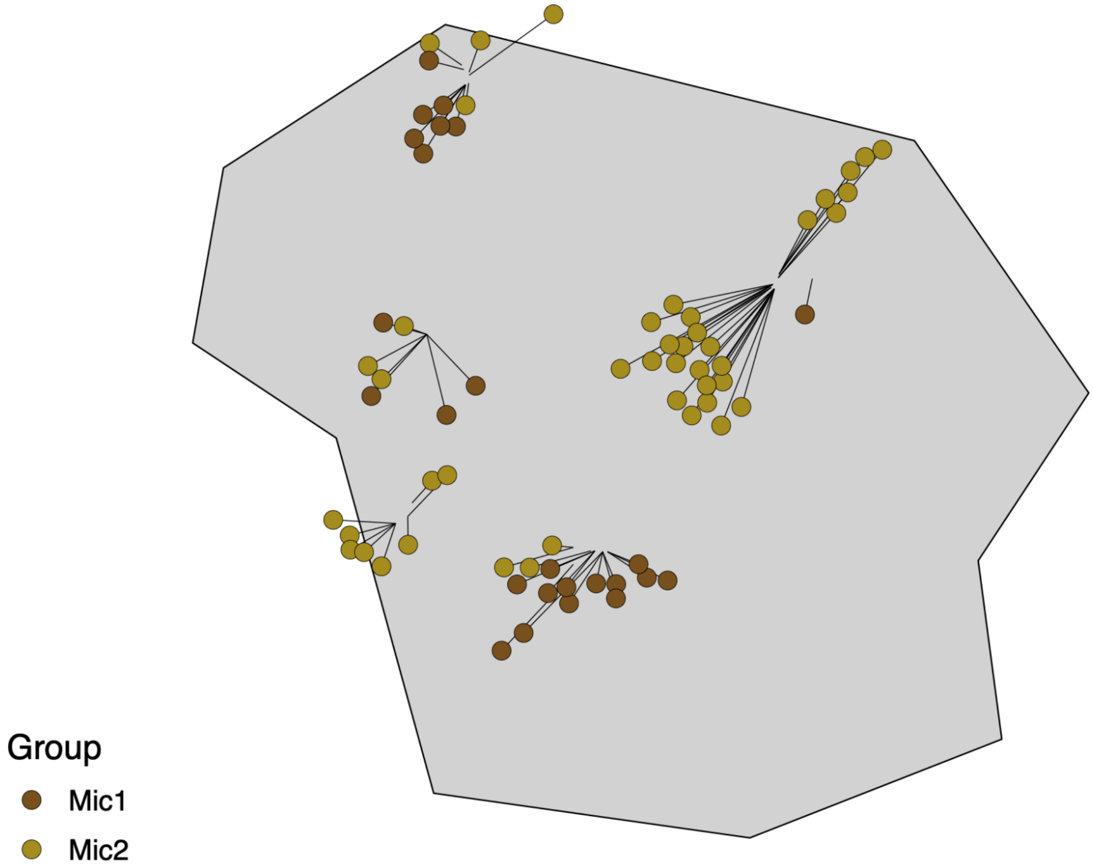


**Figure S23: Distribution of relatedness groups in Micronesia.** Colors in each panel represent the relatedness groups of each strain.

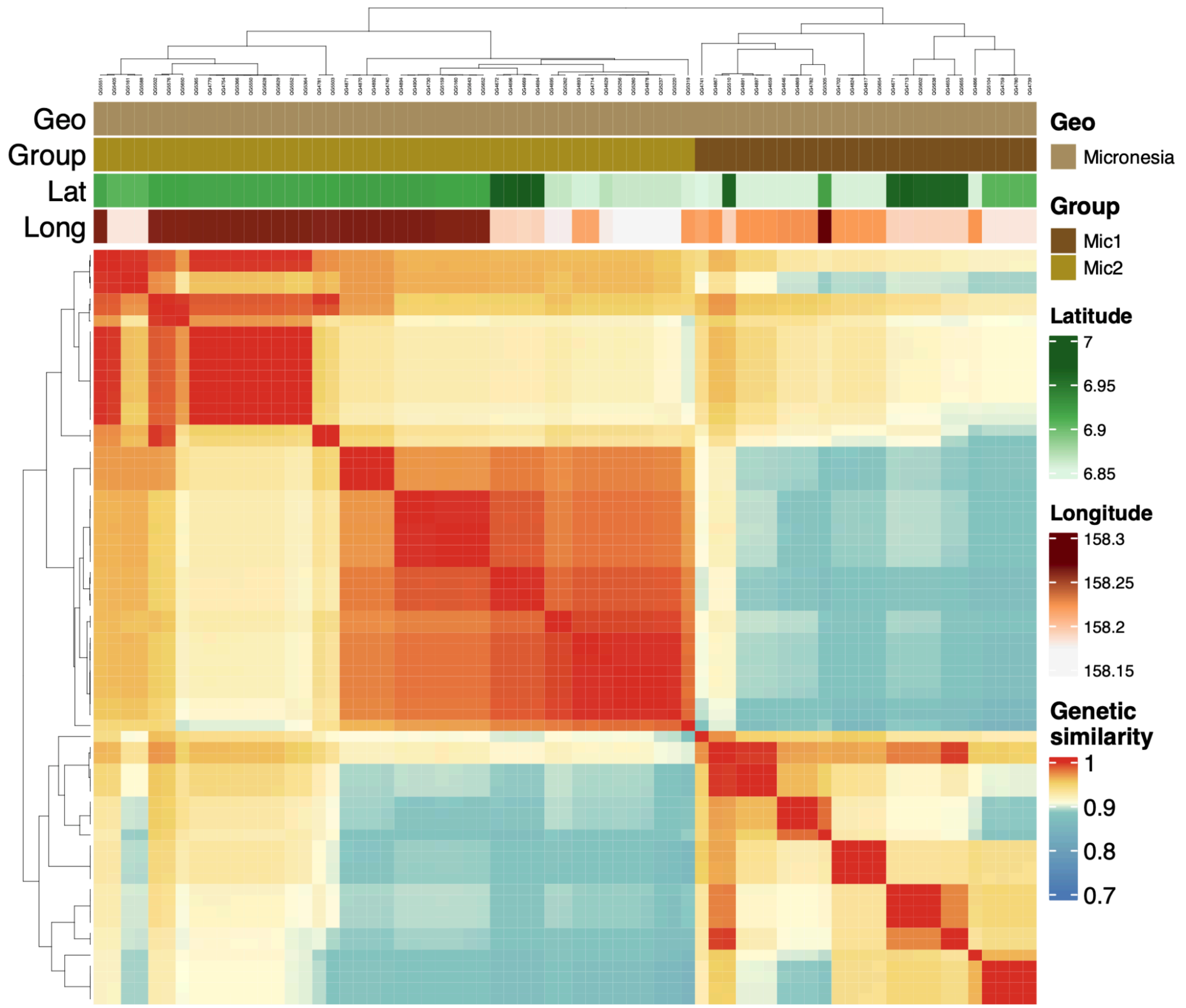


**Figure S24: The genetic similarity of the strains in Micronesia.** Pairwise genetic similarity was estimated by the proportion of identical alleles across all identified SNVs. The genetic similarity color gradient has breaks specified at 0.8853456, 0.9068701, and 0.9537771, which is consistent with the 25th, 50th, and 75th percentile of the distribution of pairwise estimates for all 622 global isotypes as shown in the Fig. 2. In order from top to bottom, bands show the geographic region, relatedness group, latitude, and longitude of the isolation site of each strain.

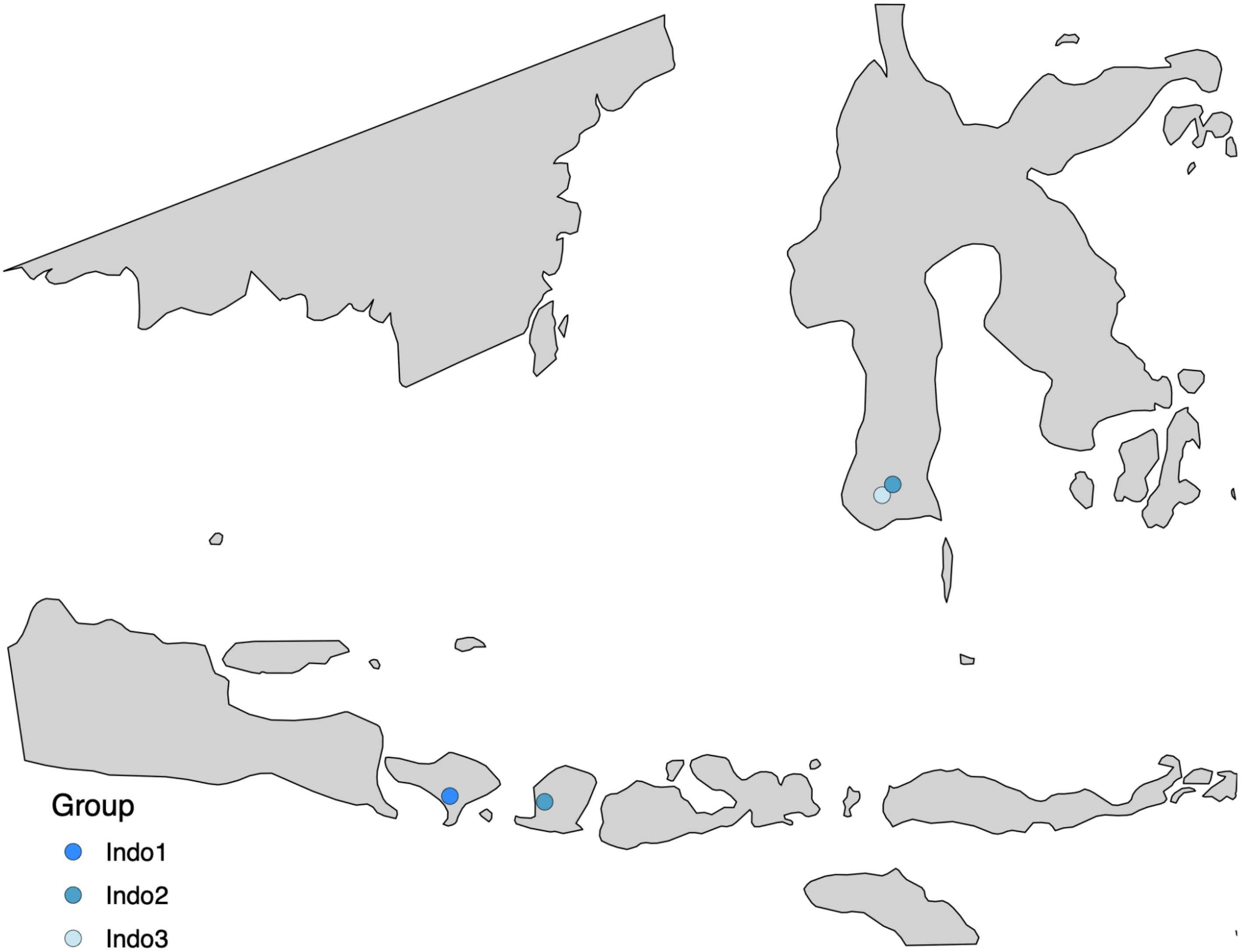


**Figure S25: Distribution of relatedness groups in the Malay Archipelago.** Colors in each panel represent the relatedness groups of each strain.

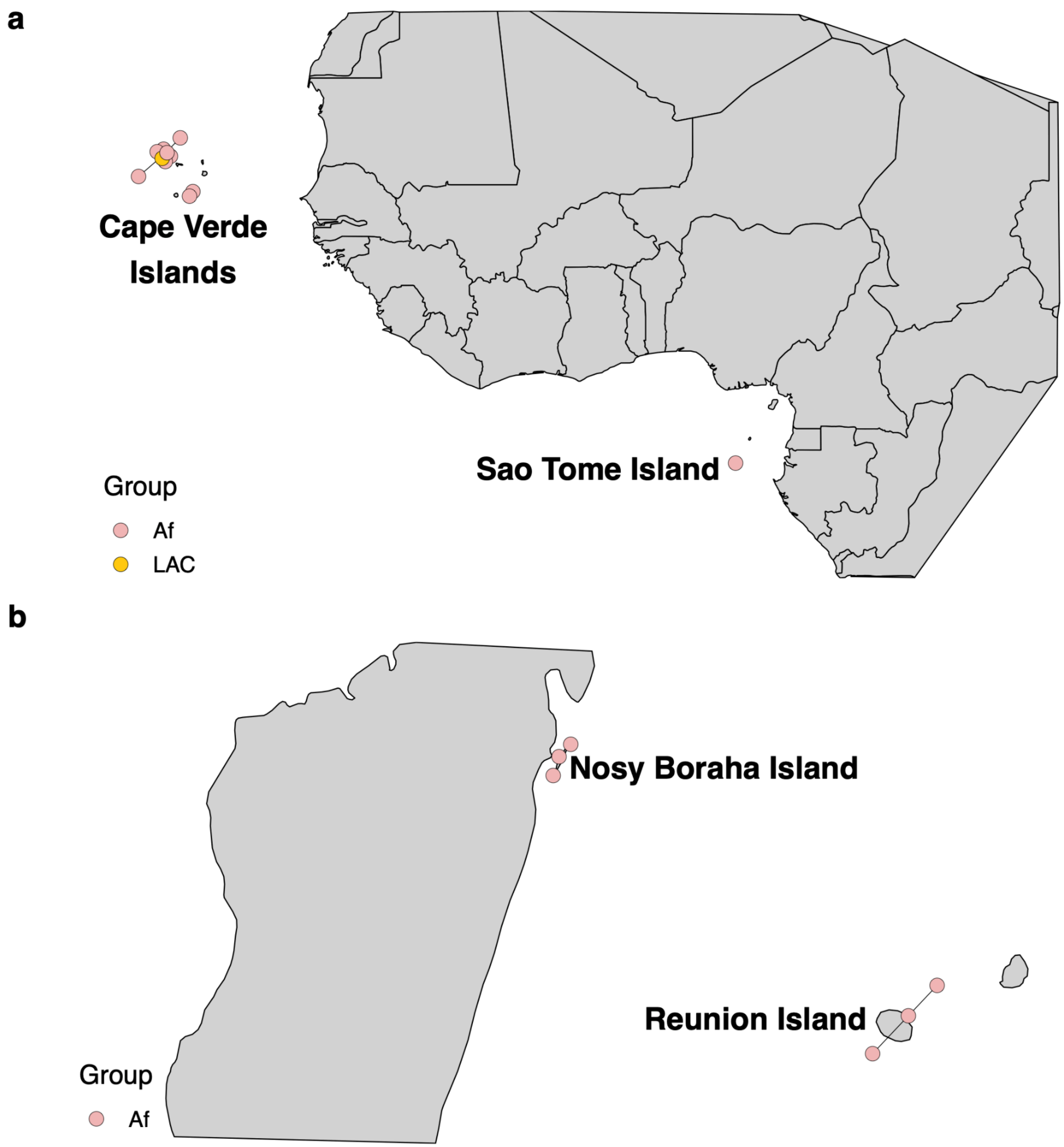


**Figure S26: Distribution of relatedness groups in Africa. a,** Distribution of relatedness groups on the islands off the west coast of Africa. **b,** Distribution of relatedness groups on the islands off the east coast of Africa. Colors in each panel represent the relatedness groups of each strain.

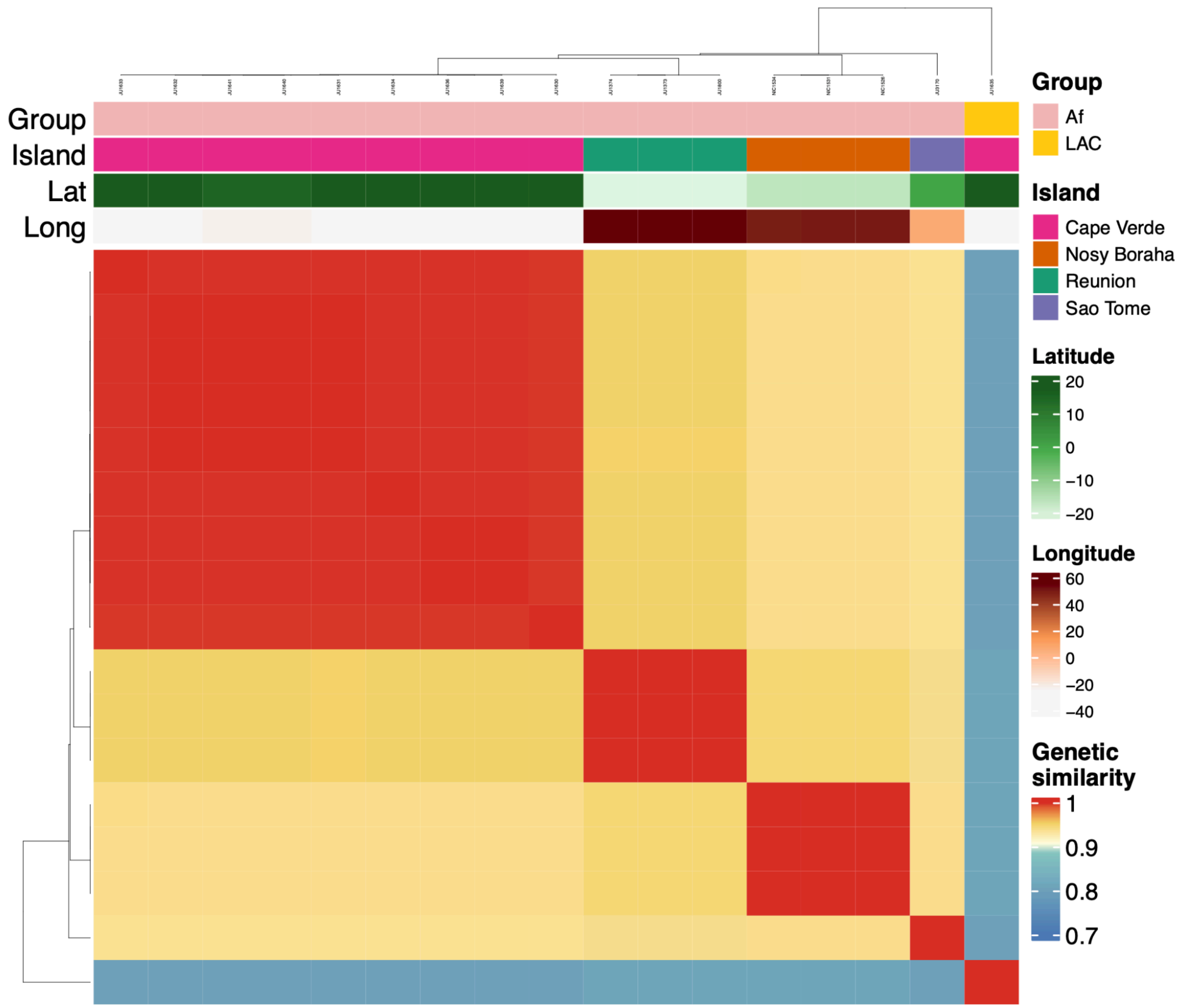


**Figure S27: The genetic similarity of the strains in Africa.** Pairwise genetic similarity was estimated by the proportion of identical alleles across all identified SNVs. The genetic similarity color gradient has breaks specified at 0.8853456, 0.9068701, and 0.9537771, which is consistent with the 25th, 50th, and 75th percentile of the distribution of pairwise estimates for all 622 global isotypes as shown in the Fig. 2. In order from top to bottom, bands show the relatedness group, island, latitude, and longitude of the isolation site of each strain.

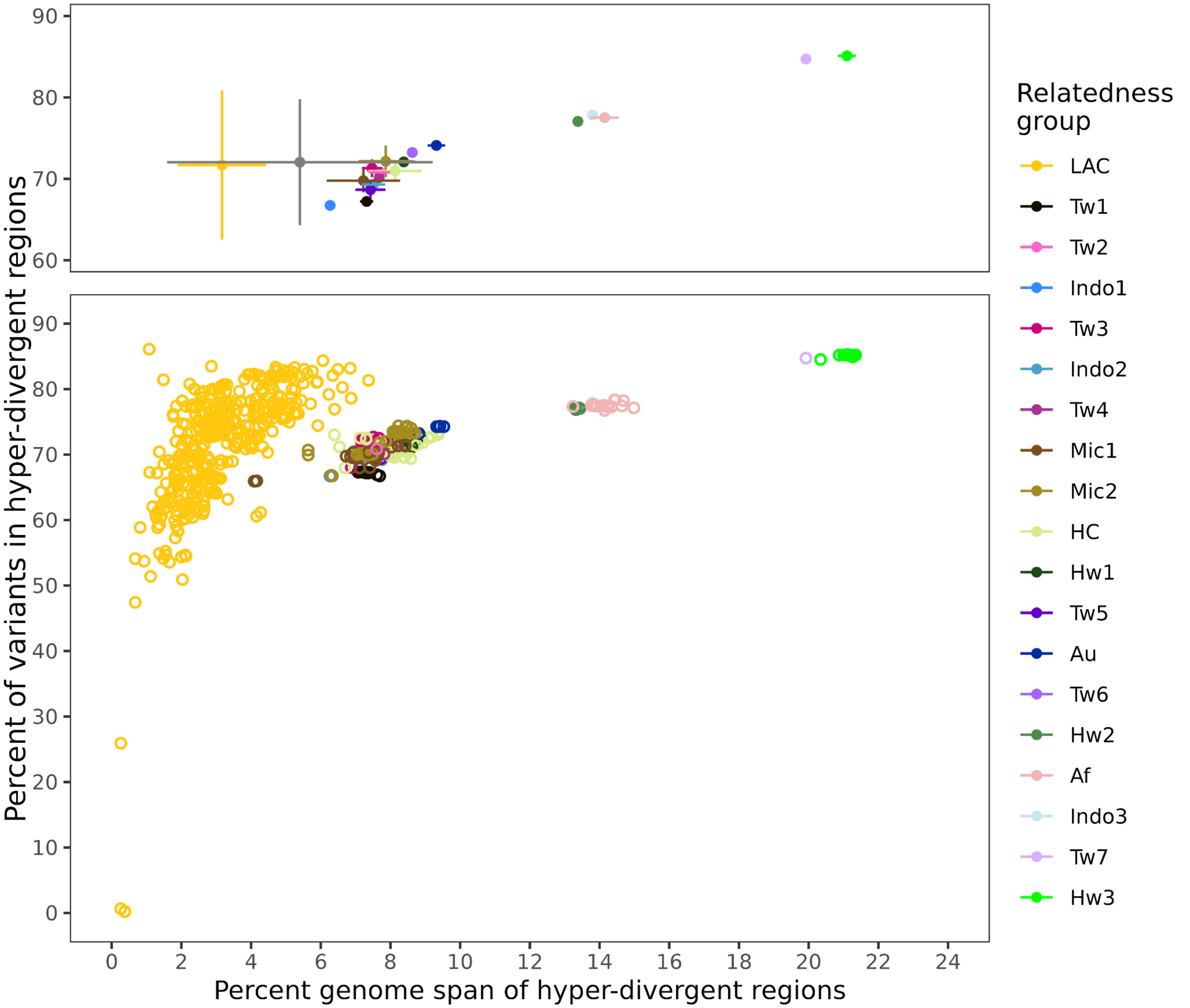


**Figure S28: Hyper-divergent regions (HDRs) contain the vast majority of genetic variation within a small proportion of the genome.** Each point in the bottom panel represents an isotype strain colored by relatedness group with the percent of variants within HDRs shown along the y-axis and the percent of genome spanned by HDRs along the x-axis. The top panel shows mean and standard deviations by relatedness group for the same estimates.

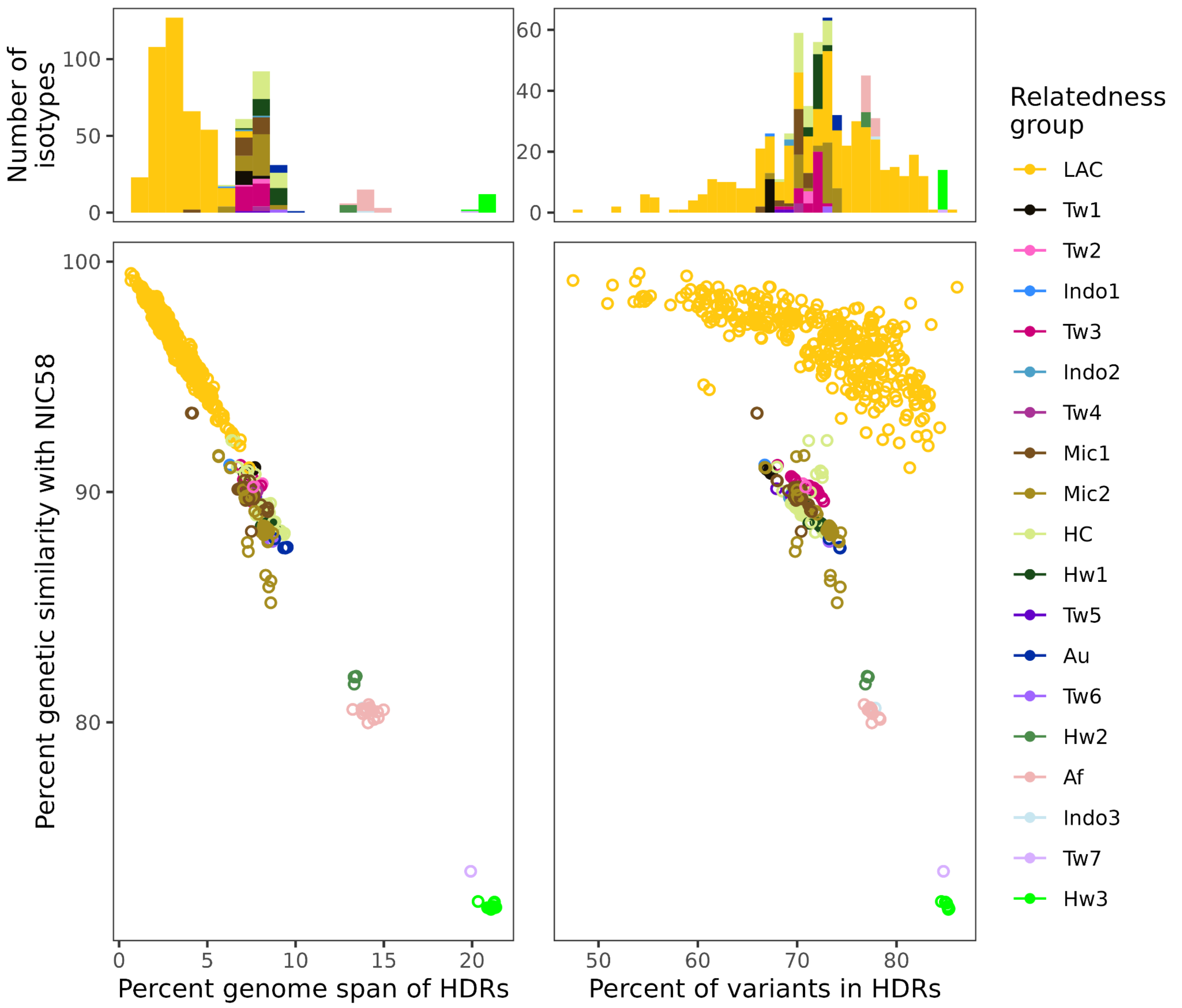


**Figure S29: The extent of HDRs decreases linearly with genetic similarity to the NIC58 reference genome.** Scatter plots show the percent genetic similarity between individual isotypes and the NIC58 reference genome relative to the percent of the genome spanned by HDRs (bottom left) or the percent of variants in HDRs (bottom right). Each point represents an isotype colored by their assigned relatedness group. Bar plots in each top panel show the number of isotypes for each relatedness group across 1% bins of hyper-divergent genome span (top left) or percent of variants in HDRs (top right).

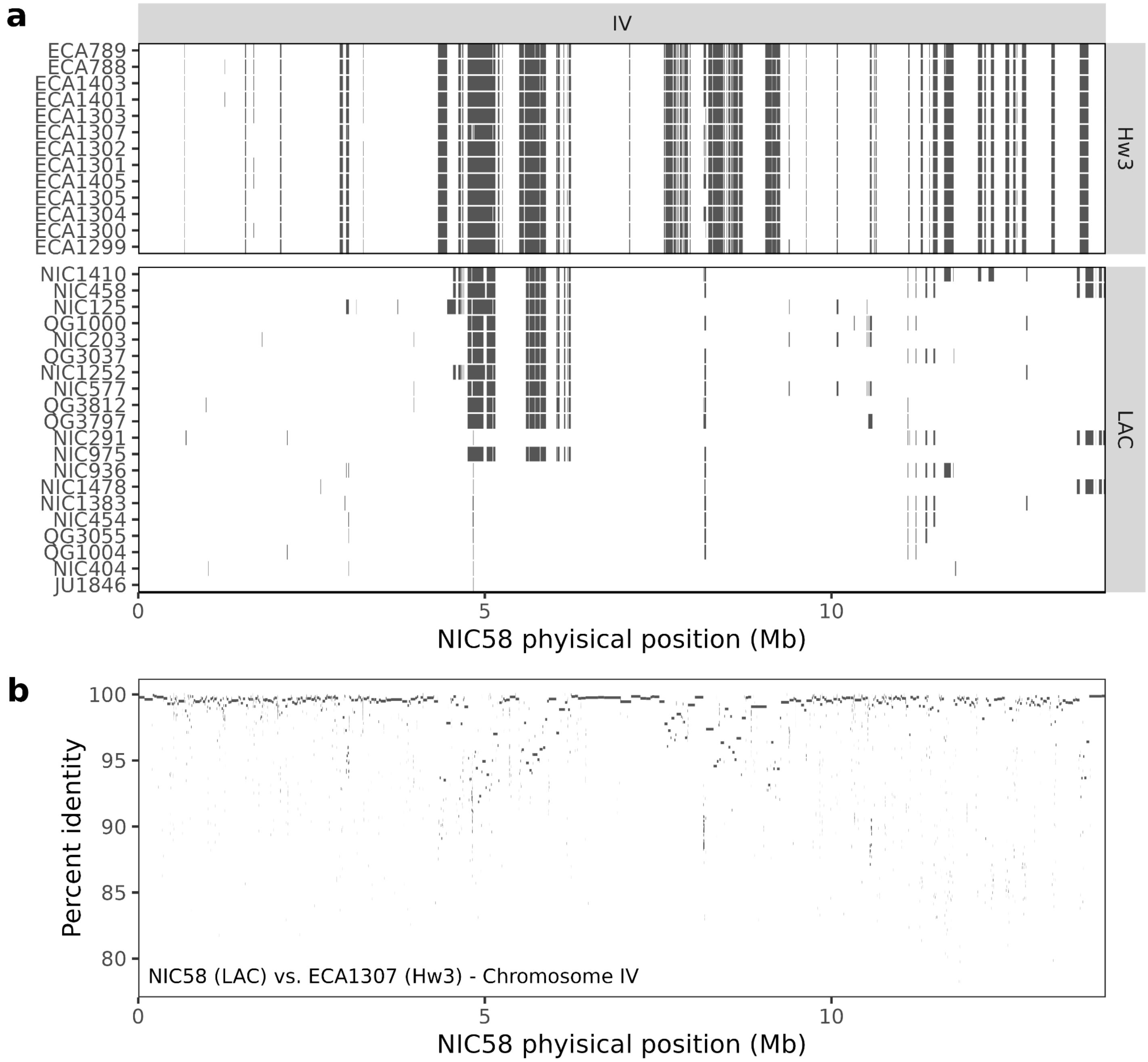


**Figure S30: Divergent Hawaiian relatedness group carries private and shared HDRs relative to the reference LAC relatedness group. a,** HDRs identified across chromosome IV for all isotype strains assigned to the Hawaii3 (Hw3, top panel) relatedness group and a subsample of 20 isotype strains assigned to the Latin America and Caribbean (LAC, bottom panel) relatedness group. The NIC58 genome coordinates are shown on the x-axis, and each row represents an individual isotype genome with HDRs marked as black boxes. **b,** genome alignments between representative genomes of both the Hw3 (ECA1307) and LAC (NIC58) relatedness groups. The identity of each ECA1307 alignment relative to NIC58 is shown on the y-axis. Alignments alternate from low-identity (<97%) at HDRs to high-identity (>97%) at non-HDRs.

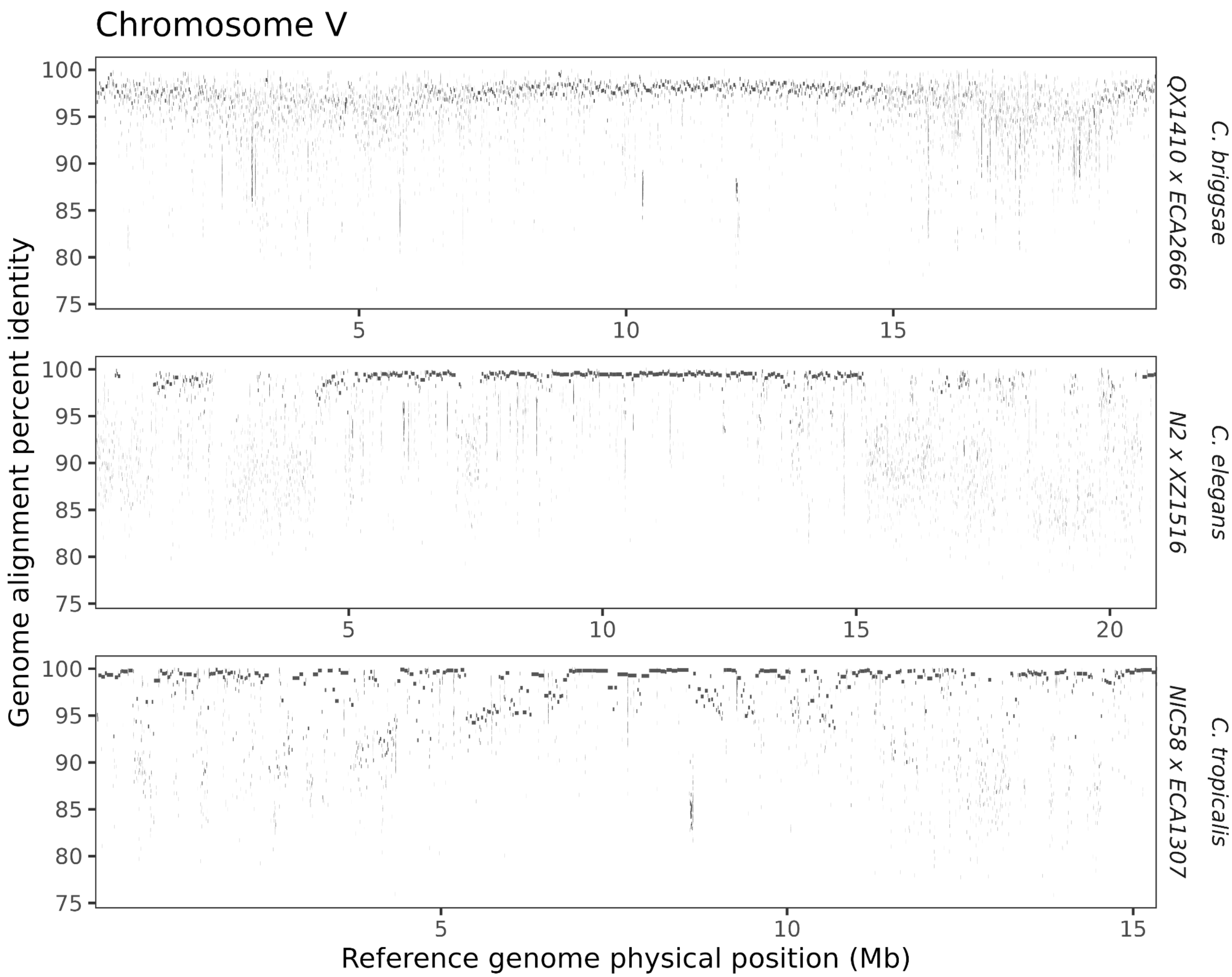


**Figure S31: Genome alignments between highly divergent isotype pairs in all three selfing species.** Examples of genome alignments in chromosome V between highly diverged isotype pairs in *C. briggase*, *C. elegans*, and *C. tropicalis*. The genomic coordinates of the reference genome of each species (*C. briggsae*, QX1410; *C. elegans*, N2; *C. tropicalis*, NIC58) are shown on the x-axis of each panel, and the identity of the genome alignments from each divergent isotype relative to each respective species reference isotype is shown on the y-axis.

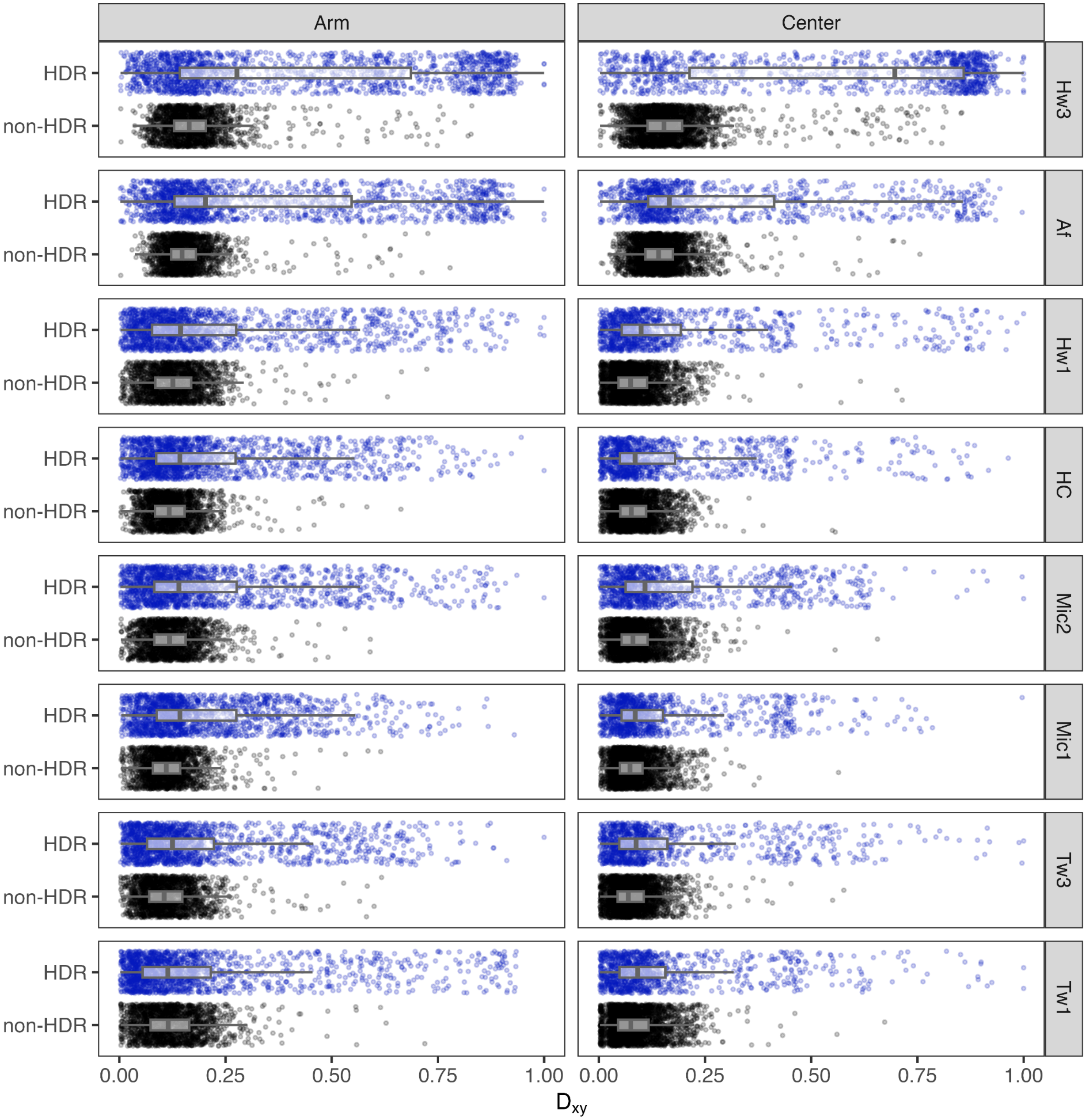


**Figure S32: HDRs display higher or no difference in absolute divergence across different relatedness group comparisons.** Boxplots show the Dxy across 10 kb genome segments between LAC and other relatedness groups comparing HDRs (blue) and non-HDRs (black) regions.

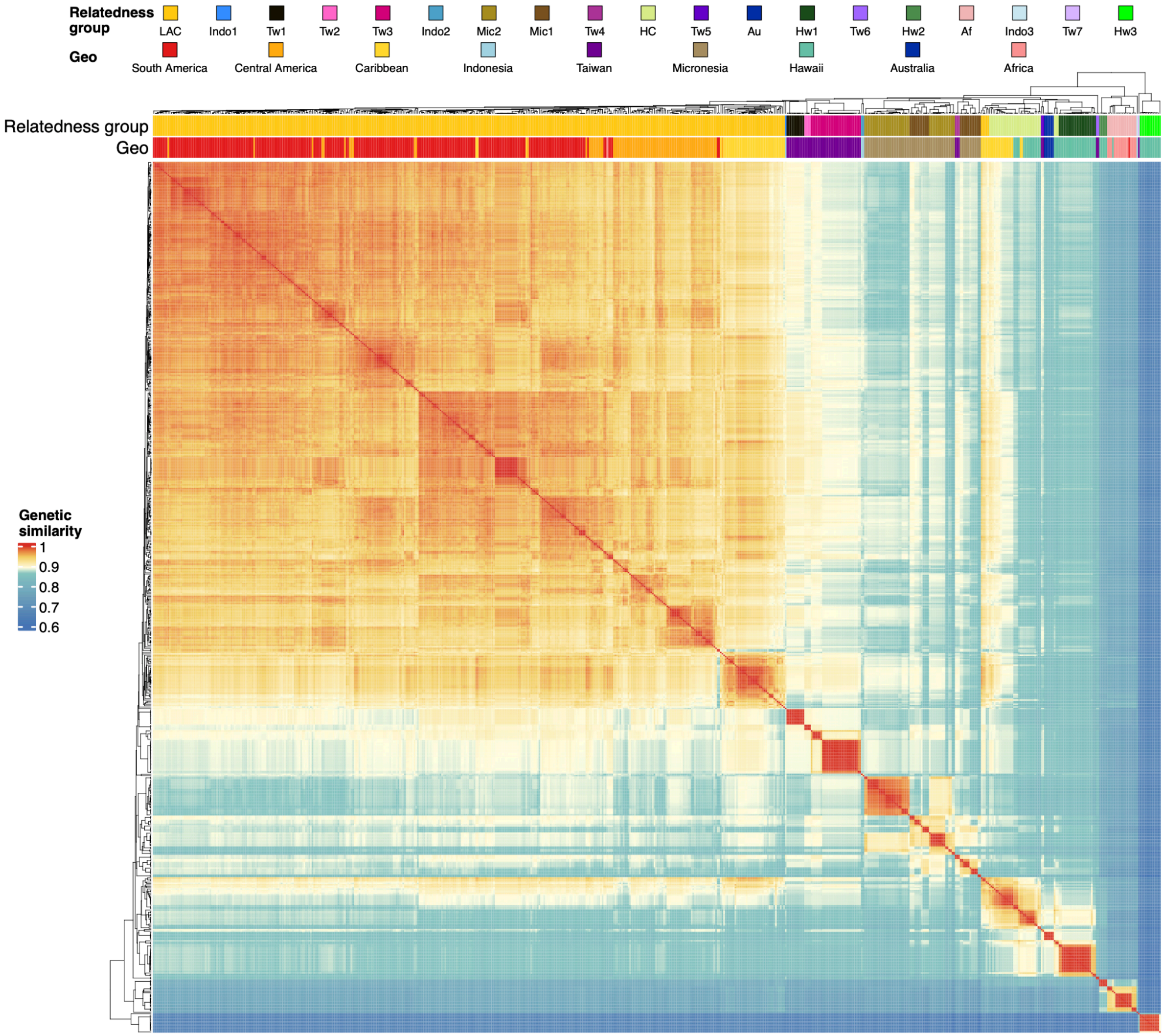


**Figure S33: Pairwise genetic similarity across the HDRs.** Heatmap showing pairwise genetic similarity across the HDRs between all 622 isotype reference strains. Pairwise genetic similarity was estimated by the proportion of identical alleles across all identified SNVs within HDRs among the 622 reference strains. The genetic similarity color gradient has breaks specified at the 25th, 50th, and 75th percentile of the distribution of all pairwise estimates. The top bands show the geographic region of the isolation site of each isotype representative strain and the assigned relatedness group.

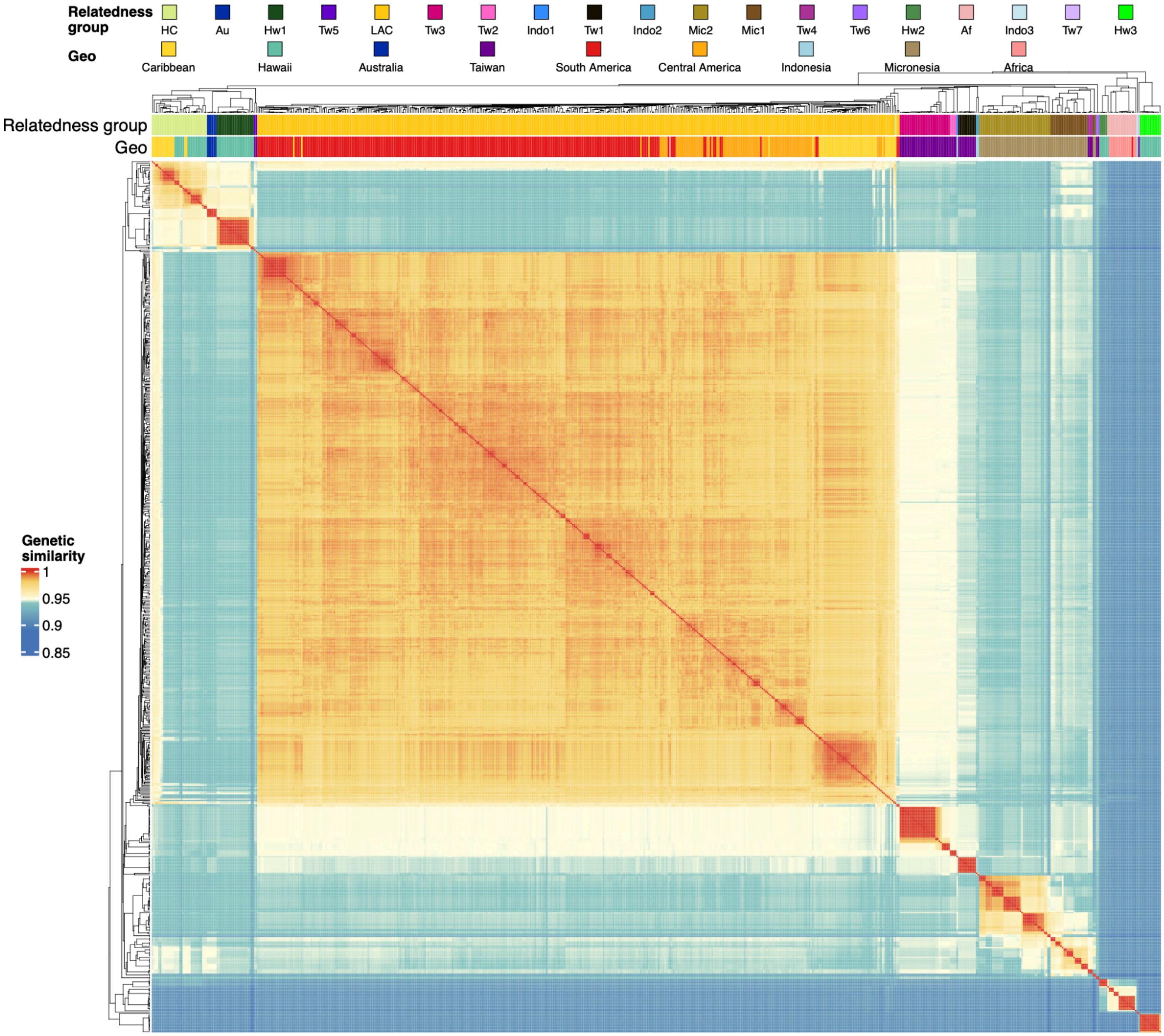


**Figure S34: Pairwise genetic similarity across the non-HDRs.** Heatmap showing pairwise genetic similarity across the non-HDRs between all 622 isotype reference strains. Pairwise genetic similarity was estimated by the proportion of identical alleles across all identified SNVs outside of HDRs among the 622 reference strains. The genetic similarity color gradient has breaks specified at the 25th, 50th, and 75th percentile of the distribution of all pairwise estimates. The top bands show the geographic region of the isolation site of each isotype representative strain and the assigned relatedness group.

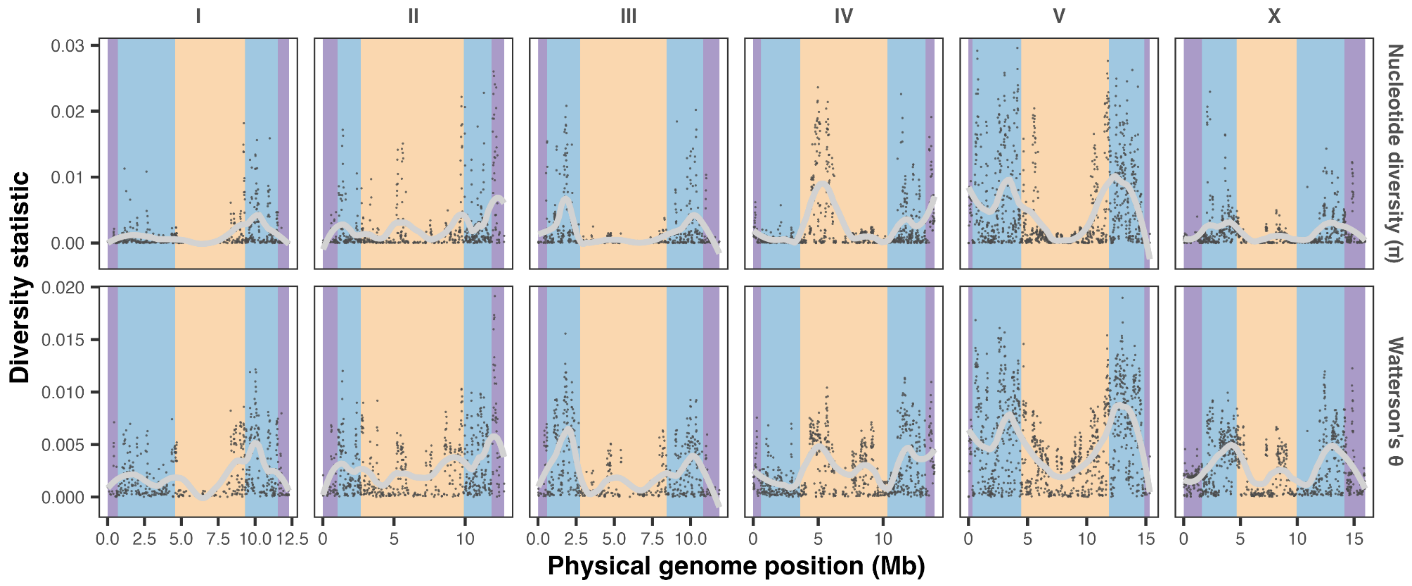


**Figure S35: Diversity statistics of non-HDRs along physical genome positions across the six chromosomes.** Each point represents an estimate of nucleotide diversity ($\pi$ or $\theta_W$) across a 10 kb window. Grey lines are weighted LOESS fits where each local regression used 30% of the data. Colored backgrounds in nucleotide diversity panels delineate the chromosomal domain boundaries (purple, tip; blue, arm; yellow, center).

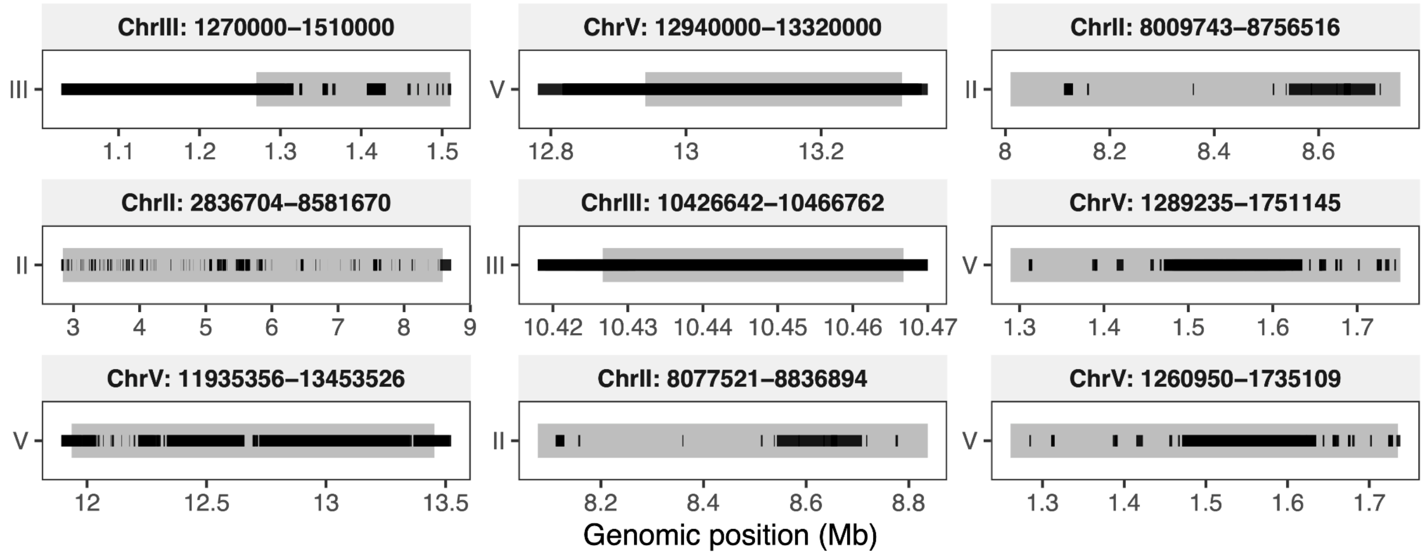


**Figure S36: Overlap between previously described toxin-antidote systems (TAs) and HDRs identified in this study.** The gray regions represent genomic locations of known TA regions, and the black boxes represent the overlapping HDRs identified in this study. Coordinates of all the TA regions are labeled in the title of each panel.

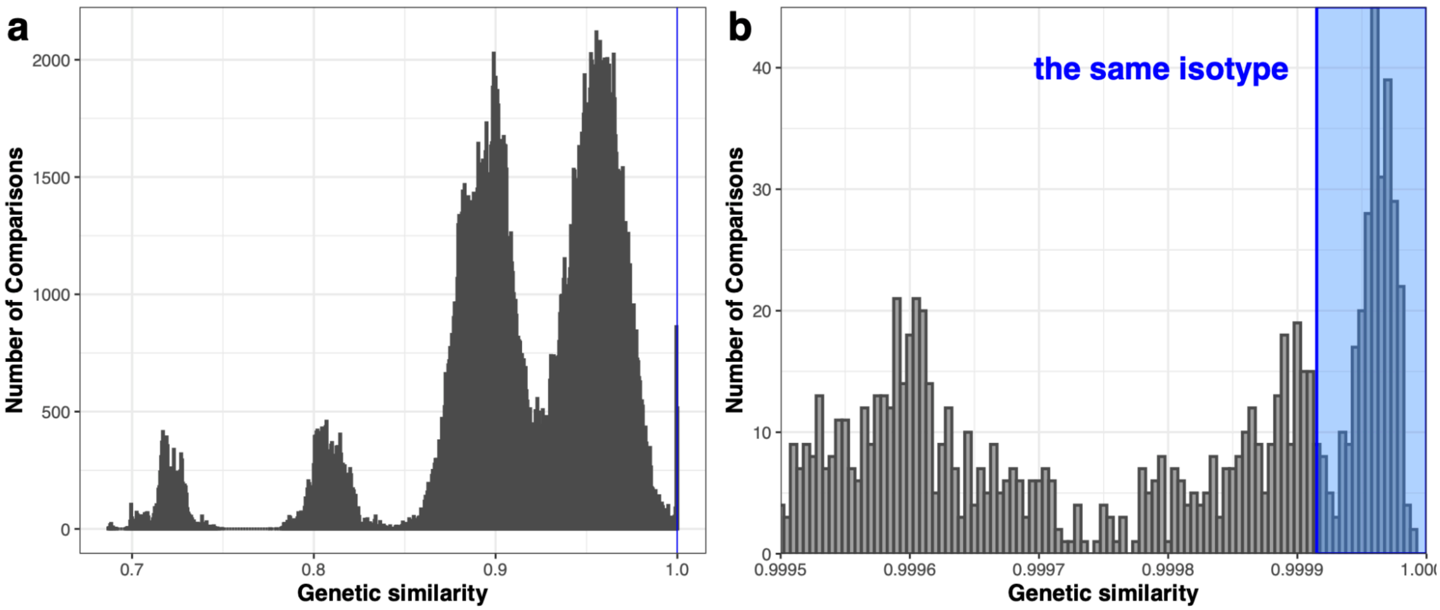


**Figure S37: Genetic similarity score distribution and isotype cutoff. a.** Histogram of genetic similarity for all pairwise comparisons. The vertical blue line indicates the threshold (99.9915%) used to define isotype identity. **b,** Zoomed-in histogram of the high similarity range (>99.9915%). The shaded blue area highlights comparisons classified as the same isotype.

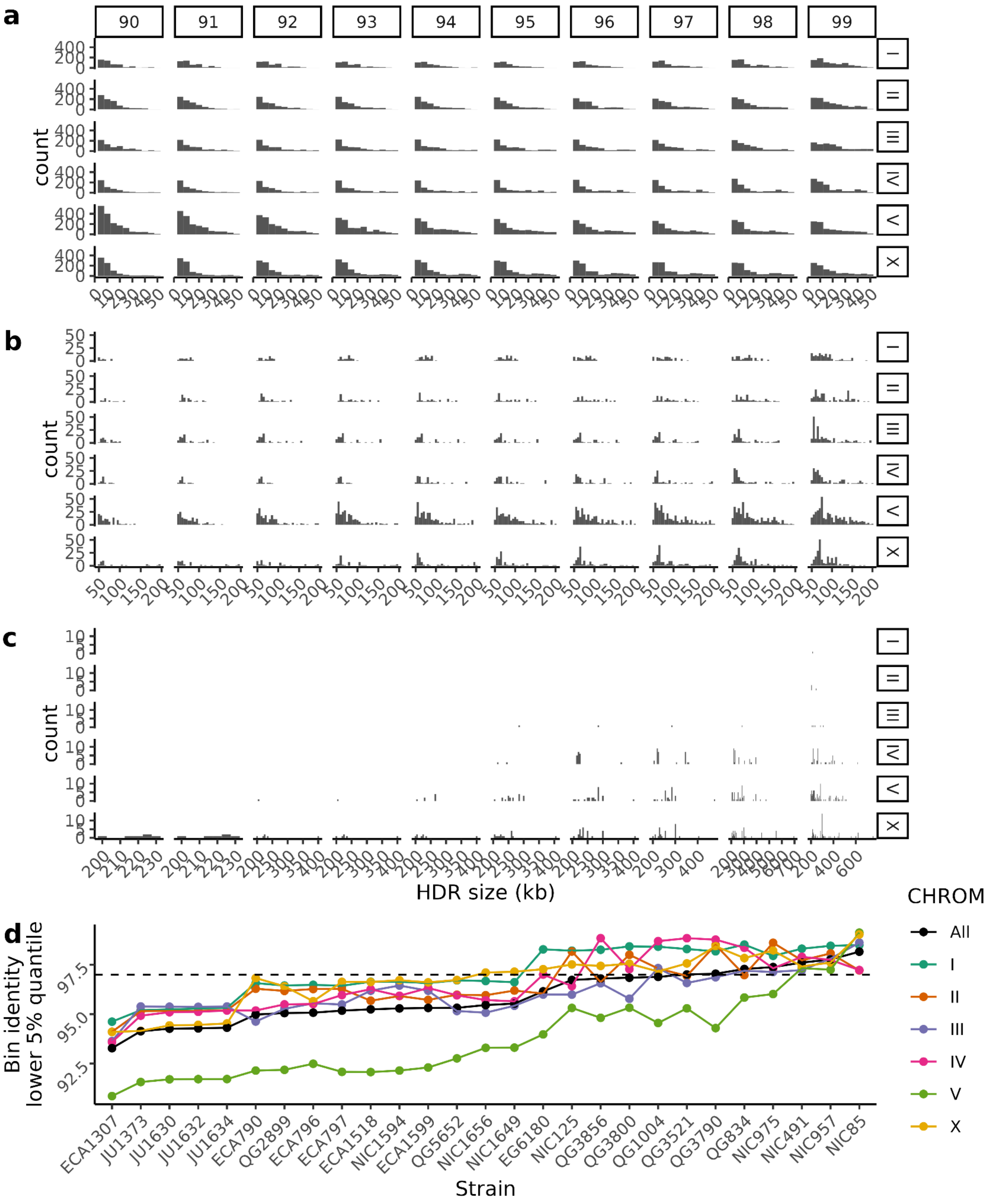


**Figure S38: Size and identity features of long-read based HDRs at variable identity thresholds.** Counts of HDRs across different identity thresholds (90-99%, represented on each vertical facet) across every chromosome (represented on each horizontal facet) for **a,** regions under 50 kb, **b,** regions over 50 kb and under 200 kb, and **c,** regions over 200 kb. **d,** for every isotype strain with a long-read genome (x-axis) across all chromosomes (black series) or each chromosome (each colored series), we estimated the lower 5% quantile of identity across all chromosome 1 kb bins (y-axis) and compared it against our selected identity threshold of 97% (dashed horizontal line).

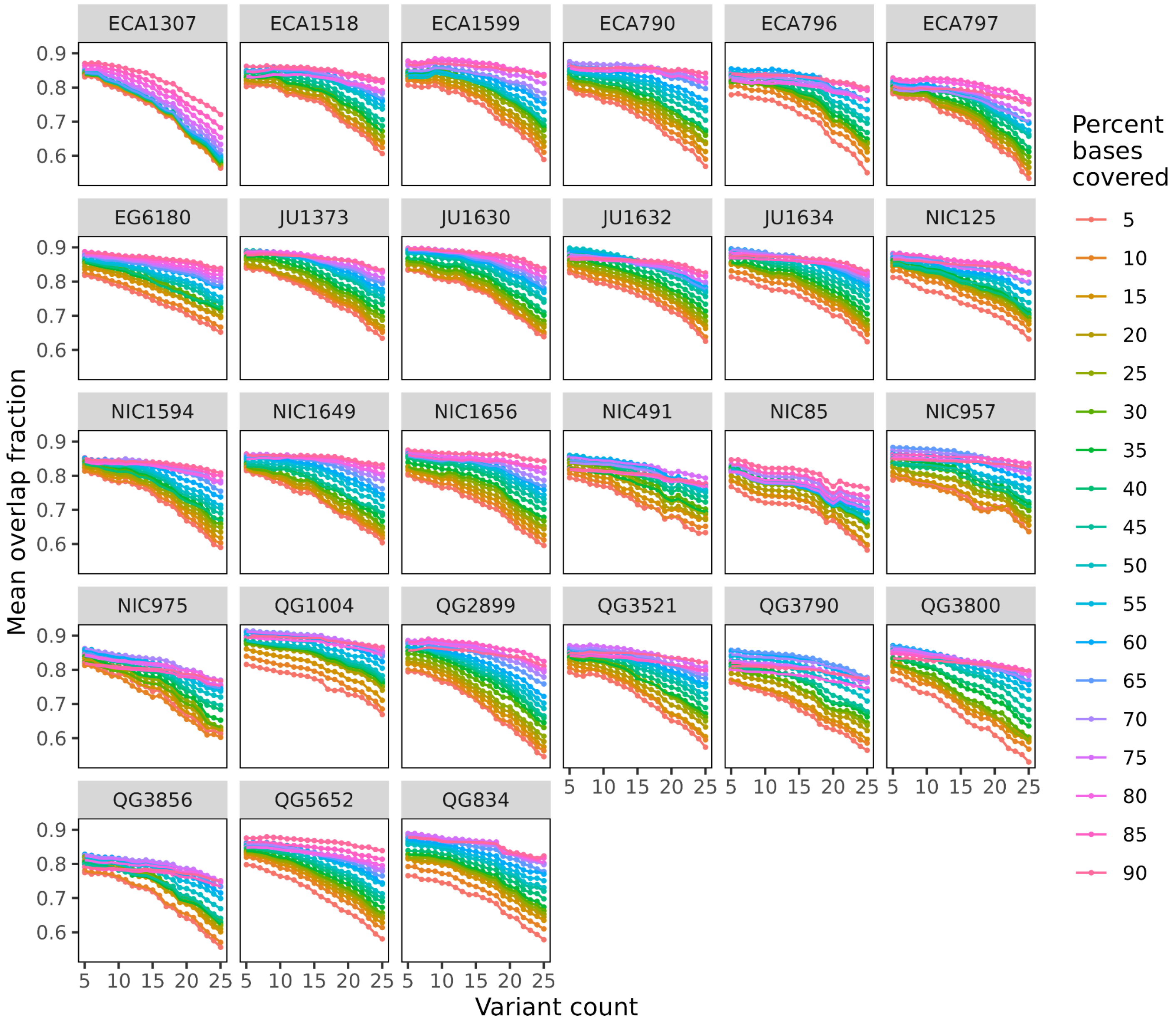


**Figure S39: Mean overlap fraction between short- and long-read HDR calls.** Estimates of HDR mean overlap fraction across 27 *C. tropicalis* strains for every pair of variant count and percent bases covered thresholds. Overlap fraction is estimated by quantifying the extent of the overlaps between short- and long-read based HDR calls, summarized into a mean overlap fraction estimate for each strain at each threshold pair. General trends indicate that mean overlap fraction improves at lower thresholds of variant count and higher thresholds of percent bases covered.

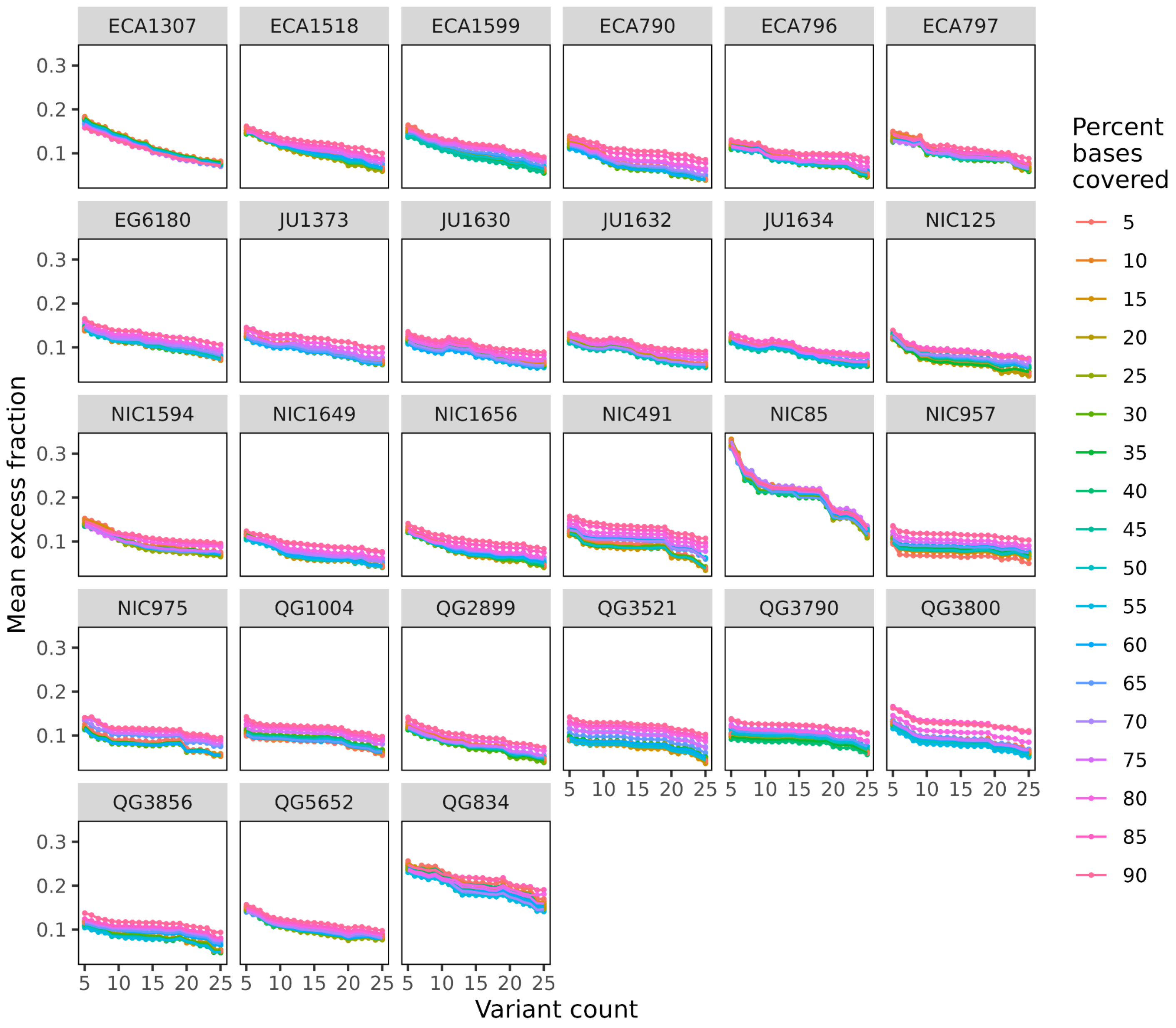


**Figure S40: Mean excess fraction between short- and long-read HDR calls.** Estimates of HDR mean excess fraction across 27 *C. tropicalis* strains for every pair of variant count and percent bases covered thresholds. Excess fraction is estimated by quantifying the extent of the short-read based hyper-divergent call that exceeds the boundaries of overlapping long-read HDR calls, summarized into a mean excess fraction estimate for each strain at each threshold pair. General trends indicate that mean excess fraction slightly improves at higher thresholds of variant count and lower thresholds of percent bases covered.

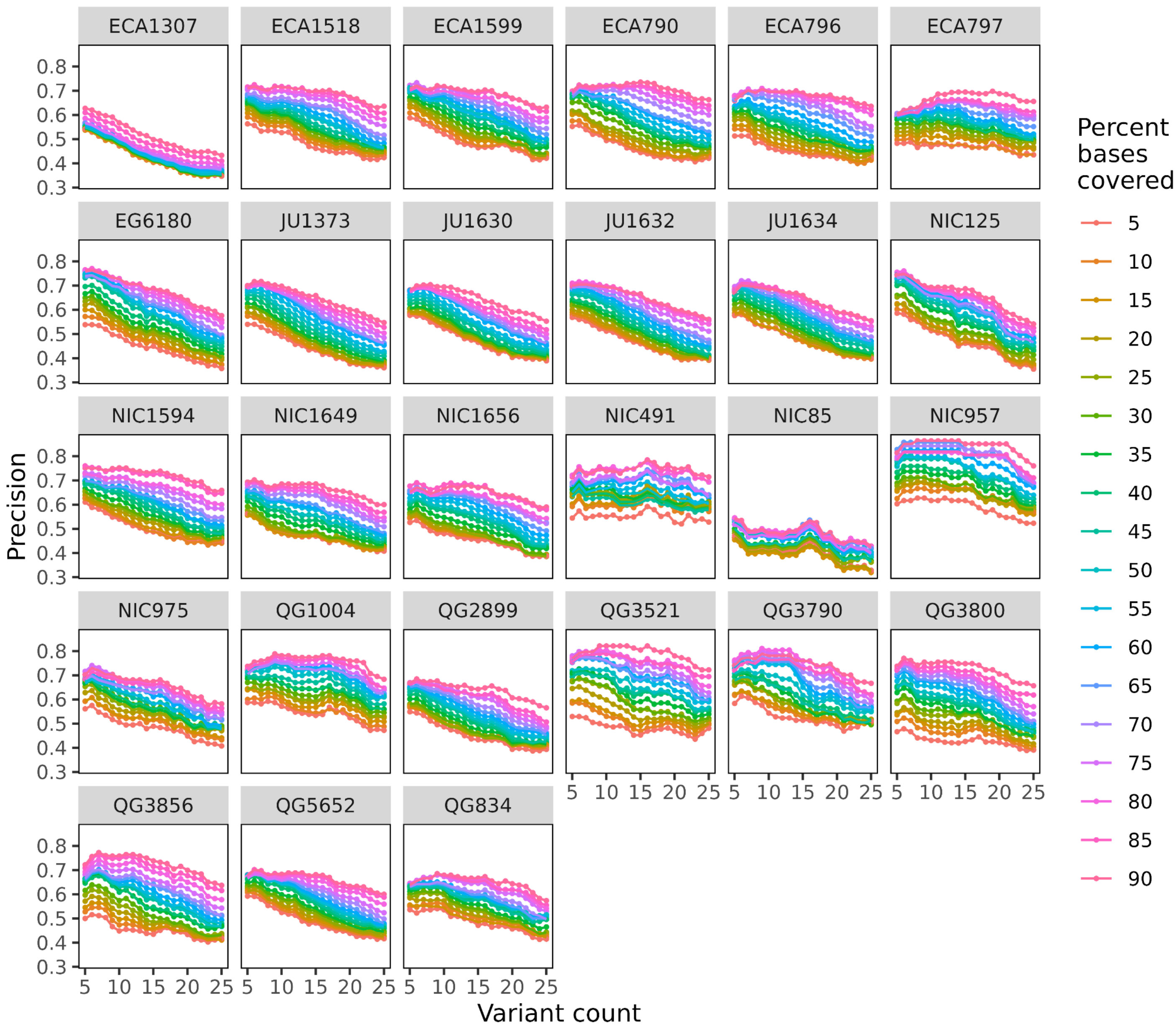


**Figure S41: Precision of short-read HDR calls.** Estimates of HDR precision across 27 *C. tropicalis* strains for every pair of variant count and percent bases covered thresholds. Precision is estimated by the proportion between the number of short-read based HDR calls that have an overlap with long-read based HDR calls relative to the total number of short-read based HDR calls. General trends indicate that precision improves at lower thresholds of variant count and higher thresholds of percent bases covered.

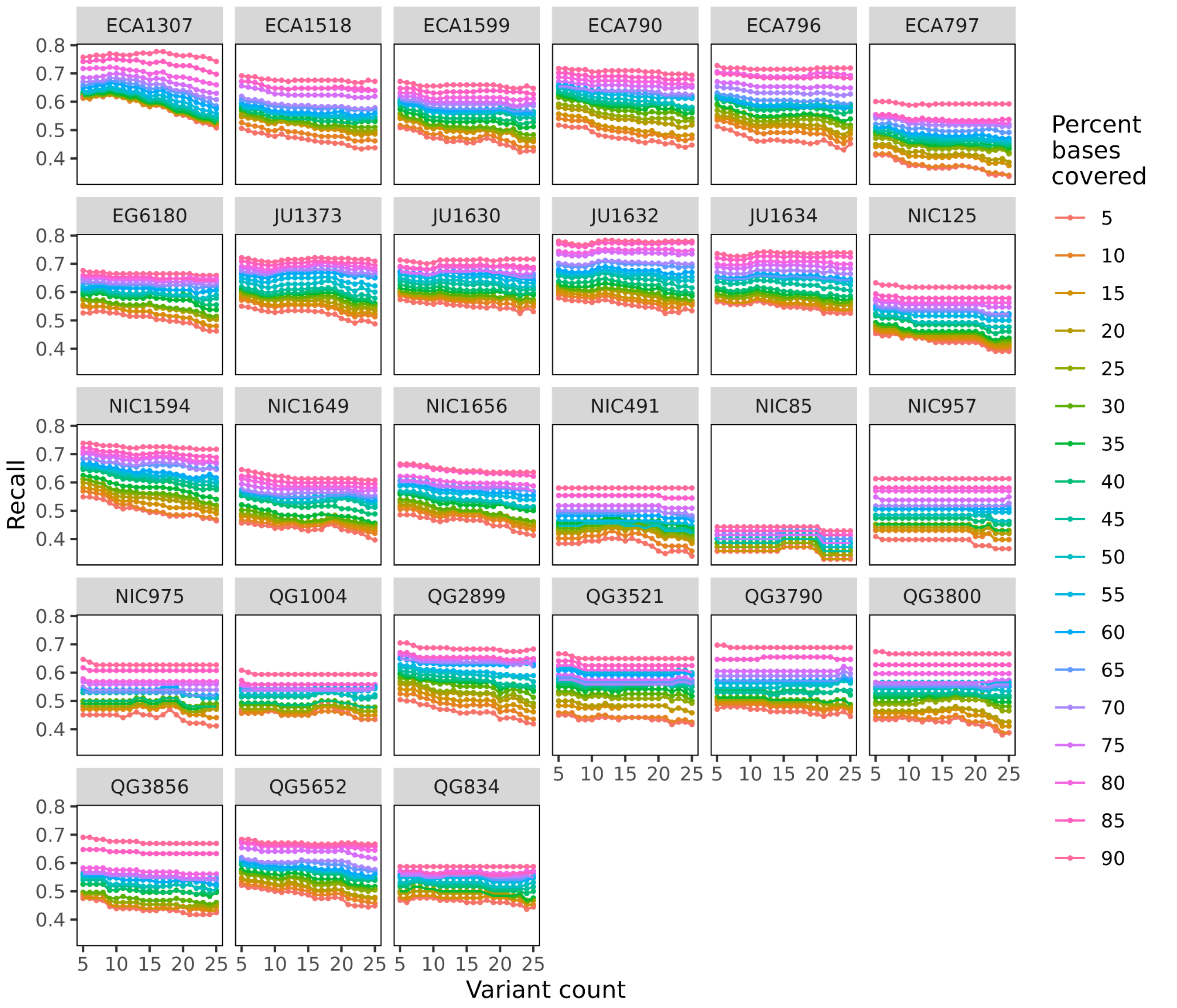


**Figure S42: Recall of long-read HDR calls.** Estimates of HDR recall across 27 *C. tropicalis* strains for every pair of variant count and percent bases covered thresholds. Recall is estimated by the proportion between the number of long-read based HDR calls that have an overlap with short-read based HDR calls relative to the total number of long-read based HDR calls. Similar to mean overlap fraction, general trends indicate that recall is nearly constant across thresholds of variant count with drastic improvement at higher thresholds of percent bases covered.

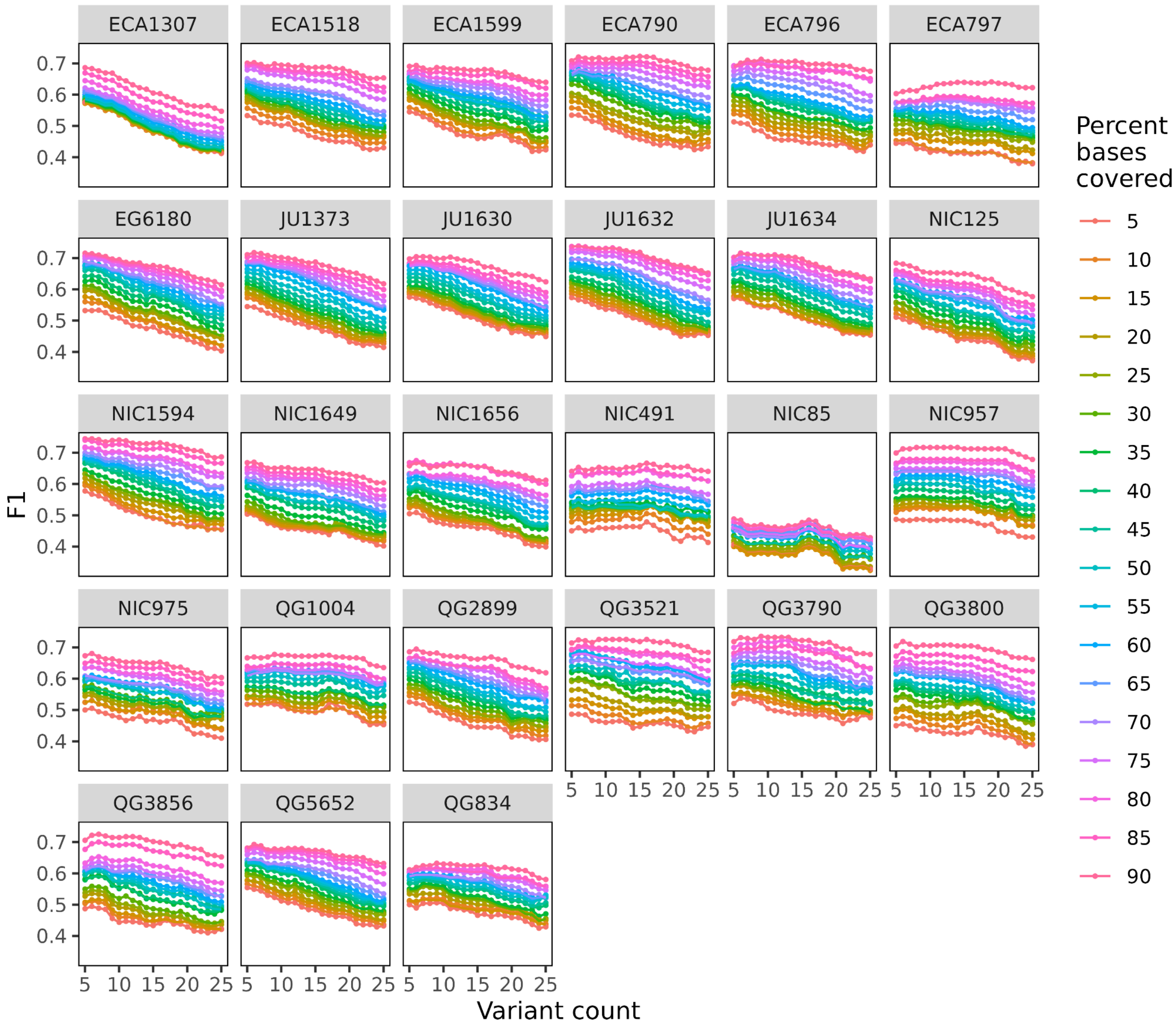


**Figure S43: F1 of short-read HDR calls.** Estimates of HDR F1 across 27 *C. tropicalis* strains for every pair of variant count and percent bases covered thresholds. F1 is estimated from the harmonic mean between recall and precision at each parameter pair.

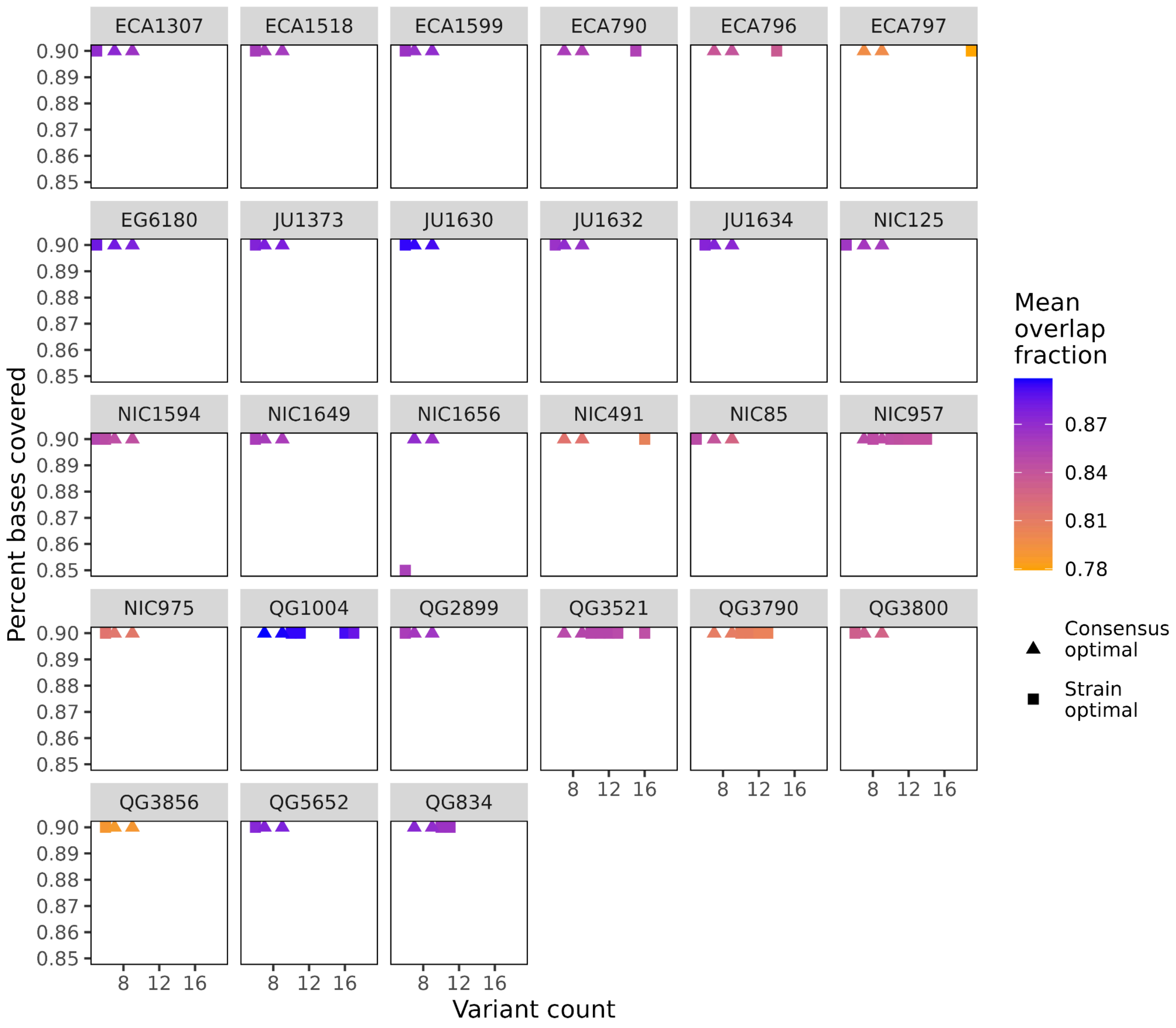


**Figure S44: Strain and consensus optimal threshold pairs for HDR calls.** A limited scatter plot showing the best threshold pair(s) optimized by F1 for each individual strain (displayed as squares, referred to as ‘strain optimal’). By selecting the top 5% (N=19) threshold pairs with the highest F1 in each strain, we identified two common threshold pairs across all 27 strains (displayed as triangles, referred to as ‘consensus optimal’). A gradient color scale describing the mean overlap fraction was applied to each threshold pair. The consensus optimal threshold pairs were a variant count of 7 or 9 SNVs with 90% bases covered. We selected 9 SNVs as the variant count threshold to call HDRs species-wide because of a minor improvement in mean overlap fraction.
